# Anatomy of the mandibular symphysis of extant cercopithecids: taxonomy and variation

**DOI:** 10.1101/2024.03.31.587451

**Authors:** Laurent Pallas, Masato Nakatsukasa, Yutaka Kunimatsu

**Affiliations:** Kyoto University, Graduate School of Science, Laboratory of Physical Anthropology, Kyoto 606-8502, Japan; Laboratoire de Paléontologie, Évolution, Paléoécosystèmes et Paléoprimatologie (PALEVOPRIM), UMR 7262, Université de Poitiers, Poitiers Cedex 86022, France; Ryukoku University, Faculty of Business Administration, Fushimi, Kyoto 612-8577, Japan

**Keywords:** Morphometry, colobines, cercopithecines, diet, evolution

## Abstract

The symphyseal anatomy of extant and fossil cercopithecids has not yet been demonstrated as a useful tool for taxonomic discrimination, and the source of variation in cercopithecid symphysis has not been addressed on a broad taxonomic scale. Here, we used linear and angular dimensions to quantify symphysis shape. Using univariate, multivariate data and allometric regressions (partial least squares and phylogenetic generalized least square regressions), we addressed the hypothesis that extant cercopithecids can be distinguished by symphysis shape. Significant differences in univariate and multivariate data and allometric regressions permitted to distinguish cercopithecids at the subfamilial, tribal, and genus levels. We showed that multivariate data followed the distribution expected under Brownian Motion and significantly discriminates taxa at different taxonomic levels. Colobine symphysis are characterized by developed inferior transverse tori, short planum alveolare, and short symphysis, whereas cercopithecine symphysis are characterized by developed superior transverse tori, long planum alveolare, and long symphysis. Exceptions to this pattern exist within each subfamily, and this study underlines the particular anatomy of *Colobus* and *Presbytis* among the colobines, *Allenopithecus* among the Cercopithecini, and *Theropithecus* and *Lophocebus* among the Papionini. We also demonstrate that the relative development of the transverse tori, the relative length of the planum alveolare and symphyseal inclination are dimorphic traits. Specifically, we show that the symphysis of *Procolobus verus*, *Nasalis larvatus*, and *Papio anubis* is strongly dimorphic.

## INTRODUCTION

The family Cercopithecidae (Gray, 1821) includes 22 extant genera (Grubb et al., 2003; Brandon-Jones et al., 2004; Groves, 2007), distributed in two subfamilies: Cercopithecinae (Gray, 1821) and Colobinae (Jerdon, 1867). A diversity of ecological preferences in locomotion and diet is observed among and between both subfamilies. Colobines are generally considered as stenotopic arboreal primates mostly restricted to closed and forested environments (Kingdon and Groves, 2013a). A greater taxonomic diversity is observed in Asian colobines (Presbytini) relative to African colobines (Colobini) (Brandon-Jones et al., 2004; Groves, 2007). Cercopithecines, which include the speciose Cercopithecini and the widely distributed Papionini, exhibit a broader, eurytopic ecological niche, as illustrated by the mixed locomotor substrate preferences and omnivorous diet of most of their representatives (Grubb et al., 2003; Kingdon and Groves, 2013b; Lo Bianco et al., 2017).

The anatomy of the mandibular symphysis of extant and fossil cercopithecids has long been recognized as taxonomically informative (Freedman, 1957; Leakey, 1982; Benefit and Pickford, 1986; De Bonis et al., 1990; Frost and Delson, 2002; Hlusko, 2006, 2007; Groves, 2007; Benefit, 2008; McKee et al., 2011; Pallas et al., 2019). The inclination of the symphyseal profile relative to the alveolar plane, the overall robustness of the symphysis, and the relative and absolute development of the tori are among the mandibular characters used to identify extant and fossil cercopithecids (Figure 1). Yet, these diagnoses often rely on qualitative assessment of the symphyseal morphology (e.g., Frost, 2001; Groves, 2007) or on crude quantitative estimation of its shape, as seen, for example, in the assessment of the symphysis length relative to molar length in Koufos et al. (2003), symphysis cross-sectional area calculated from symphysis height and breadth in Wood (1976), or symphysis length divided by its breadth in Arenson et al. (2022). In addition to diagnosis of fossil and extant cercopithecids, many data have been provided by ecomorphological and biomechanical studies (Hylander, 1984, 1985; Ravosa, 1996; Jablonski et al., 1998; Vinyard and Ravosa, 1998; Daegling and McGraw, 2001, 2007; Panagiotopoulou and Cobb, 2011) of the cercopithecid symphysis, sometimes with taxonomically rich datasets (Ravosa, 1996; Jablonski et al., 1998), or sophisticated quantification of symphyseal shape (Panagiotopoulou and Cobb, 2011). The protocol presented by Pallas et al. (2019), and repeated by Locke et al. (2020), is so far, the most complete given its quantification of tori breadth and planum alveolare length, among others. However, the computation of the symphyseal morphometric ratios of Pallas et al. (2019) was not followed by statistical analyses to firmly demonstrate significant differences between taxa, in addition to being focused on colobines only. Furthermore, no assessment of the effect of sexual dimorphism and/or allometry was given in Pallas et al. (2019). Although the source of variation behind the cercopithecid symphyseal morphology has already been assessed in light of allometry (Hylander, 1985; Vinyard and Ravosa, 1998) and sexual dimorphism (Wood, 1976; Jablonski et al., 1998; Fukase, 2011), these analyses were based on a limited set of taxa. Therefore, the level of variation of transverse tori breadth, planum alveolare length and symphyseal inclination is largely unknown in cercopithecid and remains to be assessed before accepting their efficacy in taxonomic distinctions.

Sexual dimorphism in body mass is variable in cercopithecids, with *Mandrillus* and *Nasalis* among the most sexually dimorphic cercopithecines and colobines, respectively (Leutenegger, 1982). On the other hand, the small colobine *Presbytis* is monomorphic in body mass (Leutenegger, 1982). Allometry has been shown to be a pervasive factor in taxonomic identification (Collard and Wood, 2001; Gilbert et al., 2009), with authors emphasizing the use of dimorphic traits to discriminate taxa (Gilbert, 2013). It has also been studied as a major source of variation given the positive allometry of mandibular length with body size (Hylander, 1984). Symphysis breadth was subsequently demonstrated as positively allometric in papionins (Hylander, 1985; Vinyard and Ravosa, 1998). Sexual dimorphism in the cercopithecid mandible is also observed in the shape and proportions of the dentition, and more precisely in the development of the C/P_3_ complex (Plavcan and Van Schaik, 1992). The upper and lower canines of male cercopithecids differ in dimensions from those of the female, particularly in height (Plavcan, 1993). However, the impact of sexual dimorphism of the C/P_3_ complex and canine dimensions on the shape of the symphysis is rarely discussed and is based only a limited number of taxa (e.g., *Macaca* and *Papio*) in Fukase (2011).

Here, we test the hypothesis that extant cercopithecids can be phenetically distinguished at the subfamilial, tribal, and genus levels using multivariate and univariate analyses derived from four linear measurements (symphyseal length (SL), breadth of the superior transverse torus (STT), breadth of the inferior transverse torus (ITT), and length of the planum alveolare (PAL)) and one angular measurement (symphyseal inclination). Univariate data, in the form of morphometric ratio, have been shown to be useful taxonomic discriminants (Pallas et al., 2019) and have the advantage of not relying on complete morphometric datasets, thus leaving the possibility to include damaged fossil specimens in future analyses. We computed three ratios, relative breadth of the transverse tori (STT/ITT) and two ratio of relative length of the planum alveolare (PAL/Geometric Mean*100 and PAL/SL*100), and assessed their efficiency in taxonomic discrimination. Significantly distinct phenetic differences are identified using statistical tests for univariate data (ratio) and principal component scores. We also assess, using comparative phylogenetic methods, whether a phylogenetic signal can be reliably inferred using our morphometric dataset. After assessing for shape differences, we evaluate whether sexual dimorphism significantly influences the shape of the symphysis between taxa for which we have enough specimens to properly sample variation. We also test the effect of allometry on symphyseal shape using geometric mean as the predictive variable.

## MATERIAL AND METHODS

### NEONTOLOGICAL SAMPLE

We collected data on 756 individuals from 88 species of 22 cercopithecid genera : *Allenopithecus* (n = 15), *Allochrocebus* (n = 4), *Cercocebus* (n = 14), *Cercopithecus* (n = 56), *Chlorocebus* (n = 20), *Colobus* (n = 99), *Erythrocebus* (n = 10), *Lophocebus* (n = 23), *Macaca* (n = 169), *Mandrillus* (n = 7), *Miopithecus* (n = 7), *Nasalis* (n = 48), *Papio* (n = 62), *Piliocolobus* (n = 41), *Presbytis* (n = 70), *Procolobus* (n = 23), *Pygathrix* (n = 5), *Rhinopithecus* (n = 2), *Semnopithecus* (n = 14), *Simias* (n = 25), *Theropithecus* (n = 15), *Trachypithecus* (n = 27). Information on taxonomy, accession numbers and sexes of specimens is provided in SOM Table S3.

### MORPHOMETRIC PROTOCOL

We acquired digital photographs of the cross-section of the symphysis on 3D models generated using a surface scanner (Einscan Pro 2X) used by L.P and generated 3D data from computed tomography scans collected from different sources. Sources of the data, for each specimen, can be found in SOM Table S3. Photographs of cross-sections set at the symphyseal midline were acquired using Avizo v.7.0 (Thermo Fisher Scientific, Waltham). First, the alveolar process of the mandible was aligned on the transverse plane using the trackball function of Avizo. Second, the anterior aspect of the symphysis was visualized by moving the virtual model using the software trackball function, and a cross-section, set in the sagittal plane and passing through the infradentale, was then obtained (Figure 1).

Measurements were taken using ImageJ v.1.50e (Schneider et al., 2012), with all data taken by L.P. The measurement protocol follows that presented in Pallas et al. (2019) but the length of the planum alveolare was modified. Instead of measuring the projection of the planum alveolare following the orientation of the transverse plane, planum alveolare was measured as the distance from the infradentale to the most posterior point of the STT. Illustrations of the measurement protocol and cross-sectional acquisition are shown in Figure 1. The formulae of the indices are as follows for the relative development of the transverse torus and planum alveolare length: (STT/ITT)*100, (PAL/Geometric Mean)*100, and (PAL/SL)*100. For the relative length of the planum alveolare, the geometric mean (GM) was calculated from the SL, STT and ITT measurements (see Figure 1 for abbreviations). We included two ratios for the assessment of the planum alveolare length as the PAL/SL ratio permit to includes fossil specimens that preserves both variables while PAL/GM requires to have access to symphysis also preserving the transverse tori. Both indices convey a similar signal as indicated by a high correlation (0.8) given Pearson’s correlation test and by a linear regression of PAL/SL on PAL/GM (SOM Figure S1) which gives a significant relationship between both variables.

**Figure 1:**
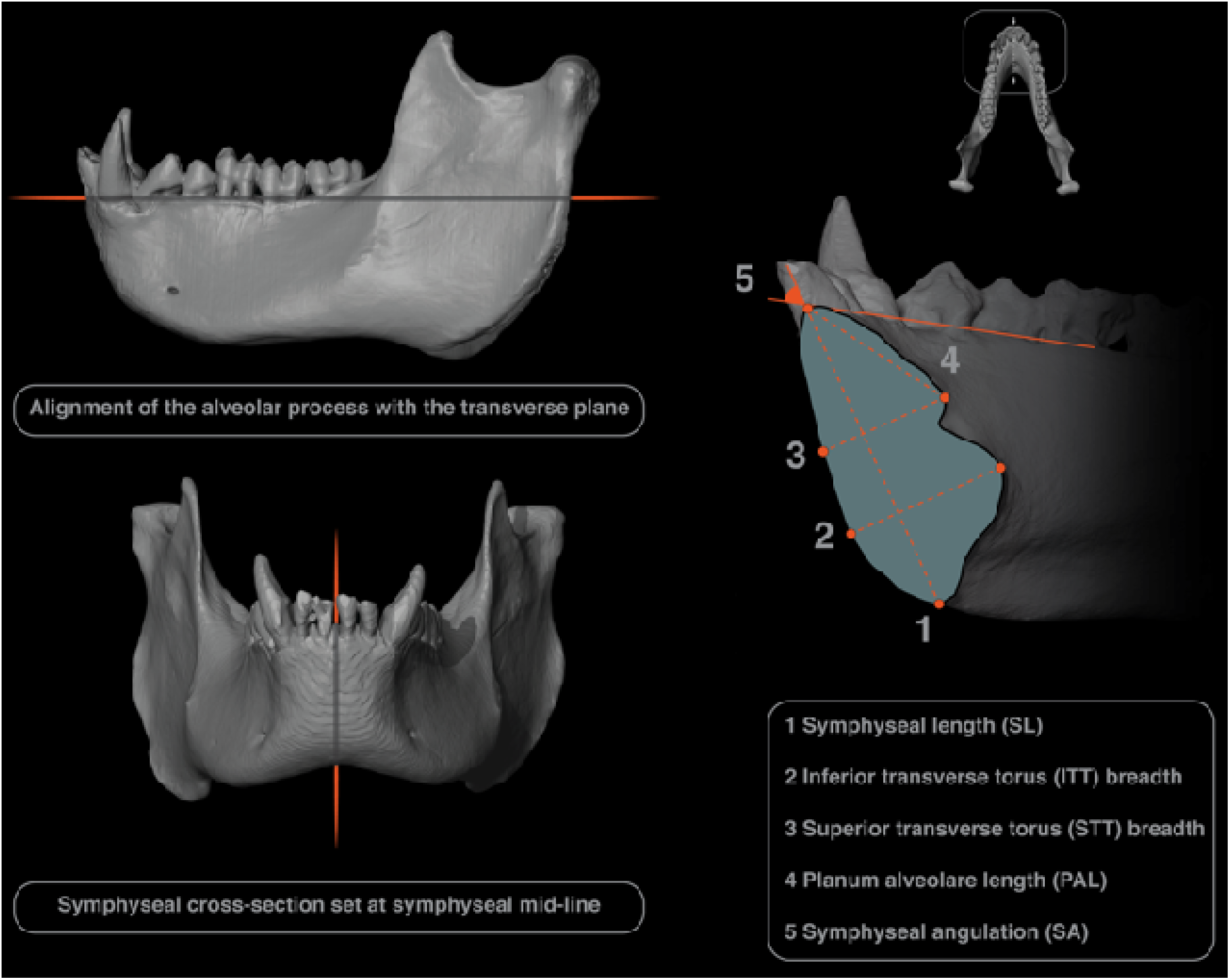
Illustration of the acquisition protocol of symphyseal cross-section and symphyseal measurements.

### STATISTICAL ANALYSES

All statistical analyses were performed using R v. 3.5.0 (R Core Team, 2021). Before testing for the significance of each statistical model, we tested the normal distribution and homogeneity variances (i.e., homoscedasticity) of the model residuals using Shapiro-Wilk and Bartlett’s tests, respectively (SOM Tables S4 to S6). Non-parametrical tests were used in cases where homoscedasticiy was not verified. The significance level for these tests were set at 5% (marginal differences), 1% (significant differences), and 0.1% (highly significant differences). Given that the ratio of planum alveolare length divided by symphyseal length gave comparable results in terms of taxonomic discrimination to the ratio of planum alveolare length divided by GM (SOM Figure S2), we focused our statistical analysis on the latter ratio. For comparative purposes, and especially fossil analyses, we have provided raw data regarding the first mentioned planum alveolare ratio (SOM Tables S1 and S2).

#### Significant differences in multivariate and univariate data

In the case of normally distributed and homoscedastic residuals, we used Analysis of Variance (ANOVA), combined with Tukey’s Honest Significant Difference (HSD) post-hoc test, to identify significantly distinct pairs of taxa. In case of non-homoscedastic residuals, we used a Kruskall-Wallis test, combined with a Dunn’s post-hoc test. The parameters of the models of ANOVA’s and Tukey’s HSD tests were calculated with the aov() and TukeyHSD() functions of the ‘stats’ package (R Core Team, 2021). The parameters of the models of Kruskall-Wallis’s and Dunn’s tests were calculated with the kruskal.test() and dunnTest() functions of the ‘stats’ and ‘FSA’ packages, respectively (Ogle, 2018, R Core Team, 2021).

In addition to looking for shape differences between taxa, we are evaluating differences in morphometric ratio values between male and female specimens of selected cercopithecid taxa for which the sample size is judged adequate. Here, we adopt the criteria of Gilbert and Grine (2009). Indeed, based on the morphometry of the papionin maxilla, they demonstrated that a sample size of approximately 15 specimens is reasonable for estimating the population mean. Accordingly, we evaluate the effect of sexual dimorphism on symphysis shape in the following taxa: *Colobus polykomos*, *Colobus guereza*, *Piliocolobus badius*, *Procolobus verus*, *Presbytis bicolor*, *Nasalis larvatus*, *Cercopithecus mitis*, *Chlorocebus aethiops, Macaca fascicularis*, *Macaca fuscata*, and *Papio anubis*. Some of these taxa are known to be highly dimorphic in body mass and canine height (e.g., *Papio anubis*), while others are monomorphic in body mass and show relatively reduced dimorphism in canine height (e.g., *Presbytis bicolor*) (Plavcan and Van Schaik, 1992). We expect taxa with a strongly dimorphic C/P_3_ complex to exhibit dimorphism in symphyseal shape, consistent with the work of Fukase (2011).

#### Comparative phylogenetic analyses

According to the Brownian Motion (BM) model of anatomical traits evolution, the independence of morphometric data among a set of taxa is proportional to their divergence time (assuming generation times are more or less constant), meaning that closely related species are expected to be much more similar than distantly related species (Pagel and Harvey, 1988; Münkemüller et al., 2012). Here we evaluate the effect of phylogenetic relationships on symphyseal shape using Pagel’s λ computed with the phylosig() function of the phytools package (Revell, 2012). Values of λ close to zero indicate that the morphometric data are not distributed under BM while values close to one point to the expected distribution under BM. Branch lengths, divergence dates and topology of the tree used in the comparative phylogenetic analysis (Figure 2) is derived from Perelman et al. (2011), with additional data regarding branch lengths and divergences dates from Ting (2008), and Liedigk et al. (2012). Precisely, the topology of the tree follows Perelman et al. (2011) with the addition of *Simias concolor* as a sister taxon to *Nasalis larvatus* following Whittaker et al. (2006). Divergence dates follow Perelman et al. (2011) with the exception of Ting (2008) for the divergence of *Colobus* from *Piliocolobus*/*Procolobus*, and Liedigk et al. (2012) for the divergence date of *Simias* and *Nasalis*. The phylomorphospace shown in Figure 3 was obtained using the R script presented in Barr (2020).

#### Allometry

We used both phylogenetic least squares regressions (PGLS) and partial least squares (PLS) regressions to study allometry. For PGLS, we used the pgls() function of the ‘caper’ package and the lm() function of the ‘stats’ package for PLS. Allometry was modeled as the natural logarithm of the morphometric variablesresponse variable and the natural logarithm of the geometric mean as the predictor variable. For each regression, the geometric mean was calculated using all morphometric variables. To justify the use of the geometric mean as a proxy for body mass, we regressed the average geometric mean of each taxon on the average body mass of male and female specimens. Detailed information regarding data used for body mass can be found in SOM Table S7.

**Figure 2:**
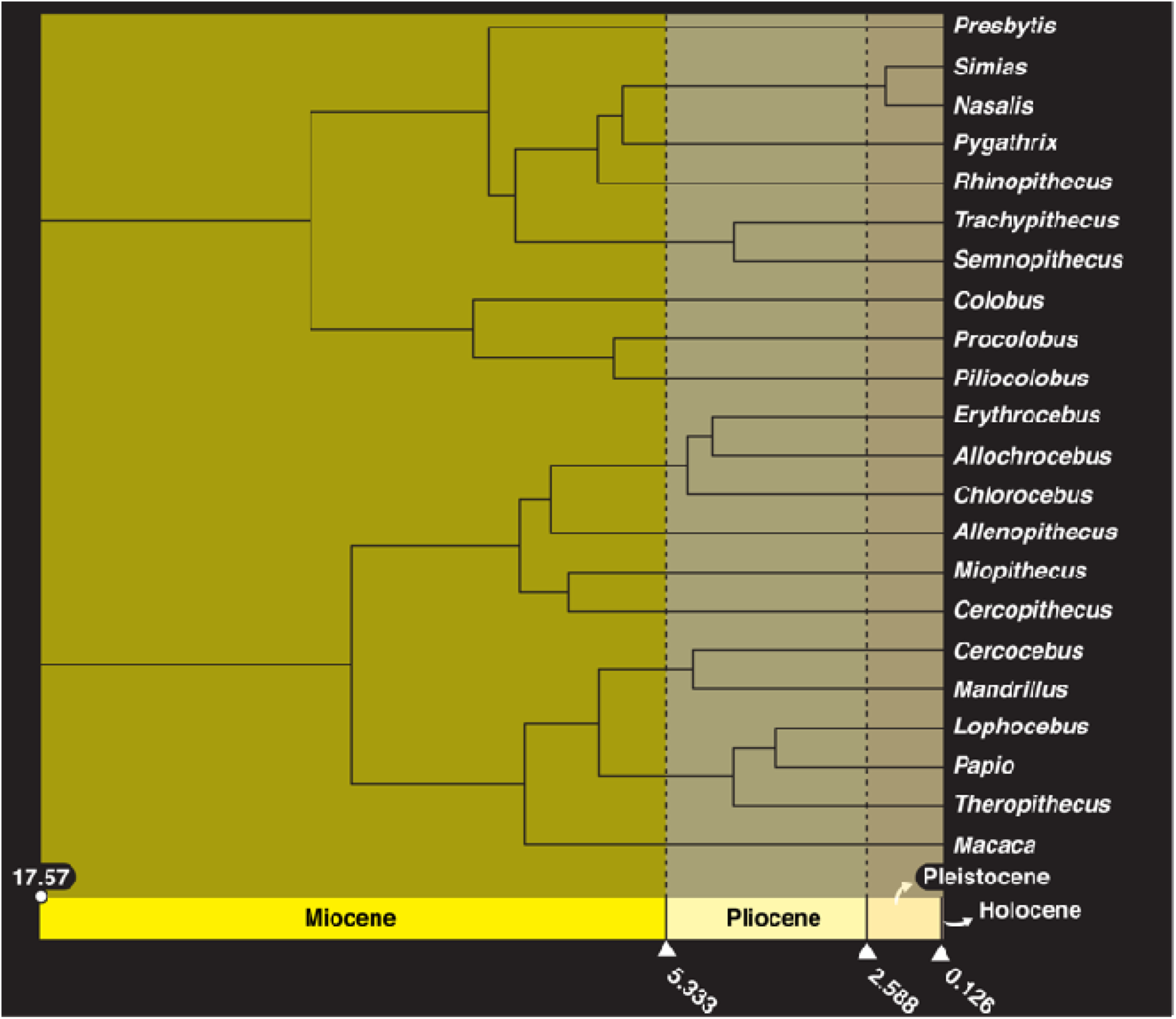
Chronophylogram of the extant cercopithecid taxa considered in this study, with topology and divergence dates following Ting (2008), Perelman et al. (2011), and Liedigk et al. (2012).

## RESULTS

### PCA SCORES AND OVERALL MORPHOMETRIC DIFFERENCES

PC1 accounts for 62.55% of the variance (Figure 3A) and positive scores on this axis are mainly due to symphyseal length, planum alveolare length and breadth of the STT (Figure 3B). The colobines show negative scores on PC1, with the exception of *Colobus* (Figure 3A). The low scores of colobines reflect their relatively short planum alveolare, small STT and short symphysis (Figure 3B). Positive PC1 scores are characteristic of Papionini, with the exclusion of *Lophocebus*. It illustrates their long planum alveolare, large STT and elongated symphysis (Figure 3B). The distribution of Cercopithecini scores on PC1 is less straightforward to interpret from a taxonomic point of view compared to colobines and papionins. Indeed, while *Cercopithecus* and *Erythrocebus* show positive scores, *Allenopithecus*, *Chlorocebus*, *Allochrocebus* and *Miopithecus* present negative scores (Figure 3A and B).

PC2 explains 22.95% of the variance (Figure 3A), with positive scores influenced by symphyseal inclination and negative ones by ITT breadth (Figure 3B). Most colobines, with the exceptions of *Procolobus*, *Presbytis*, and *Colobus*, have negative scores on PC2. This reflects their relatively large ITT and steep symphysis (Figure 3B). Papionins present either have very negative scores on PC2, as seen in *Theropithecus*, *Papio* and *Mandrillus*, or scores that cluster around zero in *Lophocebus*, *Cercocebus* and *Macaca*. While the distribution of Cercopithecini was equivocal on PC1, it is more clearly interpretable on PC2, with positive scores for all taxa except *Allenopithecus* (Figure 3A and B).

In summary, colobines (except *Colobus*) generally exhibit negative scores on PC1 and PC2. The large papionins *Theropithecus*, *Papio* and *Mandrillus* all show positive scores on PC1 and negative on PC2. *Lophocebus*, *Cercocebus* and *Macaca* can be distinguished from the large papionins by presenting positive scores on PC2. *Allenopithecus* can be differentiated from all other guenons by showing negative scores on PC1 and PC2.

**Figure 3:**
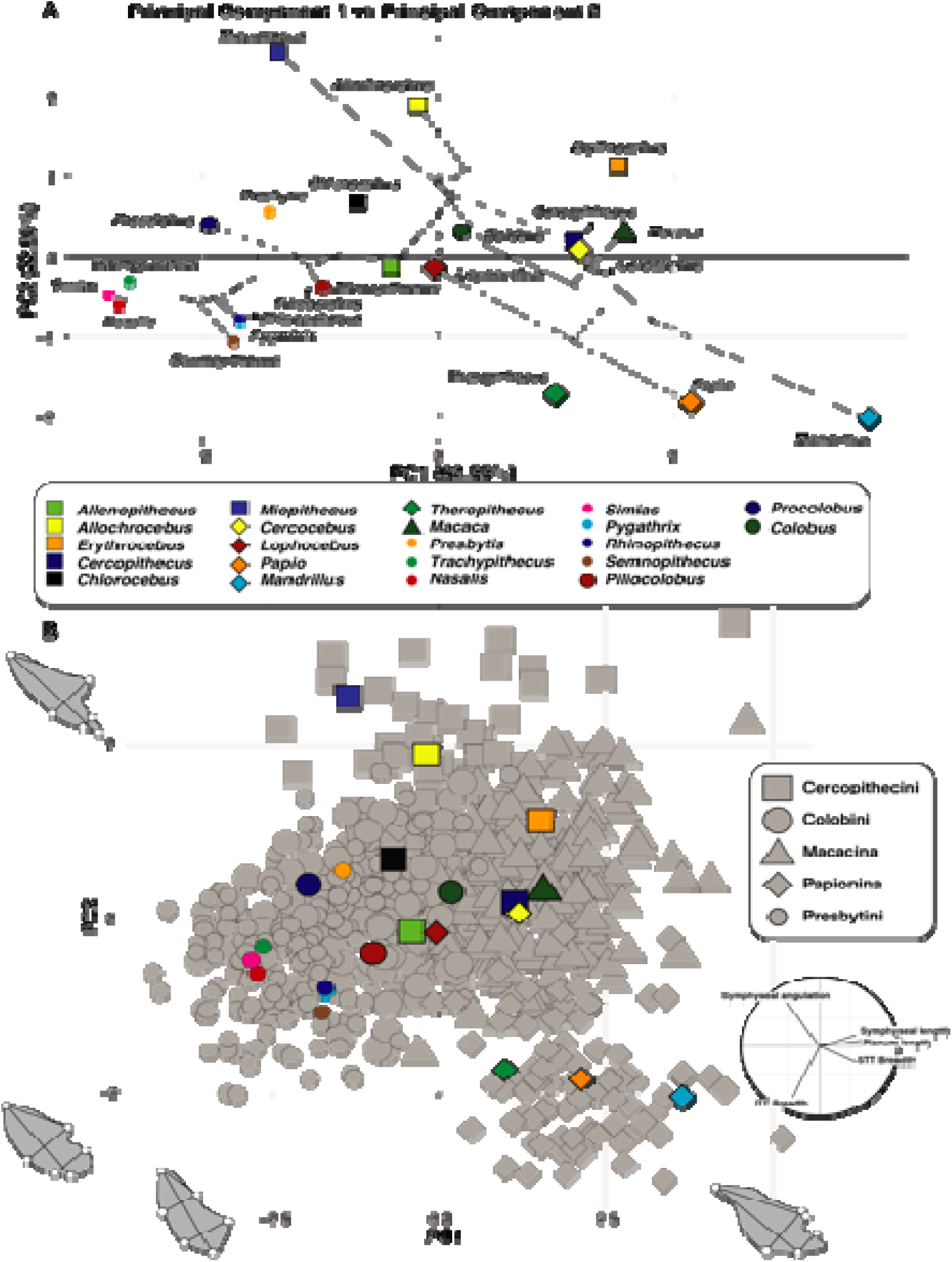
A) Phylomorphospace of PC1 and PC2, and B) biplot of PC1 and PC2. An illustration of the symphyseal morphology of the specimens that present maximum scores along PC1 and PC2 scores is provided in biplot B).

### MORPHOMETRIC RATIO AND OVERALL MORPHOMETRIC DIFFERENCES

The symphysis of the odd-nosed monkeys *Simias*, *Nasalis* and *Rhinopithecus* is more obtuse than that of other cercopithecids, especially its sister taxa, the African colobines (Figure 4A and Table 1). Among African colobines, *Piliocolobus* present an acute-angled symphysis relative to *Colobus* and *Procolobus* (Figure 4A and Table 2). An extensive overlap in the range of variation in symphyseal inclination is seen in guenons. Among papionins, *Theropithecus* exhibits an obtuse-angled symphysis, with values contrasting with the acute-angled ones in *Mandrillus* and *Macaca*. *Papio*, *Cercocebus*, and *Lophocebus* have symphyseal inclination values intermediate between those of *Theropithecus* and *Mandrillus* (Figure 4A and Table 2).

**Table 1:**
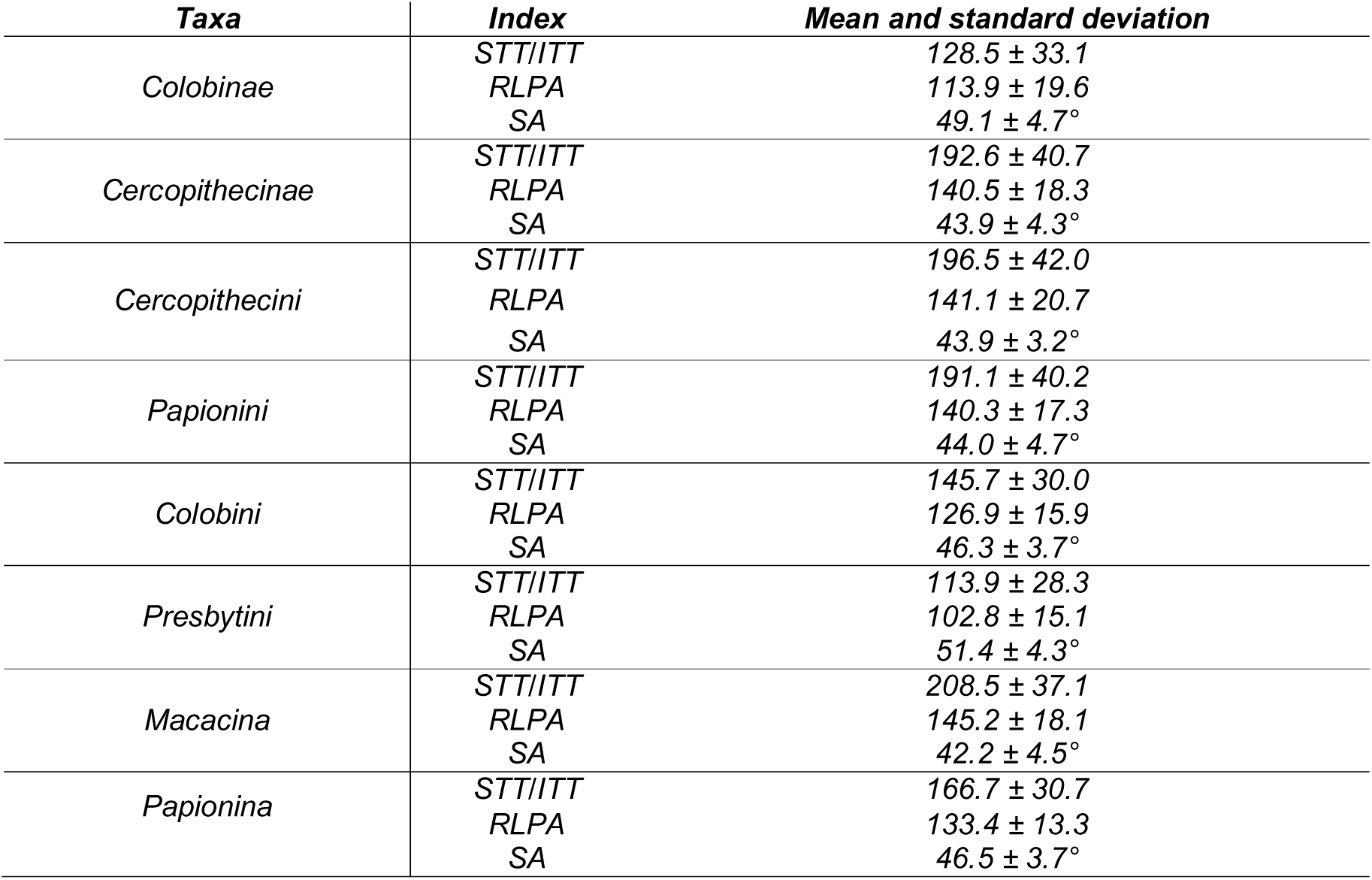
Descriptive statistics (mean and standard deviation) of the morphometric indexes (STT/ITT and RLPA) per subfamily, tribe, and subtribe.

**Table 2:**
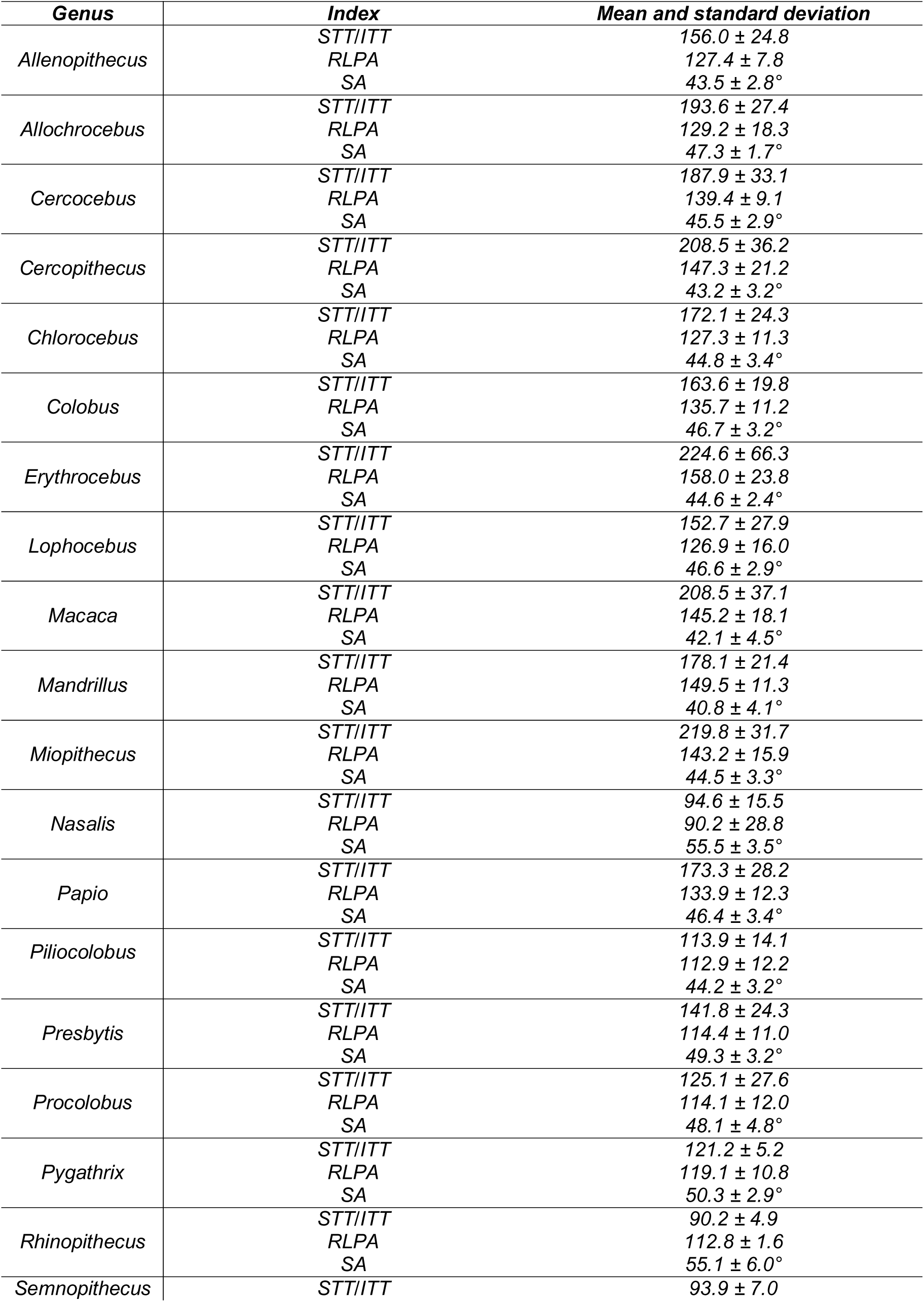
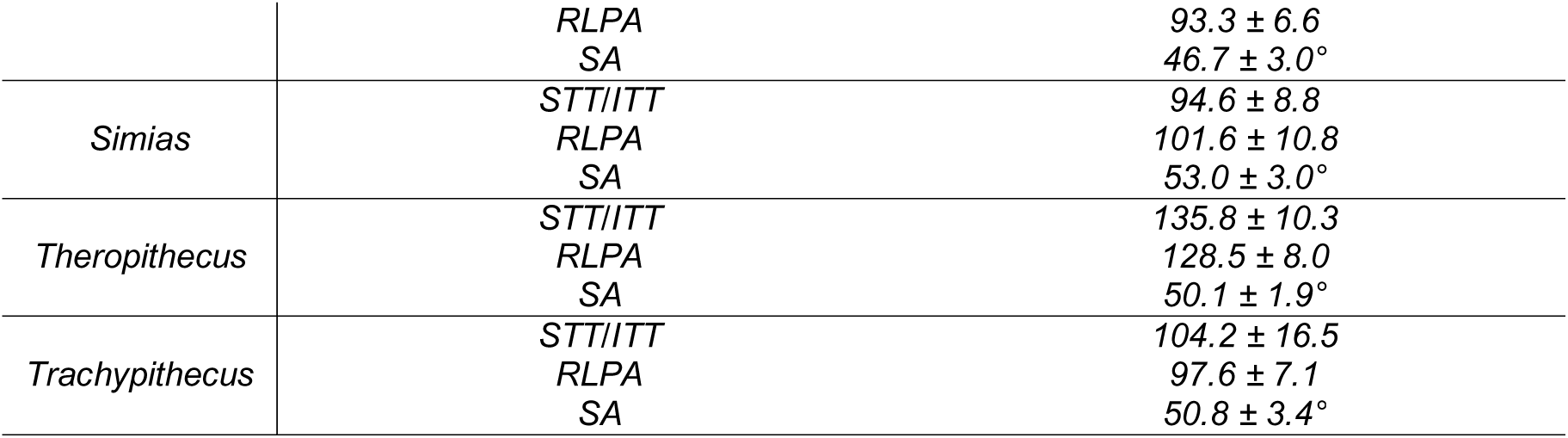
Descriptive statistics (mean and standard deviation) of the morphometric indexes (STT/ITT and RLPA) and symphyseal angulation (SA in °) per genera.

Colobines are characterized by low scores in relative breadth of the transverse tori (Figure 4B and Table 1), illustrating their relatively large STT. *Presbytis* and *Colobus* are exceptions to this pattern by showing high scores and relatively small ITT compared to all other colobines. *Allenopithecus* differs from all other Cercopithecini by presenting a relatively more developed ITT (Figure 4B and Table 2). A similar pattern is seen in *Theropithecus*, which differs from other extant papionins in having low scores, and thus a more developed ITT.

The relative length of the planum alveolare is heterogeneous within each cercopithecid tribe and subtribes (Figure 4C and Table 1). Whereas a relatively long planum is seen in *Presbytis*, a short planum is observed in *Semnopithecus* and *Nasalis* (Figure 4C and Table 2). In African colobines, the long planum of *Colobus* is distinct from the shorter planum of *Procolobus* and *Piliocolobus*. Among the Cercopithecini, the long planum alveolare of *Miopithecus*, *Cercopithecus*, and *Erythrocebus* can be distinguished from the shorter planum of *Chlorocebus*, *Allochrocebus*, and *Allenopithecus* (Figure 4C and Table 2). In the papionins, the relatively long planum of *Macaca* and *Mandrillus* contrasts with the shorter ones of *Papio*, *Lophocebus* and *Theropithecus*.

**Figure 4:**
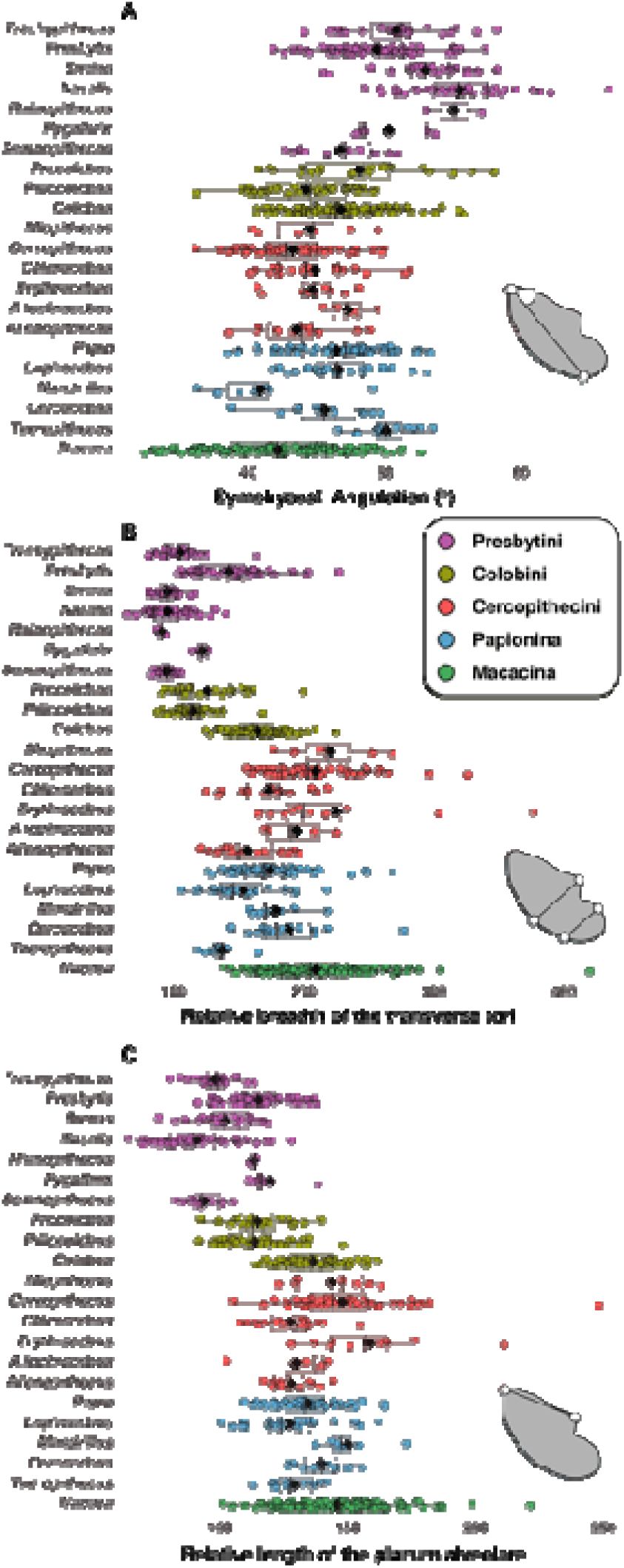
Boxplots of morphometric indices with A) symphyseal inclination, B) relative breadth of the transverse tori, and C) relative length of the planum alveolare. Means (black diamonds), medians (black bars), first quartile and third quartile are represented. An illustration of the measurement protocol is also provided in each panel.

### ALLOMETRY AND SIGNIFICANT DIFFERENCES IN PLS AND PGLS SLOPE VALUES

#### GM a surrogate of body mass

A highly significant value (*p* < 0.001) is observed for the PLS model of body mass versus GM, with an adjusted r^2^ of 0.60 (SOM Figure S3). This result justifies the use of GM as a proxy for body mass.

#### Phylogenetic generalized least squares regressions

For any given GM, the planum alveolare of cercopithecines is absolutely longer than that of colobines, with the exception of *Colobus* (Figure 5A). Similarly, the STT is absolutely larger in cercopithecines than in colobines, then again, with the exception of *Colobus* (Figure 5B). On the contrary, ITT is absolutely larger in colobines compared to most cercopithecines (Figure 5C). *Colobus* is also an exception to this pattern and is more similar to cercopithecines than to colobines. Similarly, *Theropithecus* and *Lophocebus* tend to present more developed ITT than other papionins of similar size. *Allenopithecus* is also distinct from the other Cercopithecini in having a more developed ITT relative to its GM (Figure 5C). The allometric patterns observed between PGLS and PLS are similar.

#### Partial least squares regressions

No significant difference is observed in PLS slope values between Cercopithecinae and Colobinae for planum alveolare length, STT breadth, and ITT breadth. Both subfamilies scale with positive allometry for all variables considered (Figure 6A-C and Table 3).

**Table 3:** Slope values of the PLS allometric regression per subfamily, tribe, and subtribe.

| *Taxa* | **Slope values** |  |
| --- | --- | --- |
|  | *Superior transverse torus* |  |
|  | 2.5 % | 97.5 % |
| *Cercopithecini* | 1.0617776857178 | 1.19704157758792 |
| *Colobini* | 1.24474904400434 | 1.44359947201397 |
| *Macacina* | 1.04088526441501 | 1.16866052102344 |
| *Papionina* | 1.26514297135779 | 1.37316775079648 |
| *Presbytini* | 0.880856808006998 | 1.05160980732505 |
| *Cercopithecinae* | 1.19301 | 1.238416 |
| *Colobinae* | 1.10004 | 1.24074 |
|  | *Inferior transverse torus* |  |
|  | 2.5 % | 97.5 % |
| *Cercopithecini* | 1.44344361867738 | 1.76929187093542 |
| *Colobini* | 0.88819811731835 | 1.30461429910937 |
| *Macacina* | 1.49737901293607 | 1.78711066091059 |
| *Papionina* | 1.26540607657995 | 1.4254570831541 |
| *Presbytini* | 1.85735391751641 | 2.15444898164911 |
| *Cercopithecinae* | 1.44214 | 1.53270 |
| *Colobinae* | 1.30546 | 1.61586 |
|  | *Planum alveolare* |  |
|  | 2.5 % | 97.5 % |
| *Cercopithecini* | 1.019938 | 1.2337819 |
| *Colobini* | 1.21756 | 1.474973 |
| *Macacina* | 0.837516 | 1.064517 |
| *Papionina* | 1.196959 | 1.315703 |
| *Presbytini* | 0.6817237 | 0.923796 |
| *Cercopithecinae* | 1.123564 | 1.188941 |
| *Colobinae* | 1.06302 | 1.303503 |

Apart from the isometry of Presbytini, all cercopithecid taxa have an STT that scales positively with GM (Figure 6A and Table 3). Highly significant differences in slope values are observed between Papionina and Macacina (*p* < 0.001; Tables 4 and 5) and between Papionina and Cercopithecini (*p* < 0.001). Differences in slope values are also highly significant between Colobini and Presbytini (*p* < 0.001).

**Table 4:**
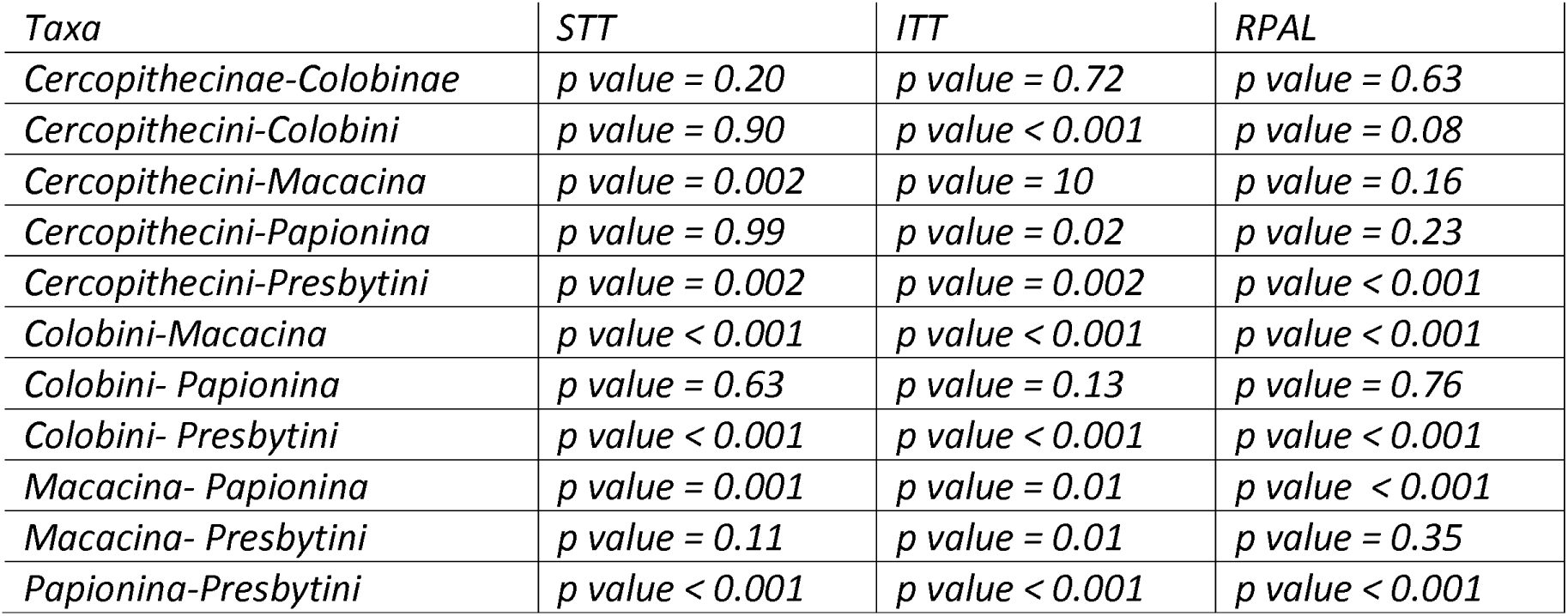
Differences in slope values (*p*-value) between subfamily, tribes and subtribes.

| *Taxa* | *STT* | *ITT* | *RPAL* |
| --- | --- | --- | --- |
| *Cercopithecinae-Colobinae* | *p value = 0.20* | *p value = 0.72* | *p value = 0.63* |
| *Cercopithecini-Colobini* | *p value = 0.90* | *p value < 0.001* | *p value = 0.08* |
| *Cercopithecini-Macacina* | *p value = 0.002* | *p value = 10* | *p value = 0.16* |
| *Cercopithecini-Papionina* | *p value = 0.99* | *p value = 0.02* | *p value = 0.23* |
| *Cercopithecini-Presbytini* | *p value = 0.002* | *p value = 0.002* | *p value < 0.001* |
| *Colobini-Macacina* | *p value < 0.001* | *p value < 0.001* | *p value < 0.001* |
| *Colobini- Papionina* | *p value = 0.63* | *p value = 0.13* | *p value = 0.76* |
| *Colobini- Presbytini* | *p value < 0.001* | *p value < 0.001* | *p value < 0.001* |
| *Macacina- Papionina* | *p value = 0.001* | *p value = 0.01* | *p value < 0.001* |
| *Macacina- Presbytini* | *p value = 0.11* | *p value = 0.01* | *p value = 0.35* |
| *Papionina-Presbytini* | *p value < 0.001* | *p value < 0.001* | *p value < 0.001* |

**Table 5:** Regression parameters of the PLS allometric regression per subfamily, tribe, and subtribe.

| *Taxa* | *Regression parameters (PLS)* |
| --- | --- |
|  | *Superior transverse torus* |
| *Cercopithecini* | $y = 1.13x - 0.80$ |
| *Colobini* | $y = 1.34x - 1.47$ |
| *Macacina* | $y = 1.10x - 0.66$ |
| *Papionina* | $y = 1.32x - 1.35$ |
| *Presbytini* | $y = 0.97x - 0.50$ |
|  | *Inferior transverse torus* |
| *Cercopithecini* | $y = 1.61x - 2.68$ |
| *Colobini* | $y = 1.10x - 1.14$ |
| *Macacina* | $y = 1.64x - 2.91$ |
| *Papionina* | $y = 1.34x - 1.93$ |
| *Presbytini* | $y = 2.00x - 3.42$ |
|  | *Planum alveolare* |
| *Cercopithecini* | $y = 1.13x - 0.38$ |
| *Colobini* | $y = 1.35x - 1.05$ |
| *Macacina* | $y = 0.95x + 0.19$ |
| *Papionina* | $y = 1.26x - 0.77$ |
| *Presbytini* | $y = 0.80x + 0.24$ |

Colobini is the only taxon with an ITT that scales isometrically while the other cercopithecid taxa scale positively (Figure 6B and Table 3). Slope value differences between Colobini and Presbytini are highly significant (*p* < 0.001; Tables 4 and 5). Similarly, significant differences are observed between Papionina and Macacina (*p* < 0.05; Tables 4 and 5).

The planum alveolare length of Macacina scales isometrically while that of Presbytini scales negatively (Figure 6C and Table 3). All other cercopithecid taxa have a planum alveolare length that scales positively.

**Figure 5:**
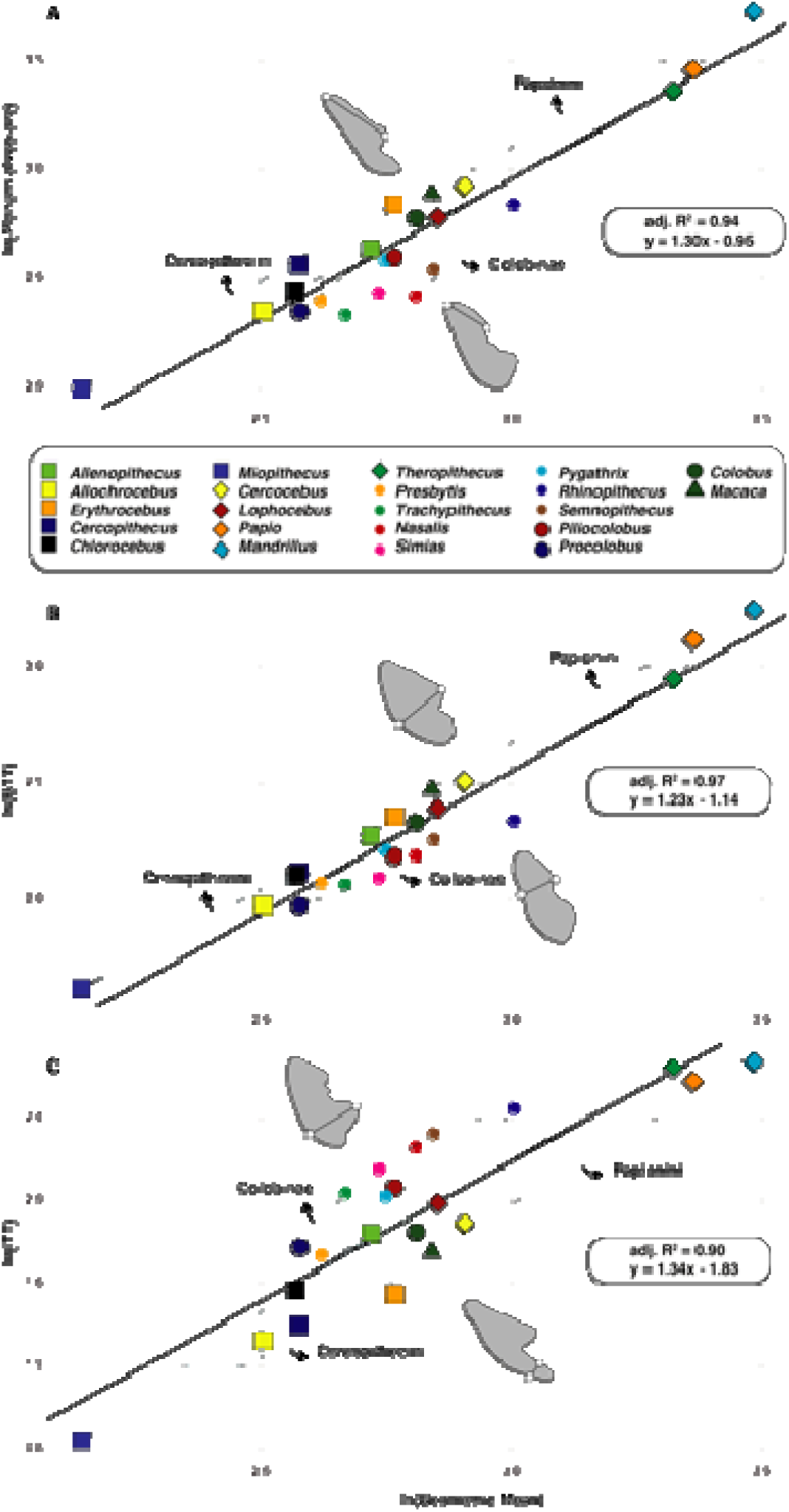
Phylogenetic generalized least squares regressions of A) the natural logarithm of planum alveolare length against the natural logarithm of size (i.e., GM), B) the natural logarithm of the superior transverse torus breadth against the natural logarithm of size, and C), the natural logarithm of the inferior transverse torus breadth against the natural logarithm of size.

**Figure 6:**
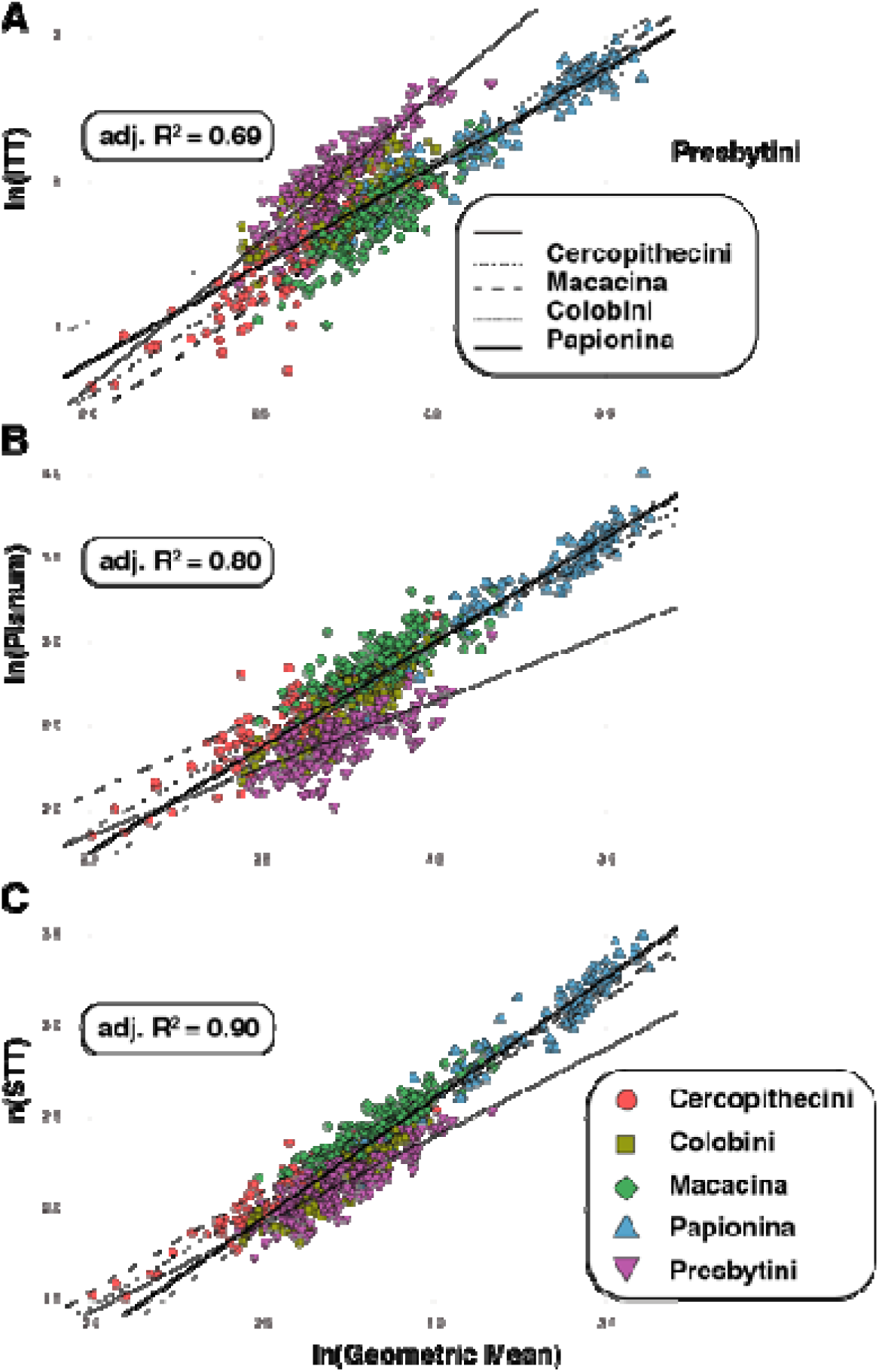
Partial least squares regressions of the natural logarithm of A) the inferior transverse torus breadth against the natural logaritm of size, B) the natural logarithm of planum alveolare length against the natural logarithm of size, and C) the natural logarithm of the superior transverse torus breadth against the natural logarithm of size

### SIGNIFICANT DIFFERENCES BETWEEN MALE AND FEMALE SPECIMENS (SEXUAL DIMORPHISM)

Significant differences are observed in the relative length of the planum alveolare between male and female specimens of *Chlorocebus aethiops* (p < 0.05), *Cercopithecus mitis* (p < 0.05), *Papio anubis* (p < 0.01), *Nasalis larvatus* (p < 0.05), *Procolobus verus* (p < 0.01) and *Colobus polykomos* (p < 0.001) (Figure 7A and Table 6). In each of the dimorphic taxa considered, the planum alveolare is relatively longer in males than in females (Figure 7A).

**Table 6:**
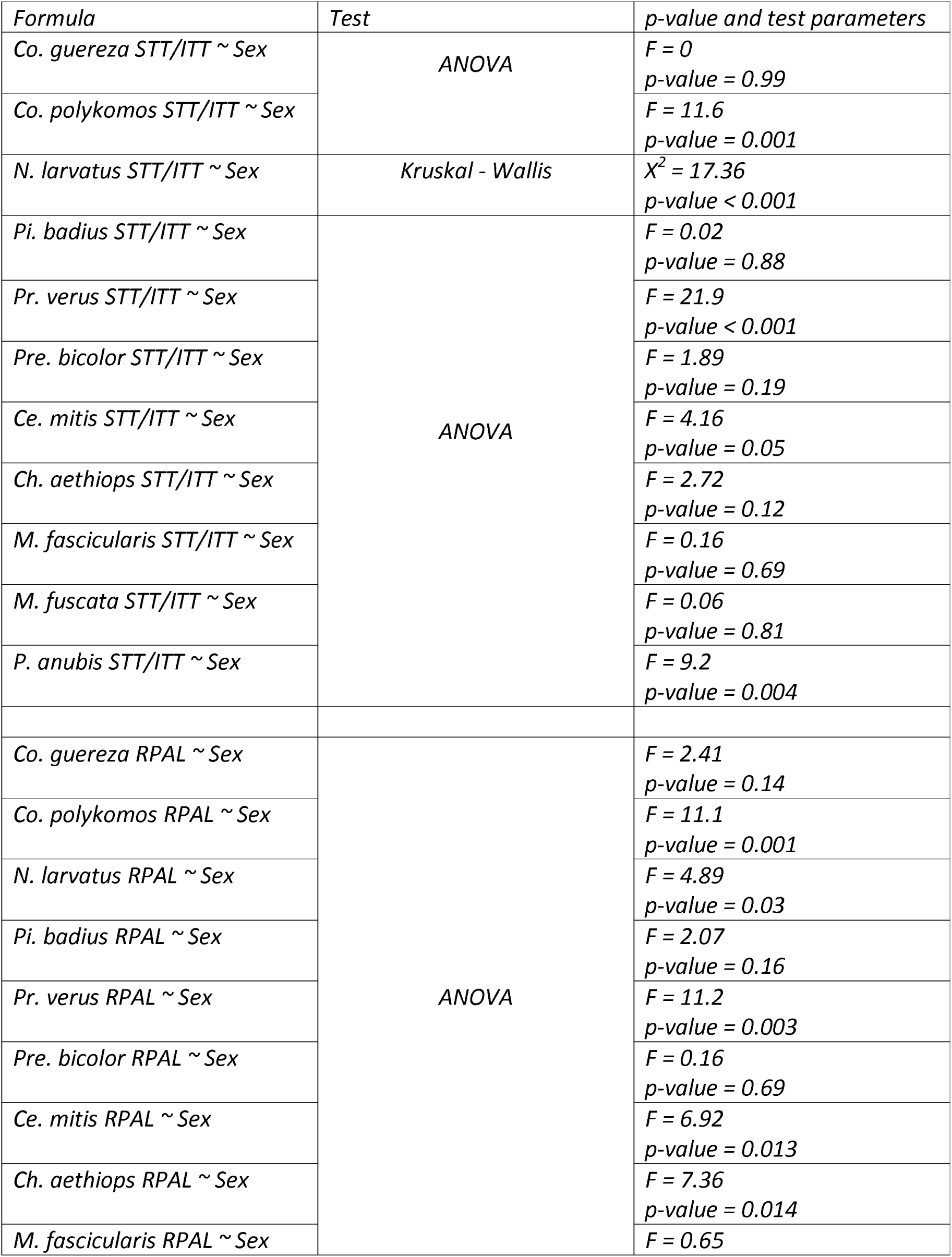
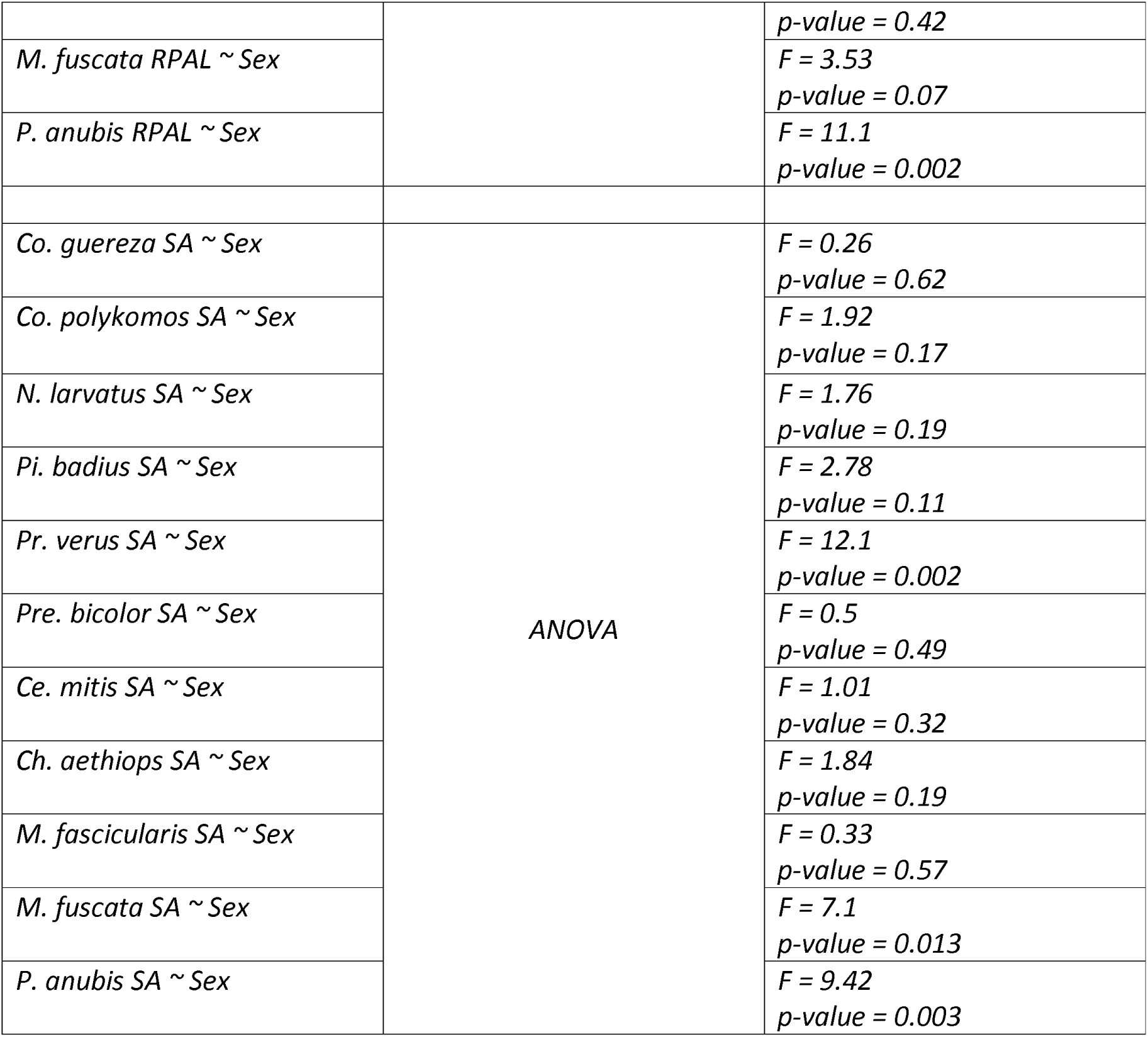
Significance and associated parameters of statistical models that test the effect of sexual dimorphism on morphometric ratios.

Significant differences are detected in the relative breadth of the transverse tori of male and female specimens of *Cer. mitis* (p < 0.05), *N. larvatus* (p < 0.001), *Pr. verus* (p < 0.001), *Co. polykomos* (p < 0.001), and *P. anubis* (*p* < 0.001) (Figure 7B and Table 6). In all dimorphic taxa, the transverse torus is relatively larger than the inferior torus in male specimens (Figure 7B).

Absolute values of the inclination of the symphysis are significantly distinct between male and female specimens of *Papio anubis* (p < 0.01), *Macaca fuscata* (p < 0.05), and *Procolobus verus* (p < 0.01) (Figure 8 and Table 6). The symphysis is more obtuse in males of *Papio anubis* and *Procolobus verus* while the reverse is true for *Macaca fuscata* where the symphysis is more obtuse in females than in males (Figure 8).

**Figure 7:**
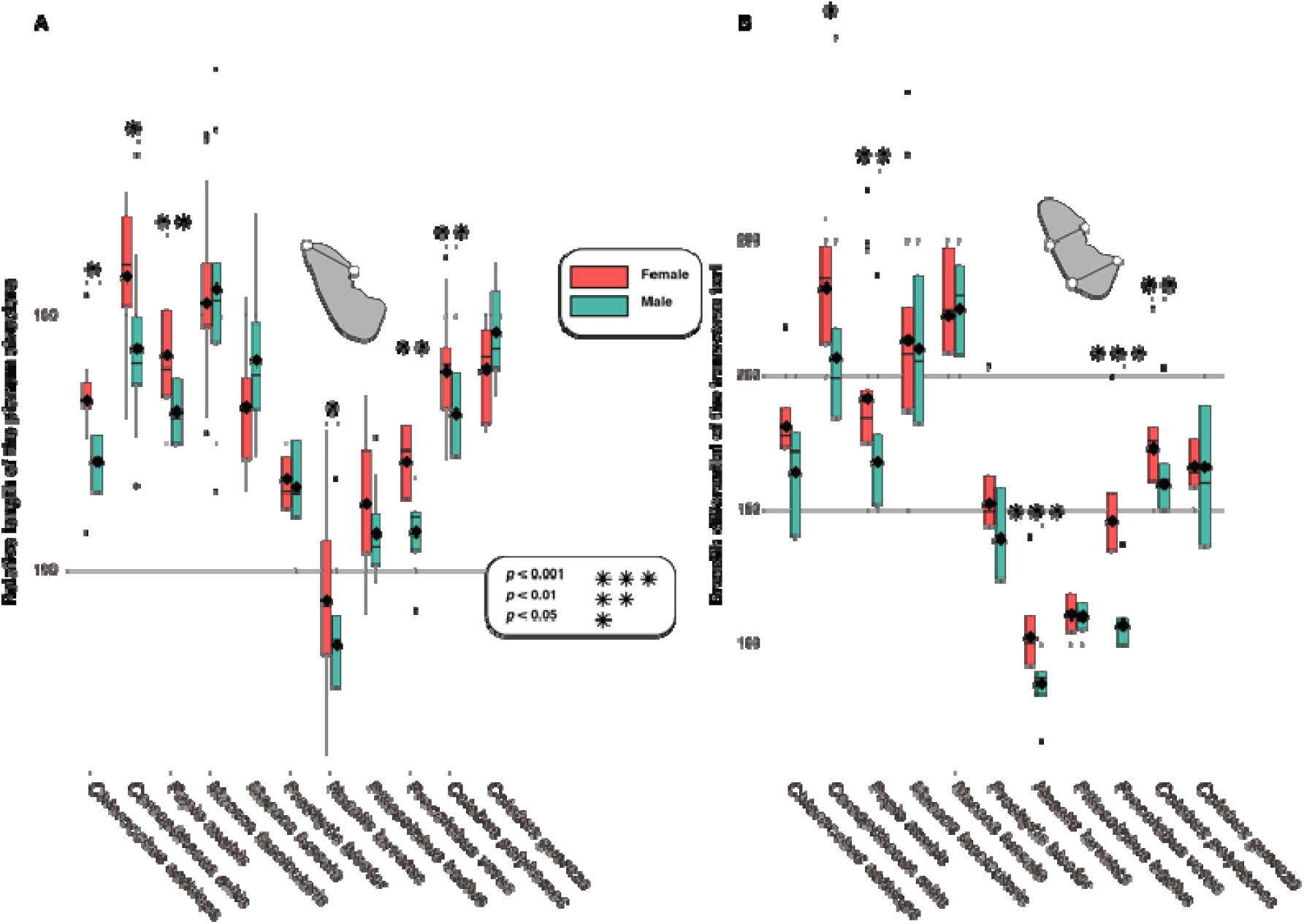
Boxplots of morphometric indices per sexes for selected cercopithecid taxa, with A) boxplots of the relative length of planum alveolare, and B) boxplots of the relative breadth of the transverse tori. Means (black diamonds), medians (black bars), first quartile and third quartile are represented. An illustration of the measurement protocol is also provided in each panel.

It is interesting to note the contrast in dimorphism between pairs of closely related colobine taxa. The symphysis of *Colobus polykomos* is notably dimorphic compared to that of *Colobus guereza* in relative length of the planum alveolare and relative breadth of the tori. Similarly, the symphysis of *Procolobus verus* is more dimorphic relative to that of *Piliocolobus badius*, not only with respect to the relative length of the planum alveolare and tori development, but also with respect to the symphyseal inclination. *Macaca fascicularis*, *Presbytis bicolor*, *Piliocolobus badius*, and *Colobus guereza* are all monomorphic in symphyseal shape.

Sexual dimorphism is also expressed differently in the mandible of dimorphic taxa. While *Chlorocebus aethiops* is dimorphic in planum alveolare length, it is not in relative breadth of the tori. An identical pattern is observed in *Papio anubis*. *Colobus polykomos* and *Nasalis larvatus* are dimorphic in relative breadth of the tori and planum alveolare length, but not in symphyseal inclination.

**Figure 8:**
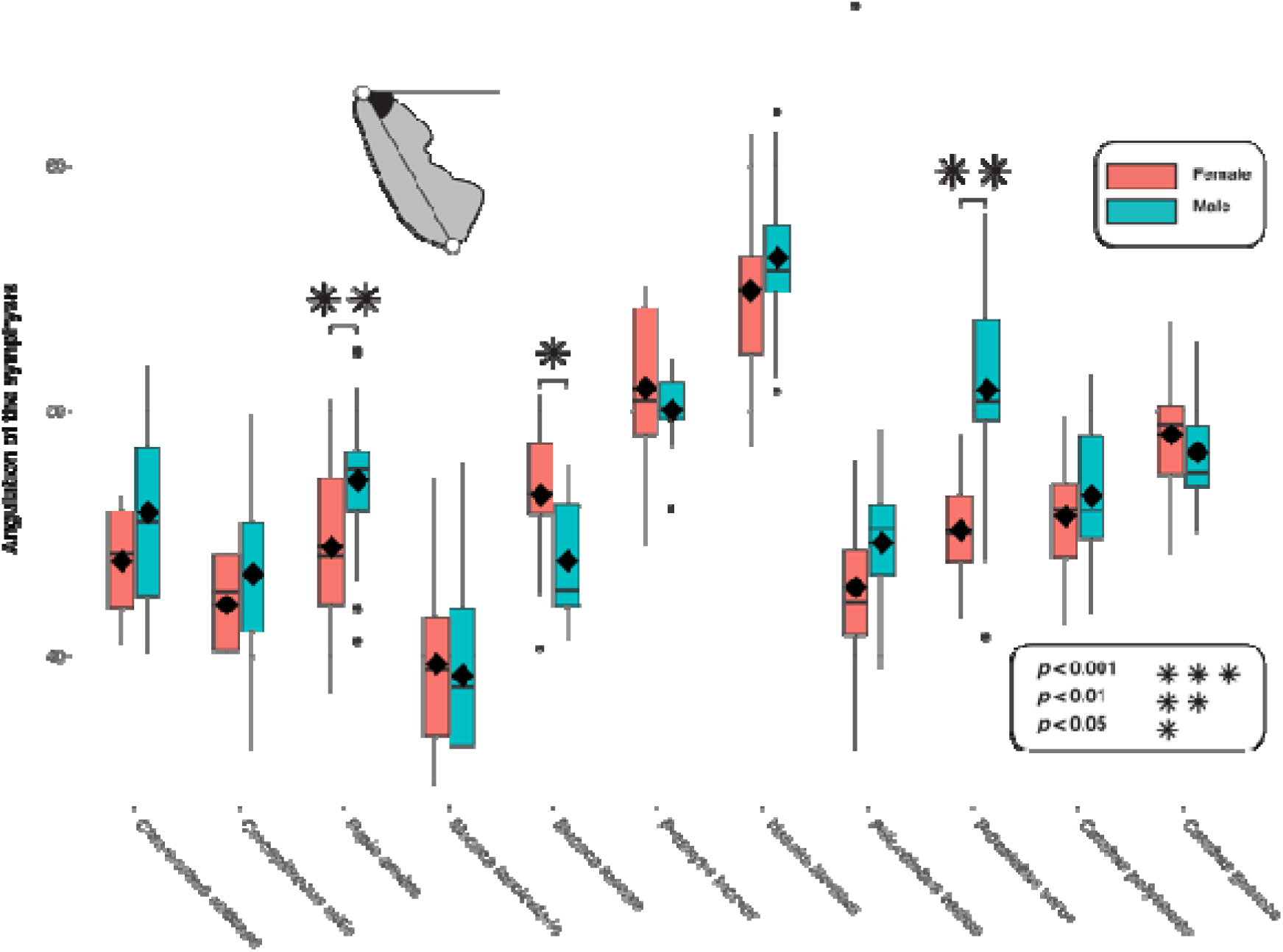
Boxplot of symphyseal inclination per sexes for selected cercopithecid taxa. Means (black diamonds), medians (black bars), first quartile and third quartile are represented.

### SIGNIFICANT DIFFERENCES BETWEEN SUBFAMILIES AND TRIBES IN MULTIVARIATE AND UNIVARIATE DATA

#### Multivariate data

A highly significant difference (*p* < 0.0001) is observed between Cercopithecinae and Colobinae on PC1 scores, but not on PC2 (*p* = 0.2) scores (Table 7).

All pairs of tribes and subtribes, with the exception of the Papionina-Macacina pair (*p* = 1.0), can be discriminated with a high level of statistical significance (*p* < 0.0001 and *p* < 0.001) on PC1 scores (Table 7). The same is true for PC2 scores, with the exception of the nonsignificant difference between Macacina-Colobini (*p* = 0.2). Unlike PC1 scores, Macacina and Papionina can be significantly discriminated on PC2 scores (Table 7).

#### Univariate data

Colobinae and Cercopithecinae can be significantly (*p* < 0.001) distinguished on the basis of the STT/ITT ratio (Table 8). Similarly, the two subfamilies are significantly distinct (*p* < 0.001) on relative length of the planum alveolare and symphyseal inclination (Table 8). Most pairs of tribes and subtribes can be differentiated with a high level of significance (*p* < 0.001) using the STT/ITT ratio. The only exception is the marginal significant difference between the Macacina-Cercopithecini pair (*p* = 0.03). Similarly, most pairs of tribes and subtribes are highly distinct (*p* < 0.001) in regard to the relative length of planum alveolare. However, while Colobini and Papionina are marginally distinct (*p* = 0.02), no significant differences are observed between Cercopithecini and Macacina and between Cercopithecini and Papionina (Table 8). Symphyseal inclination is also a powerful measure for discriminating cercopithecid tribes and subtribes, with highly significant differences for most pairs of taxa (Table 8). Cercopithecini are, however, marginally distinct from Macacina (*p* = 0.05), and no significant differences are found between Colobini and Papionina (*p* = 0.42).

**Table 7:**
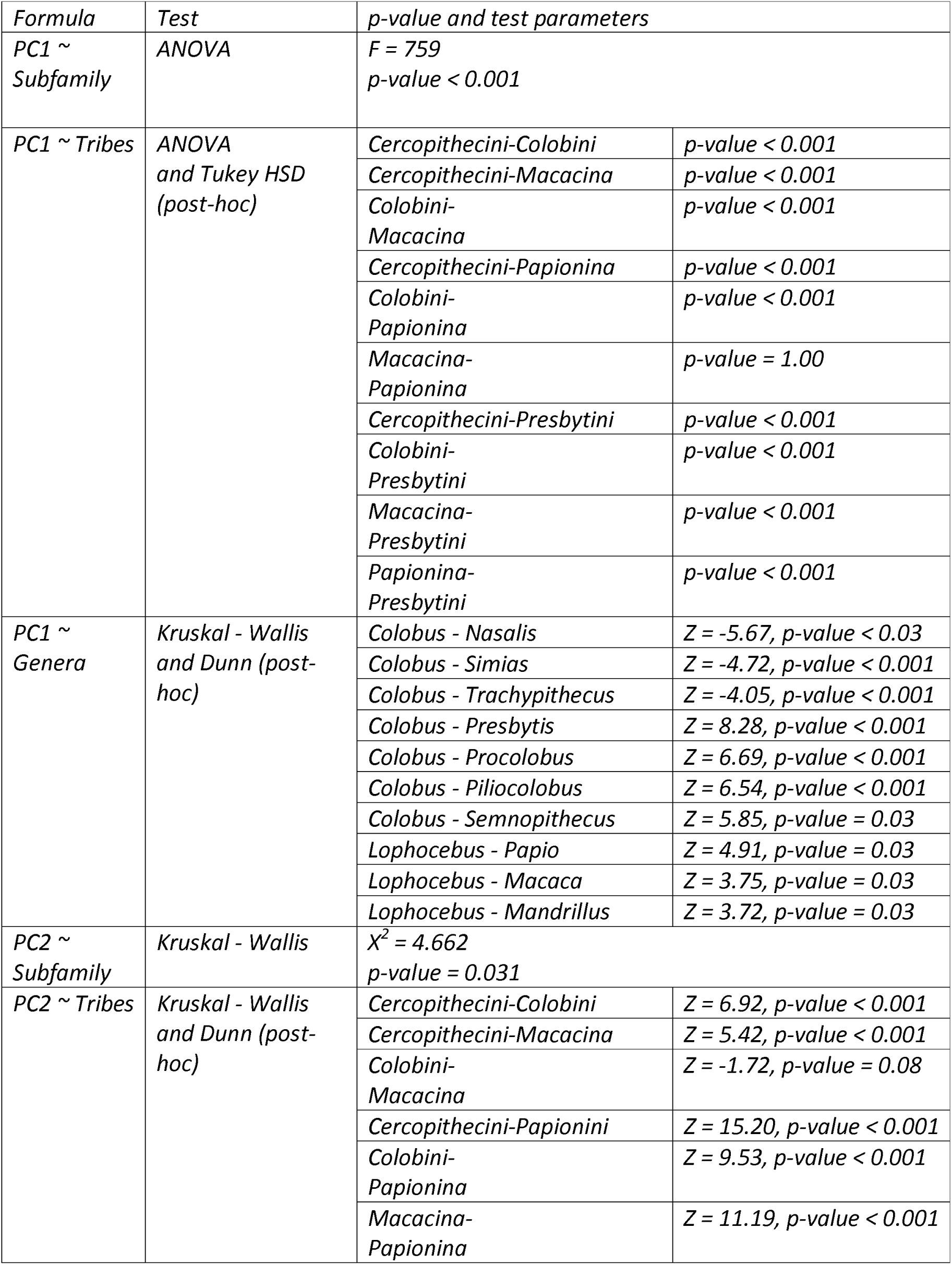
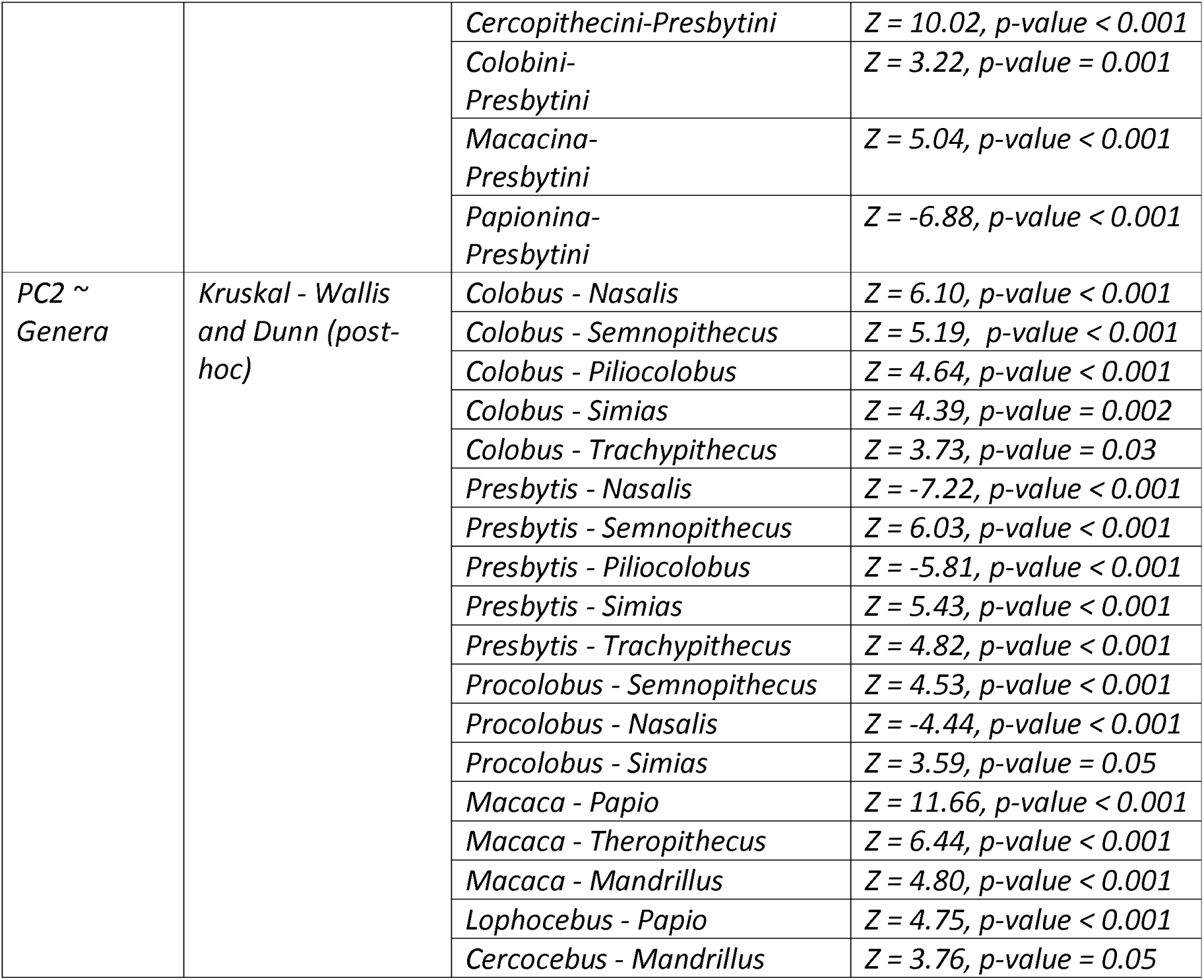
Significance and associated parameters of statistical models that test the effect of sexual dimorphism on morphometric ratios.

| Formula | Test | p-value and test parameters |  |
| --- | --- | --- | --- |
| PC1 ~ Subfamily | ANOVA | F = 759<br>p-value < 0.001 |  |
| PC1 ~ Tribes | ANOVA and Tukey HSD (post-hoc) | Cercopithecini-Colobini | p-value < 0.001 |
|  |  | Cercopithecini-Macacina | p-value < 0.001 |
|  |  | Colobini-Macacina | p-value < 0.001 |
|  |  | Cercopithecini-Papionina | p-value < 0.001 |
|  |  | Colobini-Papionina | p-value < 0.001 |
|  |  | Macacina-Papionina | p-value = 1.00 |
|  |  | Cercopithecini-Presbytini | p-value < 0.001 |
|  |  | Colobini-Presbytini | p-value < 0.001 |
|  |  | Macacina-Presbytini | p-value < 0.001 |
|  |  | Papionina-Presbytini | p-value < 0.001 |
| PC1 ~ Genera | Kruskal - Wallis and Dunn (post-hoc) | Colobus - Nasalis | Z = -5.67, p-value < 0.03 |
|  |  | Colobus - Simias | Z = -4.72, p-value < 0.001 |
|  |  | Colobus - Trachypithecus | Z = -4.05, p-value < 0.001 |
|  |  | Colobus - Presbytis | Z = 8.28, p-value < 0.001 |
|  |  | Colobus - Procolobus | Z = 6.69, p-value < 0.001 |
|  |  | Colobus - Piliocolobus | Z = 6.54, p-value < 0.001 |
|  |  | Colobus - Semnopithecus | Z = 5.85, p-value = 0.03 |
|  |  | Lophocebus - Papio | Z = 4.91, p-value = 0.03 |
|  |  | Lophocebus - Macaca | Z = 3.75, p-value = 0.03 |
|  |  | Lophocebus - Mandrillus | Z = 3.72, p-value = 0.03 |
| PC2 ~ Subfamily | Kruskal - Wallis | $\chi^2 = 4.662$<br>p-value = 0.031 | |
| PC2 ~ Tribes | Kruskal - Wallis and Dunn (post-hoc) | Cercopithecini-Colobini | Z = 6.92, p-value < 0.001 |
|  |  | Cercopithecini-Macacina | Z = 5.42, p-value < 0.001 |
|  |  | Colobini-Macacina | Z = -1.72, p-value = 0.08 |
|  |  | Cercopithecini-Papionini | Z = 15.20, p-value < 0.001 |
|  |  | Colobini-Papionina | Z = 9.53, p-value < 0.001 |
|  |  | Macacina-Papionina | Z = 11.19, p-value < 0.001 |
|  |  | *Cercopithecini-Presbytini* | *Z = 10.02, p-value < 0.001* |
|  |  | *Colobini-Presbytini* | *Z = 3.22, p-value = 0.001* |
|  |  | *Macacina-Presbytini* | *Z = 5.04, p-value < 0.001* |
|  |  | *Papionina-Presbytini* | *Z = -6.88, p-value < 0.001* |
| *PC2 ~ Genera* | *Kruskal - Wallis and Dunn (post-hoc)* | *Colobus - Nasalis* | *Z = 6.10, p-value < 0.001* |
|  |  | *Colobus - Semnopithecus* | *Z = 5.19, p-value < 0.001* |
|  |  | *Colobus - Piliocolobus* | *Z = 4.64, p-value < 0.001* |
|  |  | *Colobus - Simias* | *Z = 4.39, p-value = 0.002* |
|  |  | *Colobus - Trachypithecus* | *Z = 3.73, p-value = 0.03* |
|  |  | *Presbytis - Nasalis* | *Z = -7.22, p-value < 0.001* |
|  |  | *Presbytis - Semnopithecus* | *Z = 6.03, p-value < 0.001* |
|  |  | *Presbytis - Piliocolobus* | *Z = -5.81, p-value < 0.001* |
|  |  | *Presbytis - Simias* | *Z = 5.43, p-value < 0.001* |
|  |  | *Presbytis - Trachypithecus* | *Z = 4.82, p-value < 0.001* |
|  |  | *Procolobus - Semnopithecus* | *Z = 4.53, p-value < 0.001* |
|  |  | *Procolobus - Nasalis* | *Z = -4.44, p-value < 0.001* |
|  |  | *Procolobus - Simias* | *Z = 3.59, p-value = 0.05* |
|  |  | *Macaca - Papio* | *Z = 11.66, p-value < 0.001* |
|  |  | *Macaca - Theropithecus* | *Z = 6.44, p-value < 0.001* |
|  |  | *Macaca - Mandrillus* | *Z = 4.80, p-value < 0.001* |
|  |  | *Lophocebus - Papio* | *Z = 4.75, p-value < 0.001* |
|  |  | *Cercocebus - Mandrillus* | *Z = 3.76, p-value = 0.05* |

**Table 8:**
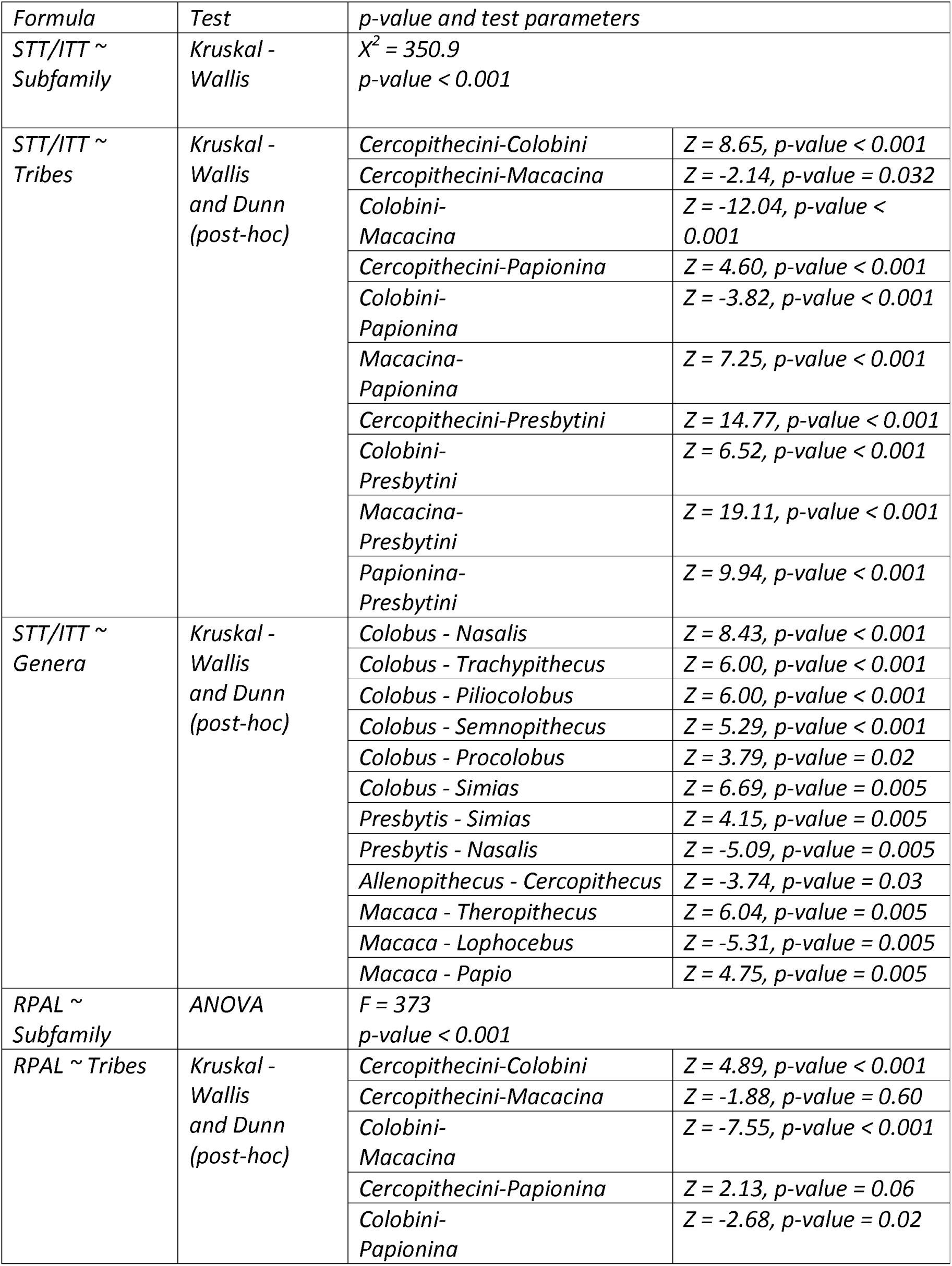
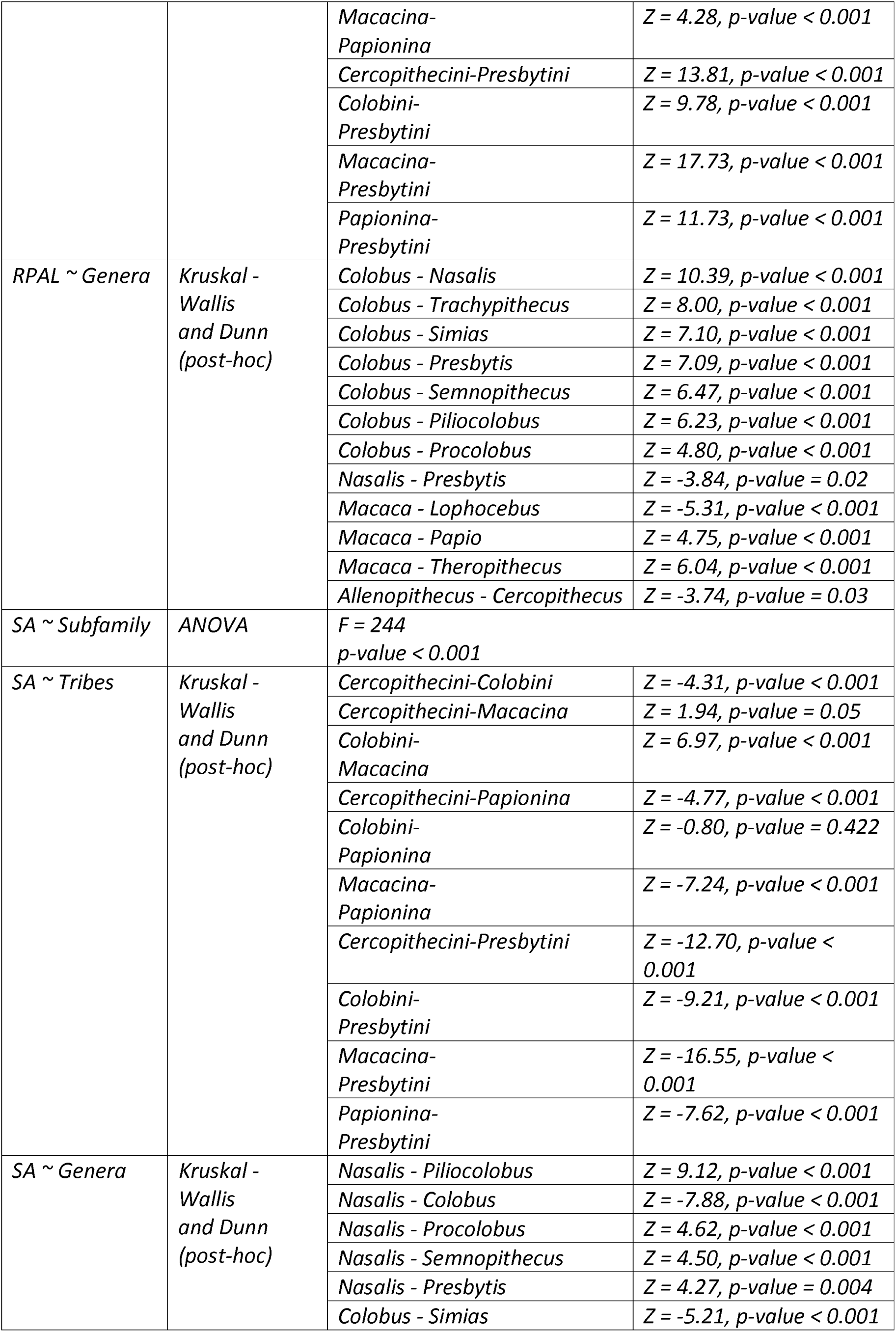
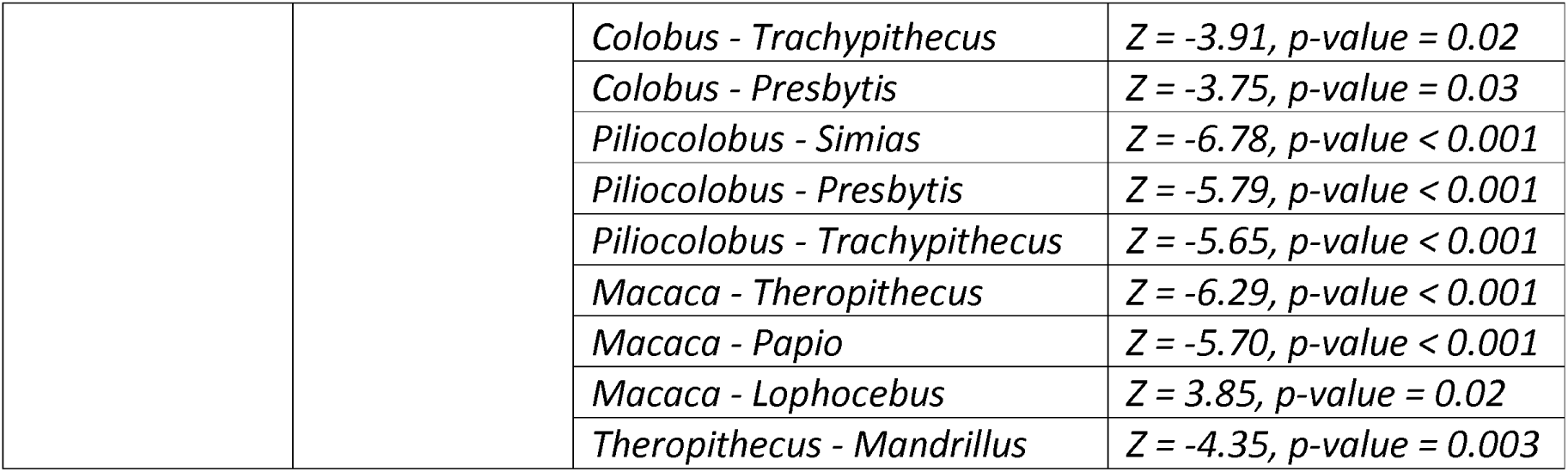
Significance and associated parameters of statistical models that test the effect of taxonomy on morphometric ratios.

### SIGNIFICANT DIFFERENCES BETWEEN GENERA IN MULTIVARIATE AND UNIVARIATE DATA

#### Multivariate data

Among Papionina, *Lophocebus* can be significantly discriminated from *Mandrillus, Papio,* and *Macaca* on PC1 scores (*p* < 0.001). Indeed, *Lophocebus* has significantly lower PC1 scores compared to *Mandrillus*, *Papio*, and *Macaca* (Table 7). On PC2 scores, *Cercocebus* is significantly different from *Mandrillus* (*p*-values < 0.001), reflecting its positive PC2 scores (Table 7). *Lophocebus* is also significantly distinct from *Papio* (*p* < 0.001) on PC2 scores, indicating its comparatively high scores (Table 7). Also on PC2, the positive scores of *Macaca* can be significantly discriminated with high confidence from the negative scores of *Mandrillus*, *Papio* and *Theropithecus*.

We detected no significant differences in PC1 and PC2 scores among Cercopithecini (Table 7). As for the Colobini, *Colobus* is significantly distinct from most colobines on PC1 scores (Table 7). These significant differences underline the positive scores of *Colobus* on PC1. *Pygathrix* and *Rhinopithecus* are not statistically distinct from *Colobus*. This lack of distinction may be explained by the low sample size of *Pygathrix* and *Rhinopithecus* and the use of non-parametric test (SOM Tables S4 and S6). Indeed, when looking at the boxplots of the relative breadth of the transverse tori and relative length of the planum alveolare, *Pygathrix* and *Rhinopithecus* fall in the lower quartile or outside the range of variation of *Colobus* (Figure 4B-C). On PC2 scores, *Colobus* is also significantly distinct from most colobines, with the exception of *Presbytis* and *Procolobus*. *Procolobus* is also significantly different from the Asian colobines *Semnopithecus*, *Nasalis*, and *Simias* on PC2 (Table 7).

Among the Presbytini, *Presbytis* is significantly different from *Nasalis*, *Simias, Semnopithecus*, and *Trachypithecus* on PC2 scores.

#### Univariate data

The high STT/ITT ratio of *Theropithecus* is significantly distinct, at a high significance level (*p* < 0.001), from that of *Macaca* (Figure 9 and Tables 8 and 9). The latter also has a significantly distinct STT/ITT ratio from *Papio* (*p* < 0.001) and *Lophocebus* (*p* < 0.001), illustrating its marked STT/ITT ratio (Table 8). The values of the relative length of the planum alveolare of *Macaca* are significantly distinct from that of *Lophocebus* (*p* < 0.001), *Papio* (*p* < 0.001) and *Theropithecus* (*p* < 0.01), illustrating the relatively long planum alveolare of *Macaca*. It also has an extremely inclined symphysis, with inclination values significantly distinct from that of *Theropithecus*, *Papio*, and *Lophocebus* (Table 8). In contrast to *Macaca*, the symphysis of *Theropithecus* is quite steep and significantly different from that of *Mandrillus* (*p* = 0.003).

Based on the STT/ITT ratio, *Allenopithecus* can be differentiated from *Cercopithecus*, *Erythrocebus*, and *Miopithecus*, with *p* < 0.001 for each comparison (Table 8). These differences are related to the low STT/ITT ratio values of *Allenopithecus* (Figure 9 and Table 9). *Chlorocebus* also has a low STT/ITT ratio, significantly distinct from *Cercopithecus* (*p* < 0.0001), *Chlorocebus* (*p* < 0.001), and *Miopithecus* (*p* < 0.05). *Allenopithecus* is also significantly distinct from *Cercopithecus* (*p* < 0.001) in regard to the relative planum alveolare length. No significant difference is observed in symphyseal inclination among Cercopithecini (Table 8).

**Table 9:** Summary of the main symphyseal characteristics that permit to distinguished cercopithecids at various taxonomic levels.

| *Taxa* | *Main symphyseal characteristics of the considered taxon* |
| --- | --- |
| *Colobinae* | Short and steep symphysis relative to cercopithecines ($49.1 \pm 4.7^\circ$), well-developed ITT with a breadth differential of the tori less pronounced than in cercopithecines ($128.5 \pm 33.1$), and a planum alveolare relatively short compared to cercopithecines ($113.9 \pm 19.6$). |
| *Cercopithecinae* | Long and inclined symphysis relative to colobines ($43.9 \pm 4.3^\circ$), well-developed STT with a breadth differential of the tori more pronounced than in cercopithecines ($192.6 \pm 40.7$), and a planum alveolare relatively long compared to cercopithecines ($140.5 \pm 18.3$). |
| *Papionina* | Steep symphysis relative to Macacina ($46.5 \pm 3.7^\circ$), well-developed STT with a breadth differential of the tori less pronounced than in Macacina ($166.7 \pm 30.7$), and a planum alveolare relatively short compared to Macacina ($133.4 \pm 13.3$). Major differences within the taxon are illustrated by the steep symphysis and developed ITT of *Theropithecus* and the highly inclined symphysis of *Mandrillus*.<br>In *Papio anubis*, sexual dimorphism is expressed in relative length of the planum alveolare, transverse tori development, and symphyseal inclination. |
| *Macacina* | Inclined symphysis relative to Papionina ($42.2 \pm 4.5^\circ$), well-developed STT with a breadth differential of the tori more pronounced than in Papionina ($208.5 \pm 37.1$), and a planum alveolare relatively long compared to Papionina ($145.2 \pm 18.1$).<br>In *Macaca fascicularis* and *Macaca fuscata*, sexual dimorphism is poorly expressed in symphyseal traits, and only visible in symphyseal inclination values of *Macaca fuscata*. |
| *Cercopithecini* | Similar inclination of the symphysis as that of Papionini ($43.9 \pm 3.2^\circ$), well-developed STT with a breadth differential of the tori slightly more pronounced than in Papionini ($196.5 \pm 42.0$), and a planum alveolare slightly longer than that of Papionini ($141.1 \pm 20.7$). A major difference within the taxon is illustrated by the developed ITT of *Allenopithecus*.<br>Sexual dimorphism in symphyseal traits is moderate in *Chlorocebus aethiops* and *Cercopithecus mitis*, but is nonetheless influencing the relative length of the planum alveolare and transverse tori development. |
| *Colobini* | *Inclined symphysis relative to Presbytini ($46.3 \pm 3.7^\circ$), well-developed STT with a breadth differential of the tori more pronounced than in Presbytini ($145.7 \pm 30.0$), and a planum alveolare relatively long compared to Presbytini ($126.9 \pm 15.9$).*<br>*Major differences within the taxon are highlighted by the reduced ITT and relatively long planum alveolare of Colobus, and the inclined symphysis of Piliocolobus.*<br>*Sexual dimorphism in symphyseal shape is marked in Procolobus verus and moderate in Colobus polykomos.* |
| *Presbytini* | *Steep symphysis relative to Colobini ($51.4 \pm 4.3^\circ$), well-developed ITT with a breadth differential of the tori less pronounced than in Colobini ($113.9 \pm 28.3$), and a planum alveolare relatively short compared to Colobini ($102.8 \pm 15.1$).*<br>*Major differences within the taxon are highlighted by the reduced ITT and relatively long planum alveolare of Presbytis, and the inclined symphysis of Semnopithecus.*<br>*Sexual dimorphism in symphyseal shape is marked in Nasalis larvatus.* |

Among the Colobini, *Colobus* is significantly distinct from *Procolobus* (*p* < 0.001) and *Piliocolobus* (*p* < 0.001) due to the high mean value of its STT/ITT ratio (Figure 9 and Tables 8 and 9). The relatively long planum alveolare of *Colobus* is also significantly distinct from that of *Piliocolobus* (*p* < 0.001) and *Procolobus* (*p* < 0.001). *Presbytis* can be significantly differentiated from other Asian colobines on the basis of its high STT/ITT ratio values, particularly in relation to *Nasalis*, *Semnopithecus*, *Simias* and *Trachypithecus* (Table 8). The relatively long planum alveolare of *Presbytis* is significantly different from *Nasalis* (*p* < 0.001), *Trachypithecus* (*p* < 0.001), *Semnopithecus* (*p* < 0.001), and *Simias* (*p* < 0.05). The symphysis of *Nasalis* is extremely steep and can be significantly distinguished from that of African colobines, *Semnopithecus* and *Presbytis* (Table 8). The distinctly sloping symphysis of *Colobus* and *Piliocolobus* is also significantly different from that of *Trachypithecus*, *Simias*, and *Presbytis*.

### PHYLOGENETIC SIGNAL IN MULTIVARIATE AND UNIVARIATE DATA

#### Multivariate data

PC1 scores follow a distribution expected under a Brownian Motion (BM) model with a Pagel’s λ equal to 0.8 (*p* < 0.001), contrary to that of PC2 and PC3 scores, which does not follow a BM model (*p* = 0.06 in each case).

#### Univariate data

Data follow a distribution expected under a BM model for the STT/ITT ratio and relative length of the planum alveolare, with a Pagel’s λ equal to 1 (*p* < 0.001) and 0.9 (*p* < 0.001), respectively.

**Figure 9:**
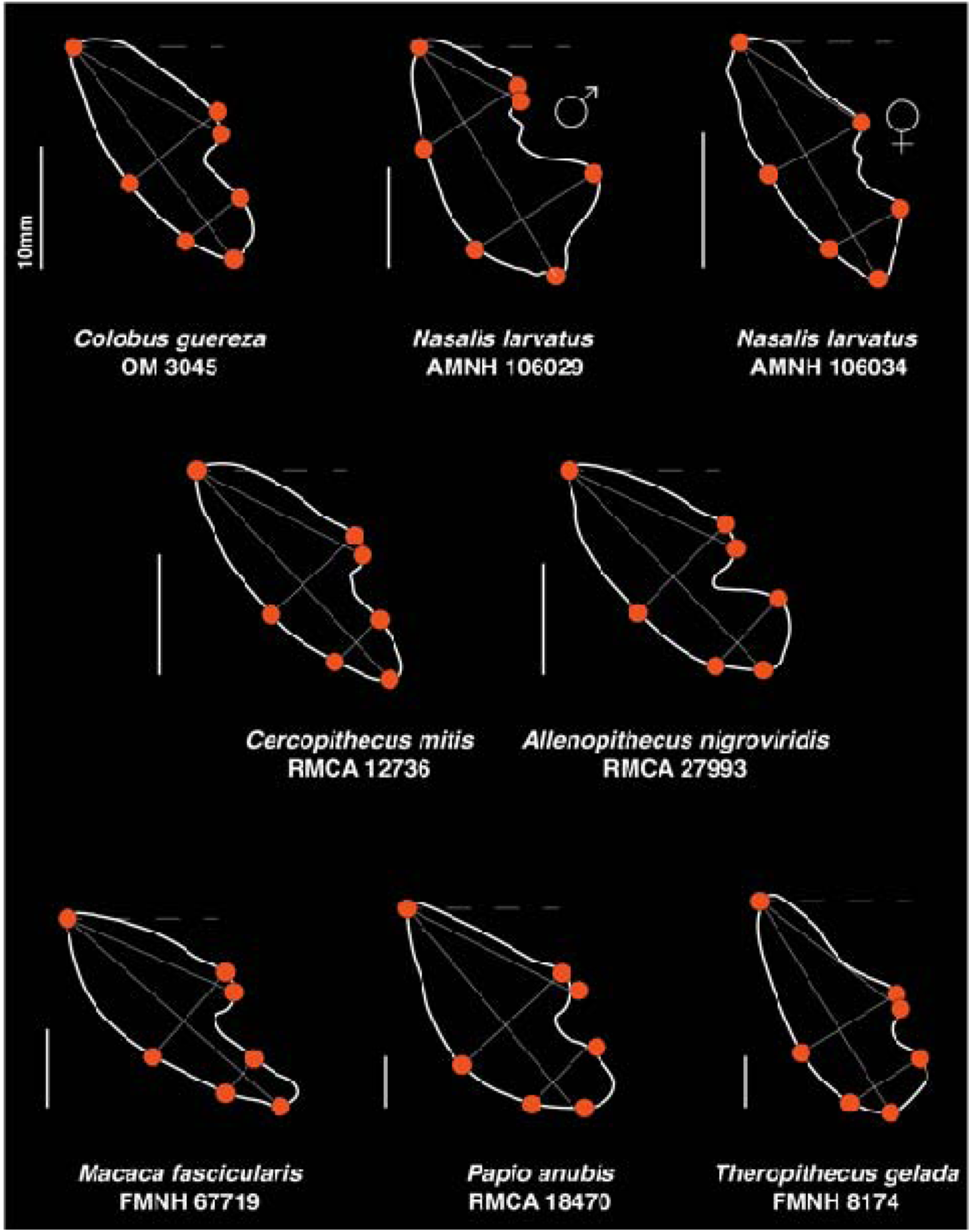
Illustration of the main symphyseal differences highlighted in this study. Not the shortened ITT and elongated planum alveolare of *Colobus* compared to *Nasalis*, the high sexual dimorphism in transverse tori development and planum alveolare length in *Nasalis*, the reduced ITT and long planum alveolare of *Cercopithecus mitis* compared to *Allenopithecus nigroviridis*, the inclined symphysis and reduced ITT of *Macaca fascicularis*, and the steep symphysis and developed ITT of *Theropithecus gelada* compared to *Papio anubis*.

## DISCUSSION

This study aimed to identify differences among cercopithecid taxa on the grounds of symphyseal traits and to identify the source of variation in such traits by evaluating allometric trajectories and sexual dimorphism. By showing clear phenetic differences between cercopithecid taxa, we join the large body of data that has contributed to this matter in hominoids (Daegling, 1993; Brown, 1997; Daegling and Jungers, 2000; Taylor, 2002; Sherwood et al., 2005; Guy et al., 2008), and cercopithecids (Hylander, 1985; Ravosa, 1996; Vinyard and Ravosa, 1998; Pallas et al., 2019; Locke et al., 2020), underscoring the usefulness of the symphysis in taxonomy. We also demonstrate the presence of significant sexual differences in symphysis shape among several cercopithecines and colobines species in addition to firmly demonstrate the relevance of tori development, planum alveolare length, and symphyseal inclination in sex identification. We also point to distinct allometric trajectories between colobine and cercopithecines in regard to tori development and planum alveolare length.

In this section, we discuss the implication of the identified symphyseal differences on the functional anatomy and evolution of cercopithecids. In particular, we emphasize the relevance of our results to taxonomic issues in extant and fossil cercopithecids. Finally, we comment on the confounding effect of sexual dimorphism in the shape of the symphysis and stress the need to identify its proximate causes.

### Impact of symphyseal shape on the cercopithecid functional morphology and evolution

Our ratios and morphometric variables significantly distinguished the cercopithecid subfamilies, and notably underlined the short planum alveolare of colobines, the short ITT of cercopithecines (especially for *Macaca* and *Cercopithecus*), the large ITT of colobines, and the relatively long symphysis of cercopithecines (Figure 9 and Table 9). We also proved that the values of relative length of the planum alveolare and relative breadth of the transverse tori were not randomly distributed in the cercopithecid phylogenetic tree but rather followed a phylogenetically structured distribution, as expected under Brownian motion. This implies that, at a broad taxonomic scale, the symphyseal shape of extant cercopithecids is constrained primarily by phylogeny. Nevertheless, we have identified several cases where closely related genera, such as *Colobus* and *Piliocolobus*, diverge in shape, demonstrating that differences in selective pressures, and associated morphological optima, can also be identified. Due to the limited sample size for some taxa, this study focused only on phylogenetic inferences at the genus level. Additional studies targeting species-level differences may reveal distinct selective pressures and selective regimes in closely related species. Speciose taxa such as *Cercopithecus* or *Piliocolobus* are prime candidates for such studies.

Most symphyseal traits of extant cercopithecids scale positively against body mass. The isometric scaling of the STT breadth of Presbytini, the ITT breadth of Colobini, and planum alveolare length of Macacina and Presbytini are exception to this pattern. Overall, the allometric regressions presented here demonstrated that the planum alveolare of cercopithecines is longer, the STT larger, and the ITT shorter than those of similarly sized colobines. These results are consistent with previous functional analyses that demonstrated that the symphyseal width, and more precisely the STT, of cercopithecines scales positively with body mass (Hylander, 1984; Ravosa, 1996; Vinyard and Ravosa, 1998). We complement these analyses by showing that while the structural robustness of the cercopithecine symphysis is determined by the breadth of the STT, that of colobines is driven by ITT breadth. *Colobus* and *Allenopithecus* are exceptions and present distinct allometric pattern compared to their closest representatives.

Hylander (1984) hypothesized that an expanded STT was a structural response to stresses generated by lateral transverse bending (i.e., wishboning), and subsequently demonstrated empirically that it was a highly strained area during simulated wishboning (Daegling et al., 2009; Bucinell et al., 2010). Other analyses have confirmed that obliquely inclined symphysis with an enlarged STT are better able to resist lateral transverse bending (Daegling and McGraw, 2001; Panagiotopoulou and Cobb, 2011). In contrast, an enlarged ITT is not significantly involved in attenuating stresses engendered by wishboning, but rather mitigates stresses engendered by sagittal bending and dorsoventral shear of the symphysis during mastication (Panagiotopoulou and Cobb, 2011). Here, we showed that the ITT of colobines is wider than that of cercopithecines. This reflects greater selective pressures on the colobine symphysis to resist dorsoventral shear rather than lateral transverse bending, as previously suggested on the basis of their deep corpora and mechanical properties of their diet (Hylander, 1979; Bouvier, 1986; Ravosa, 1996; Jablonski et al., 1998). Exception to the tendency of presenting an enlarged ITT nonetheless exists among colobines, with *Colobus* being the most distinct taxon.

*Colobus* presents a unique suite of characters among colobines, that is, a long planum alveolare, a reduced ITT, and a developed STT (Figure 9 and Table 9). These results support the observation of Benefit and Pickford (1986) regarding the diminutive ITT of the genus. However, uncertainty in the phylogenetic polarity of these traits precludes us from including them in a revised diagnosis of the genus, but complementary analyses that includes fossil colobines should resolve this issue (Pallas et al., in prep.). Nonetheless, the distinct symphyseal anatomy of *Colobus* should allow for future identification of *Colobus* representatives and refinement of taxonomic hypotheses in the fossil record, notably from the Pleistocene deposits of Koobi Fora, Shungura and Asbole (Leakey, 1987; Frost and Alemseged, 2007; Jablonski et al., 2008).

Despite differences in diet, *Colobus guereza* and *Colobus polykomos* do not differ in relative tori development. Indeed, *Co. polykomos* has a diet that includes, at a higher frequency, challenging items mechanically (seeds) compared to *Co. guereza* (Daegling and McGraw, 2001; Korstjens and Galat-Luong, 2013), with the latter preferring leaves and pulpy fruits (Fashing, 2001; Chapman et al., 2004; Fashing and Oates, 2013). Furthermore, differences in cranial anatomy between these taxa were identified as functionally relevant by Koyabu and Endo (2009) and contextualized in light of durophagy. Within this framework, the similarities in tori development between *Co. guereza* and *Co. polykomos* are perplexing and partly call into question the usefulness of the symphysis in dietary functional interpretations, at least for colobines. Phylogenetic inertia could potentially explain this result, as mitochondrial data provide a relatively young age (2.0 - 1.2 Ma) for the *Co. guereza* / *Co. polykomos* divergence (Ting, 2008). Further comparative phylogenetic analyses should complement our study by focusing specifically on the *Colobus* lineage to address this hypothesis.

Although *Co. polykomos* and *Piliocolobus badius* show similar symphyseal robustness (width of the symphysis/depth of the symphysis) according to Daegling and McGraw (2001) and Koyabu and Endo (2009), our study brings a subtler view by demonstrating that the relative breadth of the tori is distinct between these two taxa despite similar robustness values. Indeed, the STT is enlarged in *Colobus* while the ITT is enlarged in *Piliocolobus*. Furthermore, *Piliocolobus* is unique among the colobines in showing a markedly inclined symphysis, with values more in line with that of cercopithecines than that of other colobines. The reverse is true for the steep symphyseal profile of *Nasalis*, *Simias*, and *Rhinopithecus*. An inclined symphysis is hypothesized to reduce stress at the STT more efficiently than a steep symphysis (Hylander, 1984; Panagiotopoulou and Cobb, 2011). In this context, and under the model specified by Hylander (1984), more stresses are directed at the ITT in inclined symphysis. While the relatively robust ITT and inclined symphysis of *Piliocolobus* are in agreement with Hylander’s (1984) hypothesis, it fails to interpret the combination of a steeply inclined symphysis and robust ITT in *Nasalis*.

The symphyseal anatomy of *Nasalis* and *Colobus* can easily be distinguished using our morphometric protocol. Precisely, the *Nasalis* symphysis is steeply inclined, with a shorter planum alveolare, a shorter STT, and a larger ITT than in *Colobus* (Figure 9 and Table 9). Previous ecomorphological analyses of the colobine mandible have limited their comparison between the sympatric African colobines *Co. polykomos*, *Pi. badius* and *Pr. verus* (Daegling and McGraw, 2001; McGraw and Daegling, 2020). A *Nasalis* / *Colobus* comparison has the potential to emphasize the functional values of symphyseal traits, and ultimately address hypotheses regarding the selective pressures that shaped the symphysis of these taxa.

In this analysis, *Lophocebus* tends to differ from *Cercocebus* on the basis of its shorter planum alveolare, reduced STT, and enlarged ITT. Although not significant, these differences potentially underline useful functional considerations. Indeed, the two taxa are closely related and exhibit a durophagous diet but differ in their feeding behaviors (Daegling and McGraw, 2007; McGraw and Daegling, 2020). Bite forces generated during nut crushing are potentially higher at P_4_/M_1_ level in *Cercocebus,* as has been hypothesized based on its large premolars (Fleagle and McGraw, 1999, 2002), whereas *Lophocebus* uses its anterior dentition more frequently to process hard food (McGraw and Daegling, 2020). In this aspect, the relatively enlarged ITT of *Lophocebus* is consistent with greater resistance of its symphysis to coronal bending and dorsoventral shear generated by incisal biting (Hylander, 1984; Panagiotopoulou and Cobb, 2011). Differences in mandibular anatomy between *Lophocebus* and *Cercocebus* have been studied previously using alternative estimates of symphyseal shape (i.e., symphysis depth, symphysis breadth at the STT and symphysis size) in Daegling and McGraw (2007), but this did not demonstrate differences in symphyseal shape between both taxa. Our analysis provides a complementary view and point to the importance of the ITT to differentiates *Lophocebus* from *Cercocebus* on functional and taxonomic grounds.

### Taxonomic considerations

The generic status of *Procolobus* and *Piliocolobus* is debated and they are regarded either as a subgenus of *Procolobus* (Kuhn, 1967; Grubb et al., 2003; Grubb and Groves, 2013) or as separate genera (Groves, 2007). Except from symphyseal inclination, *Pr. verus* and *Pi. badius* are broadly comparable in symphyseal morphology. However, they differ markedly in the level of sexual dimorphism of symphyseal traits. In fact, *Pr. verus* shows strong sexual dimorphism in relative length of planum alveolare, relative development of the transverse tori, and symphyseal inclination, whereas no differences are observed between males and females of *Pi. badius*. This observation supports the distinction between *Procolobus* and *Piliocolobus* at the genus level, as advocated by Groves (2007), but should be further tested by including other *Piliocolobus* species, and in particular species from eastern and central Africa which are mostly absent from this study.

A sister taxon relationship has been established between *Trachypithecus* and *Presbytis* (Strasser and Delson, 1987; Sterner et al., 2006; Liedigk et al., 2012), while others considered *Presbytis* to be the sister taxon to all Asian colobines (Osterholz et al., 2008; Perelman et al., 2011), or more commonly as part of a monophyletic group including *Trachypithecus*, *Presbytis*, and *Semnopithecus*, with a sister taxon relationship between *Trachypithecus* and *Semnopithecus* (Brandon-Jones, 1984). *Presbytis* is unique within Presbytini in having a relatively small ITT and a long planum alveolare. It is also convergent with *Colobus* in its diminutive ITT. Symphyseal anatomy does not permit to justify a close phylogenetic relationship between *Presbytis* and *Trachypithecus* nor between *Presbytis* and *Semnopithecus*. An alternative hypothesis would be to explain the specificity of *Presbytis* on functional rather than phylogenetic grounds. Future analyses focused on evolutionary modeling should address this question and test whether the symphysis of *Presbytis* is the result of a distinct selective regime and autapomorphies or an expression of the plesiomorphic condition of Asian colobines.

With a relatively more developed ITT, *Allenopithecus nigroviridis* exhibits a relative breadth of the transverse tori quite distinct from other Cercopithecini (except for *Chlorocebus*; Figure 4 and Table 9). The phylogenetic relationship of *Allenopithecus* within the Cercopithecini is ambiguous, and the genus is regarded as plesiomorphic in its dentition (molar flare) and soft tissue (continuous ischial callosities in males), and also presents a specific karyotype compared to other Cercopithecini (Gautier-Hion, 2013; Lo Bianco et al., 2017). Its developed ITT is convergent with *Lophocebus* and *Theropithecus* and is another morphological character that supports its distinction from the Cercopithecini group. Interestingly, *Chlorocebus* has similar index values of relative planum alveolare length and transverse tori development with *Allenopithecus*. This result may illustrate symplesiomorphies or functional convergences, as both monkeys are omnivorous and opportunistic in their food selection (Butynski and Kingdon, 2013; Gautier-Hion, 2013). By demonstrating the distinctiveness of *Allenopithecus*, our finding is in accordance with the molecular phylogeny of Perelman et al. (2011) and Kuderna et al. (2023) inferred from nuclear sequences that place *Allenopithecus* as a basal member of the *Chlorocebus*/*Erythrocebus*/*Allochrocebus* clade.

In addition to *Allenopithecus*, *Miopithecus* was also considered as a plesiomorphic guenon, possibly the most basal member of the group according to the mitochondrial data of Guschanski et al. (2013) or the sister taxon of *Cercopithecus* following the nuclear data of Perelman et al. (2011). Our symphyseal ratios do not significantly distinguish *Miopithecus* from other guenons. However, our sample of *Miopithecus* is quite small and the inclusion of additional specimens could refine our results. Nonetheless, *Miopithecus* mean values for the transverse tori relative breadth and relative planum alveolare length are closer to those of *Cercopithecus* than those of *Allenopithecus*, demonstrating further phenetic differences between *Miopithecus* and *Allenopithecus* and favoring the topology of Perelman et al. (2011).

*Theropithecus gelada* is considered as the sister taxon to the *Papio*/*Lophocebus* clade among the Papionina (Perelman et al., 2011; Gilbert, 2013; Pugh and Gilbert, 2018), and is unique among cercopithecid by having a highly specialized grass-focused diet (Dunbar and Bose, 1991). *Theropithecus* can be differentiated from other papionines by presenting a steeply inclined symphysis and a well-developed ITT (also seen in *Lophocebus*; Figure 9 and Table 9). These symphyseal characteristics are in addition to the autapomorphic cranial and dental traits already documented in *Th. gelada*, and more broadly in the *Theropithecus* fossil record and lineage (Eck and Jablonski, 1984; Delson and Dean, 1993). The *Theropithecus* symphysis has been previously described as well buttressed by Dechow and Singer (1984) and Jolly (1972), with a weakly developed STT and a more strongly developed ITT (Delson and Dean, 1993) on the basis of a qualitative evaluation of the symphysis shape. Other from the demonstration of a long symphysis relative to cheek tooth length (Eck and Jablonski, 1984), no quantitative assessment and comparison of the symphyseal anatomy of extant and fossil *Theropithecus* has been undertaken to date. Here, we clarify the symphyseal anatomy of *Th. gelada* and provide adequate quantitative data for future comparisons with fossil *Theropithecus*. We also confirm the observations of Delson and Dean (1993). The ITT of *Th. gelada* is relatively enlarged compared to its STT and its STT is also shallower than that of similar-sized *Papio*, as previously assumed by Delson and Dean (1993). The differences we raised here between *Theropithecus* and *Papio* are particularly relevant given the taxonomic debate surrounding basal members of the group such as *Theropithecus baringensis*, which is viewed either as a basal member of the *Papio* lineage according to Iwamoto (1982), or of the *Theropithecus* lineage by Eck and Jablonski (1984). Similar questions have been raised for *Soromandrillus quadratirostris* with Gilbert (2013) advocating placing this taxon in the *Mandrillus*/*Cercocebus* lineage, while others consider it to be *Papio* (Iwamoto, 1982), *Theropithecus* (Eck and Jablonski, 1984), or a plesiomorphic member of the *Papio* lineage (Delson and Dean, 1993). We have demonstrated that the *Mandrillus* symphysis is characterized by a relatively long planum alveolare and a markedly inclined symphysis compared to *Papio* and *Th. gelada*. Although the *Mandrillus* sample presented here is unbalanced in terms of sex representativeness with an excess of male specimens, most of the *Mandrillus* traits used in phylogenetic analyses were recognized as dimorphic, with synapomorphies more readily identifiable in male specimens (Gilbert, 2013). Therefore, we advocate the inclusion of symphyseal traits in future comparative or phylogenetic analyses of fossil papionins that aims at identifying *Papio* or *Mandrillus* representatives.

*Macaca* is considered a primitive papionin with respect to African Papionina (Gilbert, 2013; Pugh and Gilbert, 2018). By showing the specificity of the symphysis of *Macaca* relative to *Papio*, and specifically its reduced ITT, long planum alveolare and inclined symphysis, we complement and corroborate previous qualitative observations drawn from phylogenetic analyses and comparative anatomy (Frost, 2001; Harrison, 2011; Gilbert, 2013). However, these analyses were limited to qualitative assessment of the inclination of the symphysis (Frost, 2001; Gilbert, 2013), or quantitative with limited comparative evidence (Harrison, 2011). The range of variation presented here for symphyseal traits will hence benefit to future phylogenetic and comparative analysis of fossil papionines by providing continuous data for traits previously treated as discrete. This is particularly true for the uncertainty that surrounds the phylogenetic relationship of the fossil papionin *Parapapio* and *Paradolichopithecus* (Leakey et al., 2003; Harrison, 2011; Gilbert, 2013; Kostopoulos et al., 2018; Pugh and Gilbert, 2018; Le Maître et al., 2023). Indeed, *Parapapio* is regarded by some authors as a paraphyletic genus, with notable criticisms concerning the taxonomic validity of its oldest representative, *Parapapio lothagamensis* (Harrison, 2011; Gilbert, 2013). The affinities of *Paradolichopithecus* to the Macacina or Papionina subtribes have not been fully demonstrated, despite compelling evidence for an affinity to Papionina from its cranial, mandibular, dental, and inner ear anatomy (Kostopoulos et al., 2018; Le Maître et al., 2023). These taxonomic debates will greatly benefit from the inclusion of the diagnostic quantitative symphyseal traits of *Papio* and *Macaca* presented in this study.

### Sexual dimorphism and symphyseal anatomy

Sexual dimorphism in symphyseal shape is markedly expressed in *Pr. verus* and *N. larvatus*. This result is expected as *N. larvatus* is considered one of the most dimorphic colobine species in terms of body mass and canine dimensions (Plavcan and Van Schaik, 1992; Plavcan, 2001). *Pr. verus* is moderately dimorphic for body mass, but nevertheless strongly dimorphic for canine dimensions (Leutenegger, 1982; Plavcan and Van Schaik, 1992; Plavcan, 2001). *Presbytis bicolor* (formerly a subspecies of *Presbytis melalophos*, but later proposed by Meyer et al. (2011) as a distinct species) does not show significant differences in transverse tori development, relative length of the planum alveolare, and symphyseal inclination between males and females. *Presbytis* is moderately dimorphic in both body mass and canine dimensions, and dimorphism in canine dimensions does not vary markedly between *Presbytis* species with different levels of sexual dimorphism (e.g., *Pre. potenziani* and *Pre. melalophos* in Plavcan and Van Schaik (1992)). The absence of dimorphism in transverse tori breadth, planum alveolare and symphyseal inclination of *Pre. bicolor* is thus in accordance with the low dimorphism observed in body mass and canine dimensions of the *Presbytis* genus.

Whereas sexual dimorphism is detected in the relative transverse tori breadth and planum alveolare length of *Co. polykomos*, no dimorphism is observed in *Co. guereza*. The latter species shows slightly less dimorphism in lower canine dimensions (Plavcan and Van Schaik, 1992), although substantial variation in canine size dimorphism is observed between western and eastern *Co. guereza* subspecies (Hayes et al., 1995). Our sample is not homogeneous and includes a mixture of western (i.e., *Co. guereza occidentalis*) and eastern (*Co. guereza kikuyensis*) specimens, but it confirms the presence of a low level of dimorphism in canine dimensions concomitant with a low level of dimorphism in symphyseal traits.

As expected, the symphysis of *P. anubis* is extremely dimorphic, a result consistent with the marked dimorphism observed on its canine dimensions and body mass (Plavcan and Van Schaik, 1992). Overall, it is tempting to link canine dimensions to sexual dimorphism in tori development, relative planum alveolare length and symphysis inclination. Fukase (2011) showed that the enlarged canine size and prolonged canine development of male *Papio hamadryas* influenced corpus depth and symphysis width. In contrast, the less dimorphic *Macaca mulatta* did not express dimorphism in corpus depth and symphysis width (Fukase, 2011). By demonstrating sexual dimorphism in relative development of the transverse tori, planum alveolare length, and symphyseal inclination in *P. anubis*, but not in *Ma. fuscata* and *Ma. fascicularis*, our data support Fukase’s (2011) hypothesis that canine dimensions significantly influence symphyseal shape. Although this hypothesis is in agreement with our data for the papionin sample, it is less convincing for colobines. Indeed, *Pi. badius* is documented as a highly dimorphic species in canine dimensions (Plavcan and Van Schaik, 1992), but we did not detect significant differences in symphyseal traits between male and female *Pi. badius*. In contrast, *Pr. verus* is highly dimorphic in symphyseal shape despite having a similar degree of dimorphism in occlusal dimensions of the lower canine as that of *Pi. badius* Plavcan and Van Schaik (1992). Subocclusal anatomy, specifically the dimensions and inclination of the canine crown and root, may be a better explanation for the lack of dimorphism in *Pi. badius* and its presence in *Pr. verus*. Future studies should address differences in canine root shape and ontogeny to validate this hypothesis. In addition, incisor proclination, root length and ontogeny should not be neglected in this problematic despite a lack of data on the incisors subocclusal anatomy.

In the context of the functional adaptations of the symphysis to the biomechanics of mastication (e.g., STT breadth and wishboning), the confounding effect of sexual dimorphism in symphysis shape demonstrates that the functional and adaptive interpretation of this anatomical part is far from simple. Rather, this study demonstrates that the symphysis responds to a dynamic and complex interplay of allometry, sexual dimorphism, phylogenetic inertia and adaptations.

## CONCLUSION

Although routinely used in the diagnosis of fossil and extant cercopithecids, a quantitative assessment of the symphysis of cercopithecids was lacking. Based on a large dataset that includes 756 specimens from 22 cercopithecid genera, we explored the taxonomic value of the breadth of the transverse tori, length of the planum alveolare, and symphyseal inclination using univariate and multivariate data as well as comparative phylogenetic methods. We also assessed the source of variation in these traits by examining sexual dimorphism and allometry. A clear phylogenetic signal is embedded in the symphyseal anatomy and statistically significant differences can be found between cercopithecid subfamilies, tribes, subtribes, and genera. Specifically, the symphysis of cercopithecines is characterized by a marked inclination, a reduced inferior transverse torus, and a long planum alveolare compared to colobines. Exceptions exist within each subfamily and our data underline the distinct symphyseal anatomy of *Colobus*, *Presbytis*, *Allenopithecus*, *Lophocebus*, *Theropithecus* and *Macaca*. Sexual dimorphism is well expressed in the symphyseal anatomy of several colobines (e.g., *Procolobus verus*) and cercopithecines (e.g., *Papio anubis*) taxa. Although the proximate cause of this dimorphism is largely indeterminate for colobine taxa, the subocclusal anatomy of incisors and canines is a prime candidate for explaining the shape differences between male and female specimens. By providing a range of variation and highlighting sound phenetic differences in symphyseal traits among cercopithecids, our study will greatly benefit future phylogenetic analyses, fossil specimen description, and ecomorphological analyses that aim to identify osteological correlates of diet and/or feeding behaviors.

## Supporting information

Supplementary Informations

## ACKNOWLEDGEMENTS

The authors express their gratitude to A. van Heteren, M. Hiermeier, and T. Nishimura for giving the opportunity to study the skeletal collections from the Bavarian State Collection of Zoology and Center for the Evolutionary Origins of Human Behavior (Kyoto University) under their care. We also thank the members of the osteological collection of the NMK for their guidance and access to the collection. We are grateful to Emmanuel Gilissen and Mathys Rotonda for access to study the skeletal collection from the Royal Museum for Central Africa (Tervuren). We especially acknowledge the NSF BCS 155248 attributed to D. Boyer. We appreciate the Domestic Research Program of Ryokoku University. This research was financially supported by the Japan Society for the Promotion of Science Kakenhi (23H02562 and 16H02757) to MN.

