## Supplementary Informations for "Anatomy of the mandibular symphysis of extant cercopithecids: taxonomy and variation"

SOM Figure S1: Linear regression of the index PAL/GM on PAL/SL.

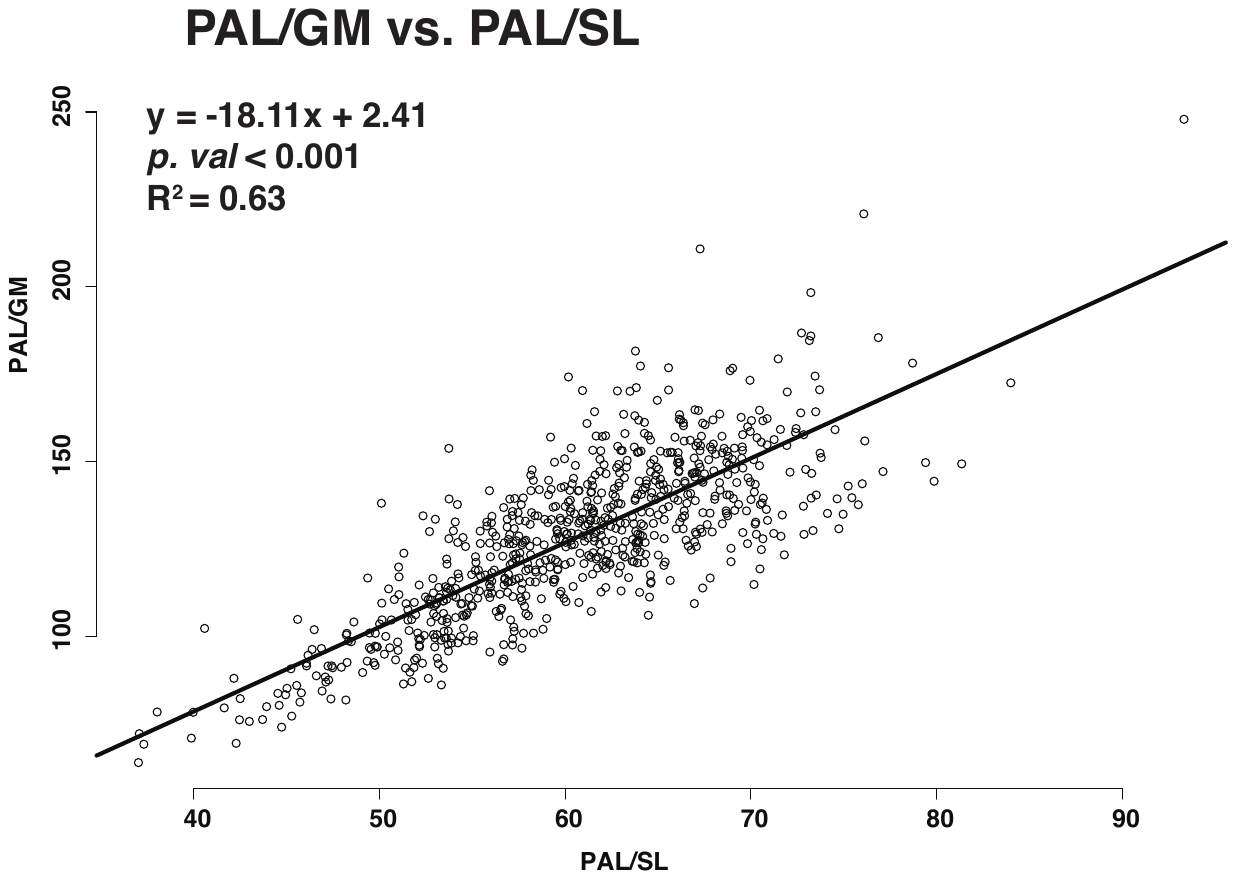

SOM Figure S2: Boxplots of A) planum alveolare length divided by symphyseal length, and B) planum alveolare length divided by GM. Boxplots with mean (losange), median (vertical bar), first and third quartiles.

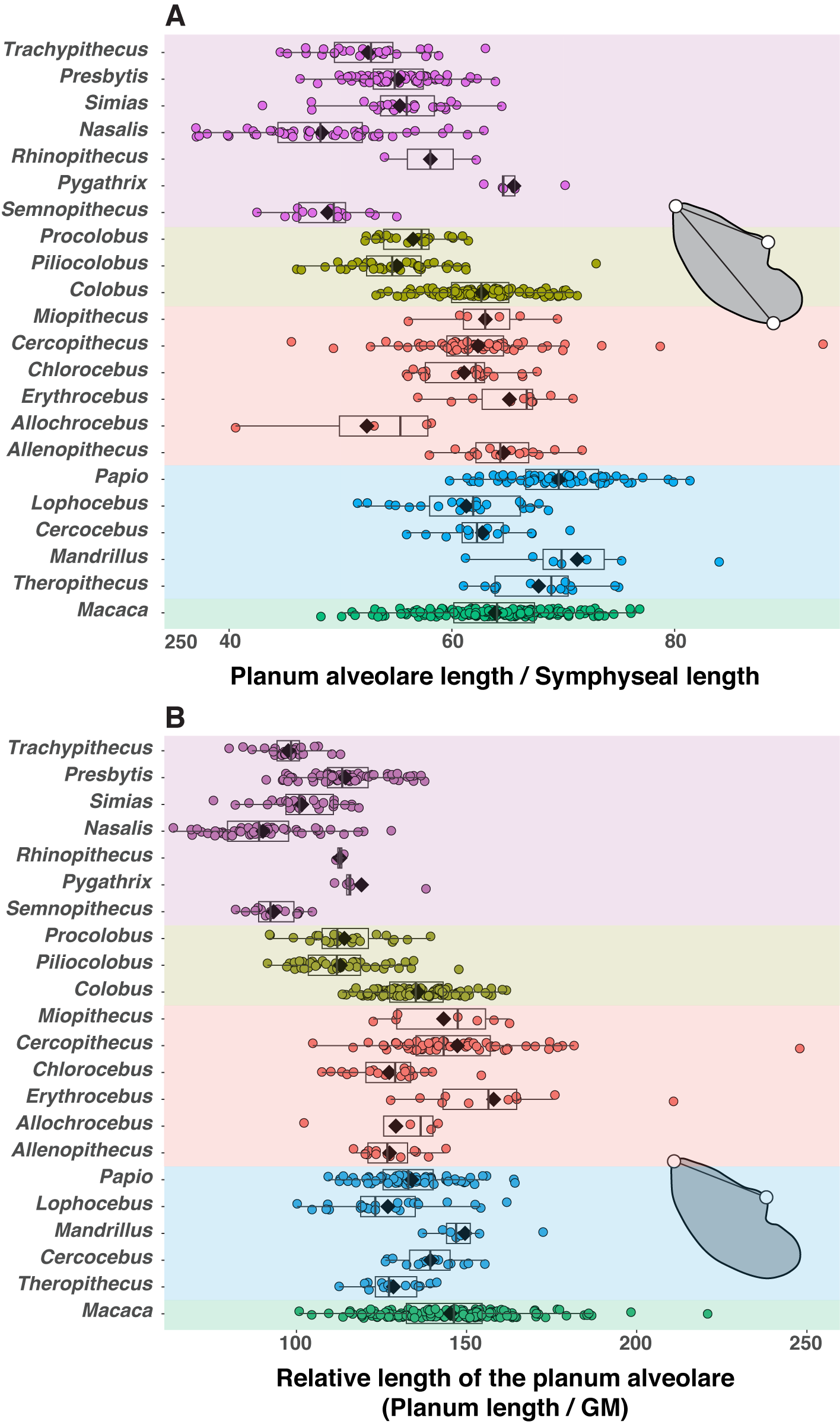

SOM Figure S3: Partial least squares regression of the natural logarithm of geometric means on the natural logarithm of body mass.

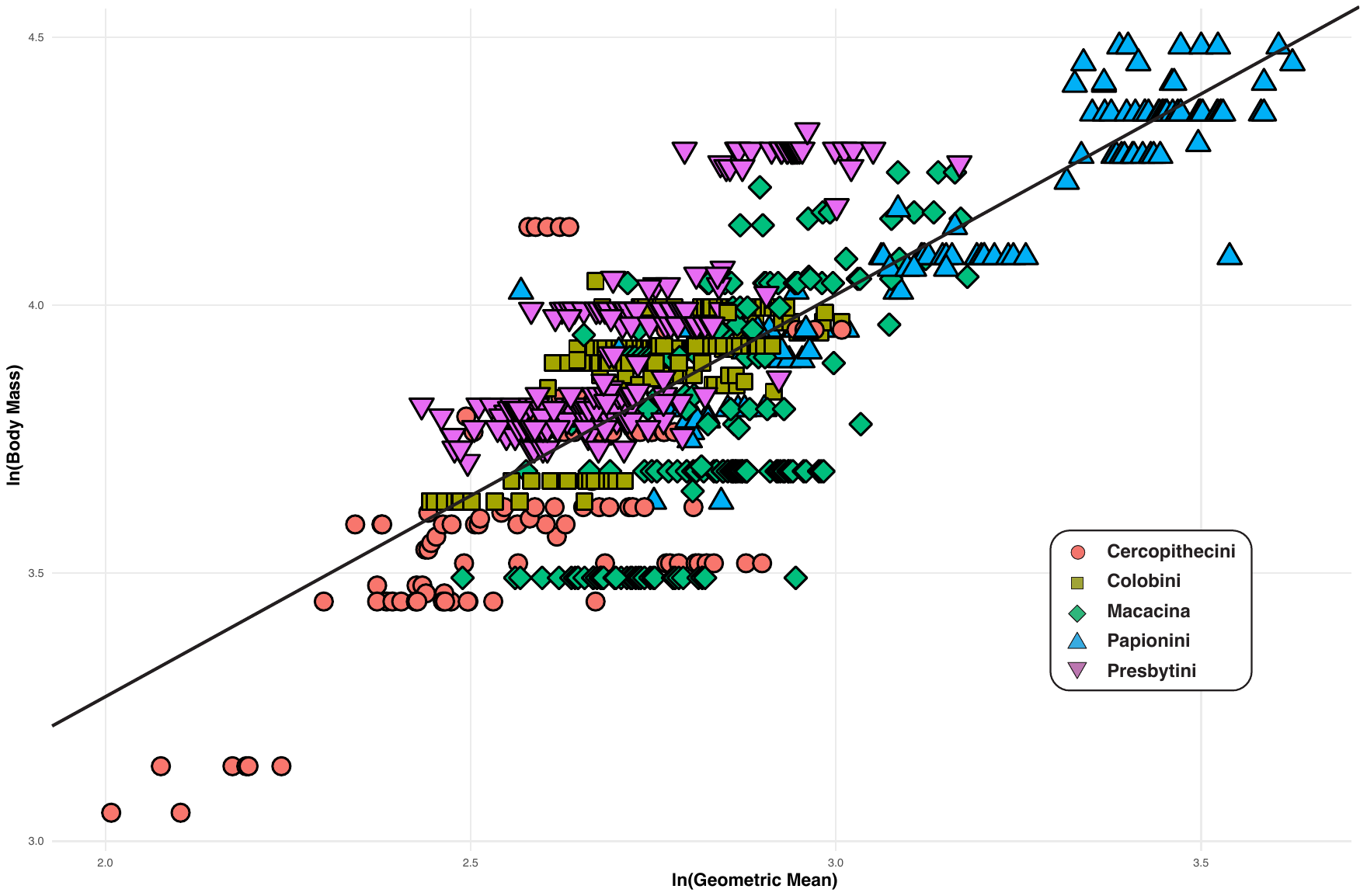

| **Taxa** | **Index** | **Mean and standard deviation** |
| --- | --- | --- |
| Colobinae | PLA/SL | 56.1 ± 6.6 |
| Cercopithecinae |  | 64.4 ± 6.1 |
| Cercopithecini |  | 62.4 ± 6.0 |
| Papionini |  | 65.2 ± 6.0 |
| Colobini |  | 59.9 ± 5.4 |
| Presbytini |  | 52.9 ± 5.9 |
| Macacina |  | 63.9 ± 5.7 |
| Papionina |  | 67.1 ± 5.9 |

SOM Table S1: Descriptive statistics (mean and standard deviation) for the index of planum alveolare length divided by symphyseal length at higher taxonomic level

SOM Table S2: Descriptive statistics (mean and standard deviation) for the index of planum alveolare length divided by symphyseal length at genus level

| **Taxa** | **Index** | **Mean and standard deviation** |
| --- | --- | --- |
| *Allenopithecus* | PLA/SL | 64.6 ± 3.6 |
| *Allochrocebus* |  | 52.4 ± 8.2 |
| *Cercocebus* |  | 62.7 ± 3.9 |
| *Cercopithecus* |  | 62.3 ± 6.8 |
| *Chlorocebus* |  | 61.1 ± 3.6 |
| *Colobus* |  | 62.7 ± 4.2 |
| *Erythrocebus* |  | 65.2 ± 4.3 |
| *Lophocebus* |  | 61.3 ± 5.0 |
| *Macaca* |  | 63.9 ± 5.6 |
| *Mandrillus* |  | 71.3 ± 7.1 |
| *Miopithecus* |  | 63.0 ± 4.3 |
| *Nasalis* |  | 48.3 ± 6.3 |
| *Papio* |  | 69.6 ± 4.7 |
| *Piliocolobus* |  | 55.0 ± 4.7 |
| *Presbytis* |  | 55.2 ± 3.4 |
| *Procolobus* |  | 56.5 ± 2.8 |
| *Pygathrix* |  | 65.6 ± 2.7 |
| *Rhinopithecus* |  | 58.1 ± 5.8 |
| *Semnopithecus* |  | 48.8 ± 3.4 |
| *Simias* |  | 55.3 ± 4.6 |
| *Theropithecus* |  | 67.8 ± 4.3 |
| *Trachypithecus* |  | 52.5 ± 4.5 |

| Genus | Species | Subspecies | Subfamily | Tribe | Accession Number | Sex | Provenance of the 3D model |
| --- | --- | --- | --- | --- | --- | --- | --- |
| *Allenopithecus* | *nigroviridis* |  | Cercopithecinae | Cercopithecini | RMCA 5613 | Male | https://www.morphosource.org |
| *Allenopithecus* | *nigroviridis* |  | Cercopithecinae | Cercopithecini | RMCA 7756M1 | Female |  |
| *Allenopithecus* | *nigroviridis* |  | Cercopithecinae | Cercopithecini | RMCA 9339 | Female |  |
| *Allenopithecus* | *nigroviridis* |  | Cercopithecinae | Cercopithecini | RMCA 10303 | Female |  |
| *Allenopithecus* | *nigroviridis* |  | Cercopithecinae | Cercopithecini | RMCA 18971 | Male |  |
| *Allenopithecus* | *nigroviridis* |  | Cercopithecinae | Cercopithecini | RMCA 23496 | Male |  |
| *Allenopithecus* | *nigroviridis* |  | Cercopithecinae | Cercopithecini | RMCA 28645 | Male |  |
| *Allenopithecus* | *nigroviridis* |  | Cercopithecinae | Cercopithecini | RMCA 28647 | Male |  |
| *Allenopithecus* | *nigroviridis* |  | Cercopithecinae | Cercopithecini | RMCA 27993 | Male |  |
| *Allenopithecus* | *nigroviridis* |  | Cercopithecinae | Cercopithecini | RMCA 27994 | Male |  |
| *Allenopithecus* | *nigroviridis* |  | Cercopithecinae | Cercopithecini | RMCA 28649 | Male |  |
| *Allenopithecus* | *nigroviridis* |  | Cercopithecinae | Cercopithecini | RMCA 28698 | Male |  |
| *Allenopithecus* | *nigroviridis* |  | Cercopithecinae | Cercopithecini | RMCA 28769 | Male |  |
| *Allenopithecus* | *nigroviridis* |  | Cercopithecinae | Cercopithecini | RMCA 28770 | Male |  |
| *Allenopithecus* | *nigroviridis* |  | Cercopithecinae | Cercopithecini | RMCA 28771 | Female |  |
| *Allochrocebus* | *lhoesti* |  | Cercopithecinae | Cercopithecini | KUPRI 11560 | Male | This study |
| *Allochrocebus* | *lhoesti* |  | Cercopithecinae | Cercopithecini | KUPRI 11561 | Female |  |
| *Allochrocebus* | *preussi* |  | Cercopithecinae | Cercopithecini | BSZM 189419 | Male |  |
| *Allochrocebus* | *preussi* |  | Cercopithecinae | Cercopithecini | BSZM AP163 | Female |  |
| *Cercocebus* | *galeritus* |  | Cercopithecinae | Papionina | AMNH 52634 | Male | https://www.morphosource.org |
| *Cercocebus* | *agilis* |  | Cercopithecinae | Papionina | AMNH 52640 | Female |  |
| *Cercocebus* | *agilis* |  | Cercopithecinae | Papionina | AMNH 52645 | Male |  |
| *Cercocebus* | *agilis* |  | Cercopithecinae | Papionina | AMNH 52648 | Male |  |
| *Cercocebus* | *galeritus* |  | Cercopithecinae | Papionina | AMNH 70063 | Male |  |
| *Cercocebus* | *torquatus* |  | Cercopithecinae | Papionina | MCZ 18612 | Female |  |
| *Cercocebus* | *torquatus* |  | Cercopithecinae | Papionina | MCZ 19184 | Male |  |
| *Cercocebus* | *torquatus* |  | Cercopithecinae | Papionina | MCZ 19982 | Male |  |
| *Cercocebus* | *atys* |  | Cercopithecinae | Papionina | MCZ 25626 | Male |  |
| *Cercocebus* | *atys* |  | Cercopithecinae | Papionina | RMCA 07M17 | ? |  |
| *Cercocebus* | *atys* |  | Cercopithecinae | Papionina | RMCA 07M27 | ? |  |
| *Cercocebus* | *galeritus* |  | Cercopithecinae | Papionina | RMCA 986 | ? |  |
| *Cercocebus* | *galeritus* |  | Cercopithecinae | Papionina | RMCA 8373 | Female |  |
| *Cercocebus* | *galeritus* |  | Cercopithecinae | Papionina | KUPRI 7115 | Male |  |
| *Cercopithecus* | *mitis* |  | Cercopithecinae | Cercopithecini | RMCA 1842 | Male |  |
| *Cercopithecus* | *mitis* |  | Cercopithecinae | Cercopithecini | RMCA 2272 | Female |  |
| *Cercopithecus* | *mitis* |  | Cercopithecinae | Cercopithecini | RMCA 2275 | Female |  |
| *Cercopithecus* | *mitis* |  | Cercopithecinae | Cercopithecini | RMCA 2796 | Male |  |
| *Cercopithecus* | *mitis* |  | Cercopithecinae | Cercopithecini | RMCA 3311 | Female |  |
| *Cercopithecus* | *mitis* |  | Cercopithecinae | Cercopithecini | RMCA 3312 | Female |  |
| *Cercopithecus* | *mitis* |  | Cercopithecinae | Cercopithecini | RMCA 5994 | Female |  |
| *Cercopithecus* | *mitis* |  | Cercopithecinae | Cercopithecini | RMCA 6135 | Male |  |
| *Cercopithecus* | *mitis* |  | Cercopithecinae | Cercopithecini | RMCA 6136 | Male |  |
| *Cercopithecus* | *mitis* |  | Cercopithecinae | Cercopithecini | RMCA 6140 | Male |  |
| *Cercopithecus* | *mitis* |  | Cercopithecinae | Cercopithecini | RMCA 9835 | Male |  |
| *Cercopithecus* | *mitis* |  | Cercopithecinae | Cercopithecini | RMCA 12346 | Male |  |
| *Cercopithecus* | *mitis* |  | Cercopithecinae | Cercopithecini | RMCA 12347 | Female |  |
| *Cercopithecus* | *mitis* |  | Cercopithecinae | Cercopithecini | RMCA 12348 | Female |  |
| *Cercopithecus* | *mitis* |  | Cercopithecinae | Cercopithecini | RMCA 12736 | Male |  |
| *Cercopithecus* | *mitis* |  | Cercopithecinae | Cercopithecini | RMCA 13115 | Male |  |
| *Cercopithecus* | *mitis* |  | Cercopithecinae | Cercopithecini | RMCA 13124 | Male |  |
| *Cercopithecus* | *mitis* |  | Cercopithecinae | Cercopithecini | RMCA 15242 | Male |  |
| *Cercopithecus* | *mitis* |  | Cercopithecinae | Cercopithecini | RMCA 16735 | Male |  |
| *Cercopithecus* | *mitis* |  | Cercopithecinae | Cercopithecini | RMCA 17918 | Male |  |
| *Cercopithecus* | *mitis* |  | Cercopithecinae | Cercopithecini | RMCA 18899 | Male |  |
| *Cercopithecus* | *mitis* |  | Cercopithecinae | Cercopithecini | RMCA 19887 | Male |  |
| *Cercopithecus* | *mitis* |  | Cercopithecinae | Cercopithecini | RMCA 20012 | Male |  |
| *Cercopithecus* | *mitis* |  | Cercopithecinae | Cercopithecini | RMCA 25450 | Male |  |
| *Cercopithecus* | *mitis* |  | Cercopithecinae | Cercopithecini | RMCA 25451 | Male |  |
| *Cercopithecus* | *mitis* |  | Cercopithecinae | Cercopithecini | RMCA 25489 | Male |  |
| *Cercopithecus* | *mitis* |  | Cercopithecinae | Cercopithecini | RMCA 26173 | Male |  |
| *Cercopithecus* | *mitis* |  | Cercopithecinae | Cercopithecini | RMCA 28005 | Male |  |
| *Cercopithecus* | *mitis* |  | Cercopithecinae | Cercopithecini | RMCA 35118 | Male |  |
| *Cercopithecus* | *mitis* |  | Cercopithecinae | Cercopithecini | RMCA 37538 | Male |  |
| *Cercopithecus* | *mitis* |  | Cercopithecinae | Cercopithecini | RMCA 37530 | Female |  |
| *Cercopithecus* | *mitis* |  | Cercopithecinae | Cercopithecini | RMCA 37531 | Female |  |
| *Cercopithecus* | *mitis* |  | Cercopithecinae | Cercopithecini | RMCA 37540 | Female |  |
| *Cercopithecus* | *albogularis* |  | Cercopithecinae | Cercopithecini | USNM 452574 | Male |  |
| *Cercopithecus* | *albogularis* |  | Cercopithecinae | Cercopithecini | USNM 452581 | Female |  |
| *Cercopithecus* | *ascanius* |  | Cercopithecinae | Cercopithecini | SBU-RS 81-5 | Male |  |
| *Cercopithecus* | *ascanius* |  | Cercopithecinae | Cercopithecini | SBU-RS 81-6 | Female |  |
| *Cercopithecus* | *ascanius* |  | Cercopithecinae | Cercopithecini | SBU-RS 81-11 | Male |  |
| *Cercopithecus* | *ascanius* |  | Cercopithecinae | Cercopithecini | SBU-RS 81-13 | Female |  |
| *Cercopithecus* | *ascanius* |  | Cercopithecinae | Cercopithecini | USNM 182355 | Female |  |
| *Cercopithecus* | *cephus* |  | Cercopithecinae | Cercopithecini | SBU-RS 85-5 | Female |  |
| *Cercopithecus* | *cephus* |  | Cercopithecinae | Cercopithecini | SBU-RS 85-8 | Female |  |
| *Cercopithecus* | *mona* |  | Cercopithecinae | Cercopithecini | USNM 480920 | ? |  |
| *Cercopithecus* | *mona* |  | Cercopithecinae | Cercopithecini | USNM 481007 | ? |  |
| *Cercopithecus* | *mona* |  | Cercopithecinae | Cercopithecini | SBU-RS 93-1 | ? |  |
| *Cercopithecus* | *neglectus* |  | Cercopithecinae | Cercopithecini | SBU-RS 81-9 | Male |  |
| *Cercopithecus* | *neglectus* |  | Cercopithecinae | Cercopithecini | SBU-RS 85-3 | Male |  |
| *Cercopithecus* | *nictitans* |  | Cercopithecinae | Cercopithecini | USNM 220377 | Male |  |
| *Cercopithecus* | *nictitans* |  | Cercopithecinae | Cercopithecini | USNM 480838 | ? |  |
| *Cercopithecus* | *nictitans* |  | Cercopithecinae | Cercopithecini | USNM 481770 | Male |  |
| *Cercopithecus* | *nictitans* |  | Cercopithecinae | Cercopithecini | USNM 537776 | Female |  |
| *Cercopithecus* | *nictitans* |  | Cercopithecinae | Cercopithecini | SBU-RS 85-9 | Male |  |
| *Cercopithecus* | *petaurista* |  | Cercopithecinae | Cercopithecini | USNM 435021 | Male |  |
| *Cercopithecus* | *petaurista* |  | Cercopithecinae | Cercopithecini | USNM 477317 | Female |  |
| *Cercopithecus* | *petaurista* |  | Cercopithecinae | Cercopithecini | USNM 481778 | Female |  |
| *Cercopithecus* | *petaurista* |  | Cercopithecinae | Cercopithecini | USNM 481779 | Male |  |
| *Chlorocebus* | *aethiops* |  | Cercopithecinae | Cercopithecini | KUPRI 1789 | Male | This study |
| *Chlorocebus* | *aethiops* |  | Cercopithecinae | Cercopithecini | KUPRI 1863 | Male |  |
| *Chlorocebus* | *aethiops* |  | Cercopithecinae | Cercopithecini | KUPRI 5470 | Male |  |
| *Chlorocebus* | *aethiops* |  | Cercopithecinae | Cercopithecini | KUPRI 5995 | Male |  |
| *Chlorocebus* | *aethiops* |  | Cercopithecinae | Cercopithecini | KUPRI 5996 | Male |  |
| *Chlorocebus* | *aethiops* |  | Cercopithecinae | Cercopithecini | KUPRI 6000 | Male |  |
| *Chlorocebus* | *aethiops* |  | Cercopithecinae | Cercopithecini | KUPRI 6009 | Female |  |
| *Chlorocebus* | *aethiops* |  | Cercopithecinae | Cercopithecini | KUPRI 6010 | Female |  |
| *Chlorocebus* | *aethiops* |  | Cercopithecinae | Cercopithecini | KUPRI 6011 | Female |  |
| *Chlorocebus* | *aethiops* |  | Cercopithecinae | Cercopithecini | KUPRI 6012 | Female |  |
| *Chlorocebus* | *aethiops* |  | Cercopithecinae | Cercopithecini | KUPRI 6013 | Female |  |
| *Chlorocebus* | *aethiops* |  | Cercopithecinae | Cercopithecini | KUPRI 6014 | Female |  |
| *Chlorocebus* | *aethiops* |  | Cercopithecinae | Cercopithecini | KUPRI 6015 | Female |  |
| *Chlorocebus* | *aethiops* |  | Cercopithecinae | Cercopithecini | KUPRI 6024 | Female |  |
| *Chlorocebus* | *aethiops* |  | Cercopithecinae | Cercopithecini | KUPRI 6045 | Female |  |
| *Chlorocebus* | *aethiops* |  | Cercopithecinae | Cercopithecini | KUPRI 6050 | Female |  |
| *Chlorocebus* | *aethiops* |  | Cercopithecinae | Cercopithecini | KUPRI 6051 | Male |  |
| *Chlorocebus* | *aethiops* |  | Cercopithecinae | Cercopithecini | KUPRI 6727 | Male |  |
| *Chlorocebus* | *aethiops* |  | Cercopithecinae | Cercopithecini | KUPRI 6731 | Male |  |
| *Chlorocebus* | *aethiops* |  | Cercopithecinae | Cercopithecini | KUPRI 7113 | Male |  |
| *Colobus* | *caudatus* |  | Colobinae | Colobini | MCZ 21151 | Female | https://www.morphosource.org |
| *Colobus* | *angolensis* | *palliatus* | Colobinae | Colobini | MCZ 22624 | Female |  |
| *Colobus* | *angolensis* | *palliatus* | Colobinae | Colobini | MCZ 22626 | Female |  |
| *Colobus* | *angolensis* | *palliatus* | Colobinae | Colobini | MCZ 22629 | Female |  |
| *Colobus* | *polykomos* |  | Colobinae | Colobini | MCZ 463680 | Female |  |
| *Colobus* | *vellerosus* |  | Colobinae | Colobini | RMCA 7313M234 | Female |  |
| *Colobus* | *polykomos* |  | Colobinae | Colobini | RMCA 28851 | Male |  |
| *Colobus* | *polykomos* |  | Colobinae | Colobini | RMCA 28852 | Male |  |
| *Colobus* | *polykomos* |  | Colobinae | Colobini | RMCA 28853 | Male |  |
| *Colobus* | *polykomos* |  | Colobinae | Colobini | RMCA 28856 | Female |  |
| *Colobus* | *polykomos* |  | Colobinae | Colobini | RMCA 28857 | Female |  |
| *Colobus* | *polykomos* |  | Colobinae | Colobini | RMCA 28859 | Female |  |
| *Colobus* | *polykomos* |  | Colobinae | Colobini | RMCA 31491 | Male |  |
| *Colobus* | *polykomos* |  | Colobinae | Colobini | RMCA 35126 | Male |  |
| *Colobus* | *polykomos* |  | Colobinae | Colobini | KUPRI 1046 | Female | This study |
| *Colobus* | *polykomos* |  | Colobinae | Colobini | KUPRI 1067 | Female |  |
| *Colobus* | *polykomos* |  | Colobinae | Colobini | KUPRI 1071 | Female |  |
| *Colobus* | *polykomos* |  | Colobinae | Colobini | KUPRI 1085 | Female |  |
| *Colobus* | *polykomos* |  | Colobinae | Colobini | KUPRI 1087 | Female |  |
| *Colobus* | *polykomos* |  | Colobinae | Colobini | KUPRI 1103 | Female |  |
| *Colobus* | *polykomos* |  | Colobinae | Colobini | KUPRI 1105 | Female |  |
| *Colobus* | *polykomos* |  | Colobinae | Colobini | KUPRI 1121 | Female |  |
| *Colobus* | *polykomos* |  | Colobinae | Colobini | KUPRI 1127 | Female |  |
| *Colobus* | *polykomos* |  | Colobinae | Colobini | KUPRI 1053 | Male |  |
| *Colobus* | *polykomos* |  | Colobinae | Colobini | KUPRI 1078 | Male |  |
| *Colobus* | *polykomos* |  | Colobinae | Colobini | KUPRI 1086 | Male |  |
| *Colobus* | *polykomos* |  | Colobinae | Colobini | KUPRI 1115 | Male |  |
| *Colobus* | *polykomos* |  | Colobinae | Colobini | KUPRI 1118 | Male |  |
| *Colobus* | *polykomos* |  | Colobinae | Colobini | KUPRI 1135 | Male |  |
| *Colobus* | *polykomos* |  | Colobinae | Colobini | KUPRI 1136 | Male |  |
| *Colobus* | *polykomos* |  | Colobinae | Colobini | KUPRI 1137 | Male |  |
| *Colobus* | *angolensis* |  | Colobinae | Colobini | RMCA 5643 | Female | https://www.morphosource.org |
| *Colobus* | *angolensis* |  | Colobinae | Colobini | RMCA 8064 | Male |  |
| *Colobus* | *angolensis* |  | Colobinae | Colobini | RMCA 8130 | Male |  |
| *Colobus* | *angolensis* |  | Colobinae | Colobini | RMCA 9912 | Male |  |
| *Colobus* | *angolensis* |  | Colobinae | Colobini | SBU-RS 81-4 | Female |  |
| *Colobus* | *angolensis* |  | Colobinae | Colobini | SBU-RS 83-3 | Male |  |
| *Colobus* | *angolensis* |  | Colobinae | Colobini | SBU-RS 83-5 | Male |  |
| *Colobus* | *angolensis* |  | Colobinae | Colobini | SBU-RS 81-10 | Male |  |
| *Colobus* | *angolensis* |  | Colobinae | Colobini | SBU-RS 81-14 | Female |  |
| *Colobus* | *angolensis* |  | Colobinae | Colobini | KAS 177 | Female | This study |
| *Colobus* | *guereza* | *occidentalis* | Colobinae | Colobini | AMNH 52209 | Male | https://www.morphosource.org |
| *Colobus* | *guereza* | *occidentalis* | Colobinae | Colobini | AMNH 52210 | Male |  |
| *Colobus* | *guereza* | *occidentalis* | Colobinae | Colobini | AMNH 52223 | Female |  |
| *Colobus* | *guereza* | *occidentalis* | Colobinae | Colobini | AMNH 52237 | Male |  |
| *Colobus* | *guereza* | *occidentalis* | Colobinae | Colobini | AMNH 52238 | Female |  |
| *Colobus* | *guereza* | *occidentalis* | Colobinae | Colobini | AMNH 52241 | Female |  |
| *Colobus* | *guereza* | *occidentalis* | Colobinae | Colobini | AMNH 52245 | Female |  |
| *Colobus* | *guereza* | *kikuyensis* | Colobinae | Colobini | OM 3042 | Female | This study |
| *Colobus* | *guereza* | *kikuyensis* | Colobinae | Colobini | OM 3044 | Female |  |
| *Colobus* | *guereza* | *kikuyensis* | Colobinae | Colobini | OM 3045 | Female |  |
| *Colobus* | *guereza* | *kikuyensis* | Colobinae | Colobini | OM 3085 | Female |  |
| *Colobus* | *guereza* | *kikuyensis* | Colobinae | Colobini | OM 3097 | Male |  |
| *Colobus* | *guereza* | *kikuyensis* | Colobinae | Colobini | OM 3053 | Male |  |
| *Colobus* | *guereza* | *kikuyensis* | Colobinae | Colobini | OM 3125 | Male |  |
| *Colobus* | *guereza* |  | Colobinae | Colobini | KUPRI 11372 | Female |  |
| *Colobus* | *guereza* |  | Colobinae | Colobini | KUPRI 8673 | Male |  |
| *Colobus* | *satanas* |  | Colobinae | Colobini | BSZM 1914 263 | Male |  |
| *Colobus* | *polykomos* |  | Colobinae | Colobini | RMCA 8107M87 | Female | https://www.morphosource.org |
| *Colobus* | *polykomos* |  | Colobinae | Colobini | RMCA 8107M88 | Male |  |
| *Colobus* | *polykomos* |  | Colobinae | Colobini | RMCA 8107M89 | Male |  |
| *Colobus* | *polykomos* |  | Colobinae | Colobini | RMCA 8107M90 | Female |  |
| *Colobus* | *polykomos* |  | Colobinae | Colobini | RMCA 8107M91 | Male |  |
| *Colobus* | *polykomos* |  | Colobinae | Colobini | RMCA 8107M92 | Female |  |
| *Colobus* | *polykomos* |  | Colobinae | Colobini | RMCA 8107M93 | Male |  |
| *Colobus* | *polykomos* |  | Colobinae | Colobini | RMCA 8107M94 | Female |  |
| *Colobus* | *polykomos* |  | Colobinae | Colobini | RMCA 8107M95 | Female |  |
| *Colobus* | *polykomos* |  | Colobinae | Colobini | RMCA 8107M96 | Female |  |
| *Colobus* | *polykomos* |  | Colobinae | Colobini | RMCA 8107M99 | Female |  |
| *Colobus* | *polykomos* |  | Colobinae | Colobini | RMCA 8107M100 | Female |  |
| *Colobus* | *polykomos* |  | Colobinae | Colobini | RMCA 8107M101 | Female |  |
| *Colobus* | *polykomos* |  | Colobinae | Colobini | RMCA 8107M102 | Male |  |
| *Colobus* | *polykomos* |  | Colobinae | Colobini | RMCA 8107M103 | Male |  |
| *Colobus* | *polykomos* |  | Colobinae | Colobini | RMCA 8107M105 | Female |  |
| *Colobus* | *polykomos* |  | Colobinae | Colobini | RMCA 8107M106 | Female |  |
| *Colobus* | *polykomos* |  | Colobinae | Colobini | RMCA 8107M107 | Female |  |
| *Colobus* | *polykomos* |  | Colobinae | Colobini | RMCA 8107M108 | Male |  |
| *Colobus* | *polykomos* |  | Colobinae | Colobini | RMCA 8107M110 | Male |  |
| *Colobus* | *polykomos* |  | Colobinae | Colobini | RMCA 8107M120 | Female |  |
| *Colobus* | *polykomos* |  | Colobinae | Colobini | RMCA 8107M126 | Male |  |
| *Colobus* | *polykomos* |  | Colobinae | Colobini | RMCA 8107M127 | Female |  |
| *Colobus* | *polykomos* |  | Colobinae | Colobini | RMCA 8107M128 | Female |  |
| *Colobus* | *polykomos* |  | Colobinae | Colobini | RMCA 8107M129 | Male |  |
| *Colobus* | *polykomos* |  | Colobinae | Colobini | RMCA 8107M134 | Female |  |
| *Colobus* | *polykomos* |  | Colobinae | Colobini | RMCA 8107M144 | Male |  |
| *Colobus* | *polykomos* |  | Colobinae | Colobini | RMCA 8107M155 | Male |  |
| *Colobus* | *polykomos* |  | Colobinae | Colobini | RMCA 8107M156 | Female |  |
| *Colobus* | *polykomos* |  | Colobinae | Colobini | RMCA 8107M159 | Male |  |
| *Colobus* | *polykomos* |  | Colobinae | Colobini | RMCA 8107M164 | Male |  |
| *Colobus* | *polykomos* |  | Colobinae | Colobini | RMCA 8107M166 | Male |  |
| *Colobus* | *polykomos* |  | Colobinae | Colobini | RMCA 8107M171 | Female |  |
| *Colobus* | *polykomos* |  | Colobinae | Colobini | RMCA 8107M172 | Male |  |
| *Colobus* | *polykomos* |  | Colobinae | Colobini | RMCA 28850 | Male |  |
| *Colobus* | *polykomos* |  | Colobinae | Colobini | RMCA 35124 | Male |  |
| *Colobus* | *polykomos* |  | Colobinae | Colobini | RMCA 84037M38 | Male |  |
| *Colobus* | *polykomos* |  | Colobinae | Colobini | USNM 477321 | Female |  |
| *Colobus* | *polykomos* |  | Colobinae | Colobini | USNM 481784 | Male |  |
| *Colobus* | *polykomos* |  | Colobinae | Colobini | USNM 481785 | Female |  |
| *Colobus* | *polykomos* |  | Colobinae | Colobini | USNM 481786 | Female |  |
| *Erythrocebus* | *patas* |  | Cercopithecinae | Cercopithecini | KUPRI 551 | Female | This study |
| *Erythrocebus* | *patas* |  | Cercopithecinae | Cercopithecini | KUPRI 4237 | Female |  |
| *Erythrocebus* | *patas* |  | Cercopithecinae | Cercopithecini | KUPRI 5399 | Male |  |
| *Erythrocebus* | *patas* |  | Cercopithecinae | Cercopithecini | KUPRI 6447 | Female |  |
| *Erythrocebus* | *patas* |  | Cercopithecinae | Cercopithecini | KUPRI 6448 | Male |  |
| *Erythrocebus* | *patas* |  | Cercopithecinae | Cercopithecini | KUPRI 7145 | Male |  |
| *Erythrocebus* | *patas* |  | Cercopithecinae | Cercopithecini | KUPRI 7170 | Female |  |
| *Erythrocebus* | *patas* |  | Cercopithecinae | Cercopithecini | KUPRI 12078 | Male |  |
| *Erythrocebus* | *patas* |  | Cercopithecinae | Cercopithecini | USNM 410882 | Male |  |
| *Erythrocebus* | *patas* |  | Cercopithecinae | Cercopithecini | USNM 270440 | Female |  |
| *Lophocebus* | *johnstoni* |  | Cercopithecinae | Papionina | AMNH 52598 | Male | https://www.morphosource.org |
| *Lophocebus* | *johnstoni* |  | Cercopithecinae | Papionina | AMNH 52606 | Female |  |
| *Lophocebus* | *johnstoni* |  | Cercopithecinae | Papionina | AMNH 52607 | Female |  |
| *Lophocebus* | *johnstoni* |  | Cercopithecinae | Papionina | AMNH 52613 | Female |  |
| *Lophocebus* | *johnstoni* |  | Cercopithecinae | Papionina | AMNH 52625 | Male |  |
| *Lophocebus* | *atterimus* |  | Cercopithecinae | Papionina | AMNH 55013 | Male |  |
| *Lophocebus* | *atterimus* |  | Cercopithecinae | Papionina | AMNH 82428 | Male |  |
| *Lophocebus* | *atterimus* |  | Cercopithecinae | Papionina | AMNH 86705 | Male |  |
| *Lophocebus* | *albigena* |  | Cercopithecinae | Papionina | USNM 220375 | Female |  |
| *Lophocebus* | *johnstoni* |  | Cercopithecinae | Papionina | USNM 452500 | Male |  |
| *Lophocebus* | *atterimus* |  | Cercopithecinae | Papionina | USNM 503882 | Female |  |
| *Lophocebus* | *albigena* |  | Cercopithecinae | Papionina | MCZ 6209 | Female |  |
| *Lophocebus* | *albigena* |  | Cercopithecinae | Papionina | MCZ 14725 | Female |  |
| *Lophocebus* | *albigena* |  | Cercopithecinae | Papionina | MCZ 18614 | Male |  |
| *Lophocebus* | *albigena* |  | Cercopithecinae | Papionina | MCZ 22737 | Female |  |
| *Lophocebus* | *albigena* |  | Cercopithecinae | Papionina | MCZ 23194 | Male |  |
| *Lophocebus* | *johnstoni* |  | Cercopithecinae | Papionina | MCZ 39396 | Male |  |
| *Lophocebus* | *johnstoni* |  | Cercopithecinae | Papionina | MCZ 39402 | Male |  |
| *Lophocebus* | *albigena* |  | Cercopithecinae | Papionina | KUPRI 1725 | Female | This study |
| *Lophocebus* | *albigena* |  | Cercopithecinae | Papionina | KUPRI 11593 | Male |  |
| *Lophocebus* | *albigena* |  | Cercopithecinae | Papionina | KUPRI 11595 | Female |  |
| *Lophocebus* | *albigena* |  | Cercopithecinae | Papionina | KUPRI 11596 | Female |  |
| *Lophocebus* | *albigena* |  | Cercopithecinae | Papionina | KUPRI 11597 | Female |  |
| *Macaca* | *fascicularis* |  | Cercopithecinae | Macacina | FMNH 1172 | Male | https://www.morphosource.org |
| *Macaca* | *fascicularis* |  | Cercopithecinae | Macacina | FMNH 14805 | ? |  |
| *Macaca* | *fascicularis* |  | Cercopithecinae | Macacina | FMNH 33508 | Female |  |
| *Macaca* | *fascicularis* |  | Cercopithecinae | Macacina | FMNH 46523 | Male |  |
| *Macaca* | *fascicularis* |  | Cercopithecinae | Macacina | FMNH 56160 | Male |  |
| *Macaca* | *fascicularis* |  | Cercopithecinae | Macacina | FMNH 56161 | Male |  |
| *Macaca* | *fascicularis* |  | Cercopithecinae | Macacina | FMNH 56435 | Male |  |
| *Macaca* | *fascicularis* |  | Cercopithecinae | Macacina | FMNH 56490 | Male |  |
| *Macaca* | *fascicularis* |  | Cercopithecinae | Macacina | FMNH 56492 | Female |  |
| *Macaca* | *fascicularis* |  | Cercopithecinae | Macacina | FMNH 56493 | Male |  |
| *Macaca* | *fascicularis* |  | Cercopithecinae | Macacina | FMNH 61026 | Male |  |
| *Macaca* | *fascicularis* |  | Cercopithecinae | Macacina | FMNH 62274 | Female |  |
| *Macaca* | *fascicularis* |  | Cercopithecinae | Macacina | FMNH 62276 | Female |  |
| *Macaca* | *fascicularis* |  | Cercopithecinae | Macacina | FMNH 62901 | Male |  |
| *Macaca* | *fascicularis* |  | Cercopithecinae | Macacina | FMNH 65440 | Male |  |
| *Macaca* | *fascicularis* |  | Cercopithecinae | Macacina | FMNH 65441 | Female |  |
| *Macaca* | *fascicularis* |  | Cercopithecinae | Macacina | FMNH 65442 | Female |  |
| *Macaca* | *fascicularis* |  | Cercopithecinae | Macacina | FMNH 65448 | Male |  |
| *Macaca* | *fascicularis* |  | Cercopithecinae | Macacina | FMNH 65449 | Female |  |
| *Macaca* | *fascicularis* |  | Cercopithecinae | Macacina | FMNH 65451 | Female |  |
| *Macaca* | *fascicularis* |  | Cercopithecinae | Macacina | FMNH 65452 | Male |  |
| *Macaca* | *fascicularis* |  | Cercopithecinae | Macacina | FMNH 66327 | Male |  |
| *Macaca* | *fascicularis* |  | Cercopithecinae | Macacina | FMNH 66328 | Female |  |
| *Macaca* | *fascicularis* |  | Cercopithecinae | Macacina | FMNH 66329 | Female |  |
| *Macaca* | *fascicularis* |  | Cercopithecinae | Macacina | FMNH 66331 | Male |  |
| *Macaca* | *fascicularis* |  | Cercopithecinae | Macacina | FMNH 66332 | Female |  |
| *Macaca* | *fascicularis* |  | Cercopithecinae | Macacina | FMNH 66341 | Male |  |
| *Macaca* | *fascicularis* |  | Cercopithecinae | Macacina | FMNH 66347 | Male |  |
| *Macaca* | *fascicularis* |  | Cercopithecinae | Macacina | FMNH 67719 | Male |  |
| *Macaca* | *fascicularis* |  | Cercopithecinae | Macacina | FMNH 67733 | Male |  |
| *Macaca* | *fascicularis* |  | Cercopithecinae | Macacina | FMNH 67736 | Female |  |
| *Macaca* | *fascicularis* |  | Cercopithecinae | Macacina | FMNH 67990 | Female |  |
| *Macaca* | *fascicularis* |  | Cercopithecinae | Macacina | FMNH 67994 | Male |  |
| *Macaca* | *fascicularis* |  | Cercopithecinae | Macacina | FMNH 67995 | Female |  |
| *Macaca* | *fascicularis* |  | Cercopithecinae | Macacina | FMNH 68701 | Male |  |
| *Macaca* | *fascicularis* |  | Cercopithecinae | Macacina | FMNH 68702 | Female |  |
| *Macaca* | *fascicularis* |  | Cercopithecinae | Macacina | FMNH 75598 | Female |  |
| *Macaca* | *fascicularis* | *philippinensis* | Cercopithecinae | Macacina | FMNH 87424 | ? |  |
| *Macaca* | *fascicularis* | *fascicularis* | Cercopithecinae | Macacina | FMNH 87434 | ? |  |
| *Macaca* | *fascicularis* | *philippinensis* | Cercopithecinae | Macacina | FMNH 87717 | Female |  |
| *Macaca* | *fascicularis* | *philippinensis* | Cercopithecinae | Macacina | FMNH 87718 | Male |  |
| *Macaca* | *fascicularis* | *fascicularis* | Cercopithecinae | Macacina | FMNH 99651 | Male |  |
| *Macaca* | *fascicularis* |  | Cercopithecinae | Macacina | FMNH 99658 | Female |  |
| *Macaca* | *fascicularis* |  | Cercopithecinae | Macacina | FMNH 99659 | Female |  |
| *Macaca* | *fascicularis* |  | Cercopithecinae | Macacina | FMNH 99660 | Female |  |
| *Macaca* | *fascicularis* |  | Cercopithecinae | Macacina | FMNH 105653 | Male |  |
| *Macaca* | *fuscata* | *fuscata* | Cercopithecinae | Macacina | KUPRI 1518 | Female | This study |
| *Macaca* | *fuscata* | *fuscata* | Cercopithecinae | Macacina | KUPRI 1531 | Male |  |
| *Macaca* | *fuscata* | *fuscata* | Cercopithecinae | Macacina | KUPRI 1533 | Female |  |
| *Macaca* | *fuscata* | *fuscata* | Cercopithecinae | Macacina | KUPRI 1543 | Male |  |
| *Macaca* | *fuscata* | *fuscata* | Cercopithecinae | Macacina | KUPRI 1569 | Male |  |
| *Macaca* | *fuscata* | *fuscata* | Cercopithecinae | Macacina | KUPRI 1742 | Female |  |
| *Macaca* | *fuscata* | *fuscata* | Cercopithecinae | Macacina | KUPRI 1743 | Male |  |
| *Macaca* | *fuscata* | *fuscata* | Cercopithecinae | Macacina | KUPRI 1754 | Male |  |
| *Macaca* | *fuscata* | *fuscata* | Cercopithecinae | Macacina | KUPRI 1790 | Male |  |
| *Macaca* | *fuscata* | *fuscata* | Cercopithecinae | Macacina | KUPRI 1791 | Male |  |
| *Macaca* | *fuscata* | *fuscata* | Cercopithecinae | Macacina | KUPRI 1825 | Female |  |
| *Macaca* | *fuscata* | *fuscata* | Cercopithecinae | Macacina | KUPRI 1827 | Female |  |
| *Macaca* | *fuscata* | *fuscata* | Cercopithecinae | Macacina | KUPRI 2069 | Female |  |
| *Macaca* | *fuscata* | *yakui* | Cercopithecinae | Macacina | KUPRI 5227 | Female |  |
| *Macaca* | *fuscata* | *yakui* | Cercopithecinae | Macacina | KUPRI 5416 | Male |  |
| *Macaca* | *fuscata* | *yakui* | Cercopithecinae | Macacina | KUPRI 6084 | Male |  |
| *Macaca* | *fuscata* | *yakui* | Cercopithecinae | Macacina | KUPRI 6627 | Female |  |
| *Macaca* | *fuscata* | *yakui* | Cercopithecinae | Macacina | KUPRI 2265 | Female |  |
| *Macaca* | *fuscata* | *yakui* | Cercopithecinae | Macacina | KUPRI 2473 | Female |  |
| *Macaca* | *fuscata* | *yakui* | Cercopithecinae | Macacina | KUPRI 2595 | Female |  |
| *Macaca* | *fuscata* | *yakui* | Cercopithecinae | Macacina | KUPRI 2597 | Female |  |
| *Macaca* | *fuscata* | *yakui* | Cercopithecinae | Macacina | KUPRI 3126 | Male |  |
| *Macaca* | *fuscata* | *yakui* | Cercopithecinae | Macacina | KUPRI 3127 | Male |  |
| *Macaca* | *fuscata* | *yakui* | Cercopithecinae | Macacina | KUPRI 3454 | Male |  |
| *Macaca* | *fuscata* | *yakui* | Cercopithecinae | Macacina | KUPRI 3958 | Female |  |
| *Macaca* | *fuscata* | *yakui* | Cercopithecinae | Macacina | KUPR4461 | Female |  |
| *Macaca* | *fuscata* | *yakui* | Cercopithecinae | Macacina | KUPRI 4941 | Male |  |
| *Macaca* | *fuscata* | *yakui* | Cercopithecinae | Macacina | KUPRI 7999 | Male |  |
| *Macaca* | *arctoides* | *melanotis* | Cercopithecinae | Macacina | AMNH 54665 | ? | https://www.morphosource.org |
| *Macaca* | *arctoides* |  | Cercopithecinae | Macacina | USNM 111966 | Female |  |
| *Macaca* | *arctoides* |  | Cercopithecinae | Macacina | AMNH 112727 | Female |  |
| *Macaca* | *arctoides* |  | Cercopithecinae | Macacina | AMNH 112730 | Male |  |
| *Macaca* | *arctoides* |  | Cercopithecinae | Macacina | USNM 256825 | Male |  |
| *Macaca* | *assamensis* |  | Cercopithecinae | Macacina | AMNH 43083 | Female |  |
| *Macaca* | *assamensis* |  | Cercopithecinae | Macacina | AMNH 112736 | Male |  |
| *Macaca* | *assamensis* |  | Cercopithecinae | Macacina | AMNH 112738 | Male |  |
| *Macaca* | *assamensis* |  | Cercopithecinae | Macacina | USNM 15255 | Female |  |
| *Macaca* | *assamensis* |  | Cercopithecinae | Macacina | USNM 259725 | Female |  |
| *Macaca* | *brunnescens* | | Cercopithecinae | Macacina | AMNH 30613 | Male |  |
| *Macaca* | *brunnescens* | | Cercopithecinae | Macacina | AMNH 30615 | Female |  |
| *Macaca* | *cyclopis* |  | Cercopithecinae | Macacina | AMNH 184956 | ? |  |
| *Macaca* | *cyclopis* |  | Cercopithecinae | Macacina | AMNH 184957 | ? |  |
| *Macaca* | *cyclopis* |  | Cercopithecinae | Macacina | USNM 296795 | Male |  |
| *Macaca* | *cyclopis* |  | Cercopithecinae | Macacina | USNM 358616 | Female |  |
| *Macaca* | *nemestrina* |  | Cercopithecinae | Macacina | AMNH 11090 | Male |  |
| *Macaca* | *leonina* |  | Cercopithecinae | Macacina | USNM 241022 | Male |  |
| *Macaca* | *leonina* |  | Cercopithecinae | Macacina | USNM 124022 | Female |  |
| *Macaca* | *maura* |  | Cercopithecinae | Macacina | AMNH 153592 | Female |  |
| *Macaca* | *maura* |  | Cercopithecinae | Macacina | AMNH 153594 | Male |  |
| *Macaca* | *maura* |  | Cercopithecinae | Macacina | AMNH 153595 | Male |  |
| *Macaca* | *mulatta* |  | Cercopithecinae | Macacina | AMNH 87264 | Male |  |
| *Macaca* | *mulatta* |  | Cercopithecinae | Macacina | AMNH 112740 | Female |  |
| *Macaca* | *nemestrina* |  | Cercopithecinae | Macacina | AMNH 11091 | ? |  |
| *Macaca* | *nemestrina* | *pagensis* | Cercopithecinae | Macacina | AMNH 103399 | Female |  |
| *Macaca* | *nemestrina* | *nemestrina* | Cercopithecinae | Macacina | AMNH 106037 | Male |  |
| *Macaca* | *nemestrina* | *nemestrina* | Cercopithecinae | Macacina | AMNH 106050 | Female |  |
| *Macaca* | *nemestrina* | *nemestrina* | Cercopithecinae | Macacina | USNM 114502 | Female |  |
| *Macaca* | *nemestrina* | *nemestrina* | Cercopithecinae | Macacina | USNM 123144 | Male |  |
| *Macaca* | *nemestrina* | *nemestrina* | Cercopithecinae | Macacina | USNM 154367 | Male |  |
| *Macaca* | *nemestrina* | *nemestrina* | Cercopithecinae | Macacina | USNM 305069 | Female |  |
| *Macaca* | *nemestrina* | *nemestrina* | Cercopithecinae | Macacina | USNM 399506 | ? |  |
| *Macaca* | *nigra* |  | Cercopithecinae | Macacina | AMNH 196410 | Female |  |
| *Macaca* | *radiata* |  | Cercopithecinae | Macacina | AMNH 163078 | Male |  |
| *Macaca* | *radiata* |  | Cercopithecinae | Macacina | USNM 398463 | Female |  |
| *Macaca* | *siberu* |  | Cercopithecinae | Macacina | USNM 546835 | Male |  |
| *Macaca* | *silenus* |  | Cercopithecinae | Macacina | USNM 574135 | Male |  |
| *Macaca* | *sinica* |  | Cercopithecinae | Macacina | USNM 15259 | Male |  |
| *Macaca* | *sinica* |  | Cercopithecinae | Macacina | USNM 271190 | Female |  |
| *Macaca* | *sylvanus* |  | Cercopithecinae | Macacina | AMNH 19014 | Female |  |
| *Macaca* | *sylvanus* |  | Cercopithecinae | Macacina | USNM 476791 | Male |  |
| *Macaca* | *sylvanus* |  | Cercopithecinae | Macacina | BSZM 1961-168 | Female | This study |
| *Macaca* | *sylvanus* |  | Cercopithecinae | Macacina | BSZM 1981-488 | Male |  |
| *Macaca* | *sylvanus* |  | Cercopithecinae | Macacina | BSZM AM1423 | Female |  |
| *Macaca* | *sylvanus* |  | Cercopithecinae | Macacina | USNM 255979 | Male | https://www.morphosource.org |
| *Macaca* | *sylvanus* |  | Cercopithecinae | Macacina | USNM 476782 | Female |  |
| *Macaca* | *thibetana* |  | Cercopithecinae | Macacina | AMNH 129 | Male |  |
| *Macaca* | *thibetana* |  | Cercopithecinae | Macacina | AMNH 83994 | Female |  |
| *Macaca* | *thibetana* |  | Cercopithecinae | Macacina | AMNH 84472 | Male |  |
| *Macaca* | *thibetana* |  | Cercopithecinae | Macacina | USNM 241162 | Female |  |
| *Macaca* | *thibetana* |  | Cercopithecinae | Macacina | USNM 241163 | Male |  |
| *Macaca* | *tonkeana* | *?* | Cercopithecinae | Macacina | AMNH 153388 | Female |  |
| *Macaca* | *tonkeana* | *hecki* | Cercopithecinae | Macacina | AMNH 152907 | Male |  |
| *Macaca* | *tonkeana* | *?* | Cercopithecinae | Macacina | AMNH 153382 | Male |  |
| *Macaca* | *tonkeana* | *?* | Cercopithecinae | Macacina | AMNH 153401 | Male |  |
| *Macaca* | *tonkeana* | *hecki* | Cercopithecinae | Macacina | AMNH 196405 | Male |  |
| *Macaca* | *tonkeana* | *hecki* | Cercopithecinae | Macacina | AMNH 196407 | Female |  |
| *Macaca* | *fascicularis* | *philippinensis* | Cercopithecinae | Macacina | USNM 114140 | Female |  |
| *Macaca* | *fascicularis* | *fuscus* | Cercopithecinae | Macacina | USNM 114162 | Female |  |
| *Macaca* | *fascicularis* | *fuscus* | Cercopithecinae | Macacina | USNM 114165 | Female |  |
| *Macaca* | *fascicularis* | *fuscus* | Cercopithecinae | Macacina | USNM 114167 | Male |  |
| *Macaca* | *fascicularis* | *fuscus* | Cercopithecinae | Macacina | USNM 114168 | Male |  |
| *Macaca* | *fascicularis* | *fuscus* | Cercopithecinae | Macacina | USNM 114169 | Male |  |
| *Macaca* | *fascicularis* | *fascicularis* | Cercopithecinae | Macacina | USNM 114410 | Male |  |
| *Macaca* | *fascicularis* | *fuscus* | Cercopithecinae | Macacina | USNM 121511 | Male |  |
| *Macaca* | *fascicularis* | *fuscus* | Cercopithecinae | Macacina | USNM 121512 | Male |  |
| *Macaca* | *fascicularis* | *fuscus* | Cercopithecinae | Macacina | USNM 121513 | Female |  |
| *Macaca* | *fascicularis* | *fascicularis* | Cercopithecinae | Macacina | USNM 121869 | Female |  |
| *Macaca* | *fascicularis* | *fascicularis* | Cercopithecinae | Macacina | USNM 121871 | Male |  |
| *Macaca* | *fascicularis* | *fascicularis* | Cercopithecinae | Macacina | USNM 121872 | Male |  |
| *Macaca* | *fascicularis* | *fascicularis* | Cercopithecinae | Macacina | USNM 122848 | Female |  |
| *Macaca* | *fascicularis* | *fascicularis* | Cercopithecinae | Macacina | USNM 123992 | Female |  |
| *Macaca* | *fascicularis* | *fascicularis* | Cercopithecinae | Macacina | USNM 125102 | Male |  |
| *Macaca* | *fascicularis* | *fascicularis* | Cercopithecinae | Macacina | USNM 143582 | Male |  |
| *Macaca* | *fascicularis* | *philippinensis* | Cercopithecinae | Macacina | USNM 144678 | Male |  |
| *Macaca* | *fascicularis* | *philippinensis* | Cercopithecinae | Macacina | USNM 144681 | Female |  |
| *Macaca* | *fascicularis* | *philippinensis* | Cercopithecinae | Macacina | USNM 144683 | Female |  |
| *Macaca* | *fascicularis* | *philippinensis* | Cercopithecinae | Macacina | USNM 144684 | Female |  |
| *Macaca* | *fascicularis* | *philippinensis* | Cercopithecinae | Macacina | USNM 144685 | Female |  |
| *Macaca* | *fascicularis* | *fascicularis* | Cercopithecinae | Macacina | USNM 151830 | Female |  |
| *Macaca* | *fascicularis* | *fascicularis* | Cercopithecinae | Macacina | USNM 156291 | Male |  |
| *Macaca* | *fascicularis* | *fascicularis* | Cercopithecinae | Macacina | USNM 156456 | Female |  |
| *Macaca* | *fascicularis* | *fascicularis* | Cercopithecinae | Macacina | USNM 156458 | Male |  |
| *Macaca* | *fascicularis* | *fascicularis* | Cercopithecinae | Macacina | USNM 175896 | Female |  |
| *Macaca* | *fascicularis* | *tua* | Cercopithecinae | Macacina | USNM 197662 | Male |  |
| *Macaca* | *fascicularis* | *?* | Cercopithecinae | Macacina | USNM 251660 | ? |  |
| *Macaca* | *fascicularis* | *?* | Cercopithecinae | Macacina | USNM 344989 | Female |  |
| *Macaca* | *fascicularis* | *philippinensis* | Cercopithecinae | Macacina | USNM 477842 | Male |  |
| *Macaca* | *fascicularis* | *philippinensis* | Cercopithecinae | Macacina | USNM 477843 | ? |  |
| *Macaca* | *fascicularis* | *philippinensis* | Cercopithecinae | Macacina | USNM 477845 | ? |  |
| *Macaca* | *fascicularis* | *philippinensis* | Cercopithecinae | Macacina | USNM 477846 | Male |  |
| *Macaca* | *fascicularis* | *philippinensis* | Cercopithecinae | Macacina | USNM 477847 | Female |  |
| *Macaca* | *fascicularis* | *?* | Cercopithecinae | Macacina | USNM 521839 | Male |  |
| *Macaca* | *fascicularis* | *?* | Cercopithecinae | Macacina | USNM 536025 | Male |  |
| *Mandrillus* | *sphinx* |  | Cercopithecinae | Papionina | AMNH A99-1-2047 | Male |  |
| *Mandrillus* | *sphinx* |  | Cercopithecinae | Papionina | AMNH A99-1-2048 | Male |  |
| *Mandrillus* | *sphinx* |  | Cercopithecinae | Papionina | AMNH A99-1-2049 | Male |  |
| *Mandrillus* | *sphinx* |  | Cercopithecinae | Papionina | AMNH A99-1-2056 | Male |  |
| *Mandrillus* | *sphinx* |  | Cercopithecinae | Papionina | USNM 283109 | ? |  |
| *Mandrillus* | *leucophaeus* |  | Cercopithecinae | Papionina | USNM 395698 | Male |  |
| *Mandrillus* | *sphinx* |  | Cercopithecinae | Papionina | USNM 598494 | Male |  |
| *Miopithecus* | *ogouensis* |  | Cercopithecinae | Cercopithecini | USNM 598519 | Male |  |
| *Miopithecus* | *ogouensis* |  | Cercopithecinae | Cercopithecini | USNM 598534 | Female |  |
| *Miopithecus* | *ogouensis* |  | Cercopithecinae | Cercopithecini | MCZ 19976 | Male |  |
| *Miopithecus* | *ogouensis* |  | Cercopithecinae | Cercopithecini | MCZ 23196 | Male |  |
| *Miopithecus* | *ogouensis* |  | Cercopithecinae | Cercopithecini | MCZ 23197 | Male |  |
| *Miopithecus* | *ogouensis* |  | Cercopithecinae | Cercopithecini | MCZ 34264 | Female |  |
| *Miopithecus* | *ogouensis* |  | Cercopithecinae | Cercopithecini | MCZ 37278 | Male |  |
| *Nasalis* | *larvatus* |  | Colobinae | Presbytini | AMNH 28255 | Female |  |
| *Nasalis* | *larvatus* |  | Colobinae | Presbytini | AMNH 85171 | Male |  |
| *Nasalis* | *larvatus* |  | Colobinae | Presbytini | AMNH 85172 | Female |  |
| *Nasalis* | *larvatus* |  | Colobinae | Presbytini | AMNH 103365 | Female |  |
| *Nasalis* | *larvatus* |  | Colobinae | Presbytini | AMNH 103402 | Female |  |
| *Nasalis* | *larvatus* |  | Colobinae | Presbytini | AMNH 103458 | Female |  |
| *Nasalis* | *larvatus* |  | Colobinae | Presbytini | AMNH 103464 | Female |  |
| *Nasalis* | *larvatus* |  | Colobinae | Presbytini | AMNH 103465 | Female |  |
| *Nasalis* | *larvatus* |  | Colobinae | Presbytini | AMNH 103466 | Female |  |
| *Nasalis* | *larvatus* |  | Colobinae | Presbytini | AMNH 103669 | Female |  |
| *Nasalis* | *larvatus* |  | Colobinae | Presbytini | AMNH 106029 | Male |  |
| *Nasalis* | *larvatus* |  | Colobinae | Presbytini | AMNH 106030 | Female |  |
| *Nasalis* | *larvatus* |  | Colobinae | Presbytini | AMNH 106032 | Male |  |
| *Nasalis* | *larvatus* |  | Colobinae | Presbytini | AMNH 106033 | Female |  |
| *Nasalis* | *larvatus* |  | Colobinae | Presbytini | AMNH 106034 | Female |  |
| *Nasalis* | *larvatus* |  | Colobinae | Presbytini | AMNH 106272 | Male |  |
| *Nasalis* | *larvatus* |  | Colobinae | Presbytini | AMNH 107101 | Male |  |
| *Nasalis* | *larvatus* |  | Colobinae | Presbytini | MCZ 37339 | Female |  |
| *Nasalis* | *larvatus* |  | Colobinae | Presbytini | MCZ 41560 | Female |  |
| *Nasalis* | *larvatus* |  | Colobinae | Presbytini | MCZ 41562 | Female |  |
| *Nasalis* | *larvatus* |  | Colobinae | Presbytini | USNM 142217 | Male |  |
| *Nasalis* | *larvatus* |  | Colobinae | Presbytini | USNM 196799 | Male |  |
| *Nasalis* | *larvatus* |  | Colobinae | Presbytini | USNM 198277 | Female |  |
| *Nasalis* | *larvatus* |  | Colobinae | Presbytini | FMNH 68682 | Male |  |
| *Nasalis* | *larvatus* |  | Colobinae | Presbytini | FMNH 68683 | Male |  |
| *Nasalis* | *larvatus* |  | Colobinae | Presbytini | FMNH 68684 | Male |  |
| *Nasalis* | *larvatus* |  | Colobinae | Presbytini | FMNH 33546 | Female |  |
| *Nasalis* | *larvatus* |  | Colobinae | Presbytini | FMNH 43106 | Male |  |
| *Nasalis* | *larvatus* |  | Colobinae | Presbytini | FMNH 43107 | Female |  |
| *Nasalis* | *larvatus* |  | Colobinae | Presbytini | AMNH 103671 | Male |  |
| *Nasalis* | *larvatus* |  | Colobinae | Presbytini | USNM 142218 | Male |  |
| *Nasalis* | *larvatus* |  | Colobinae | Presbytini | USNM 142224 | Female |  |
| *Nasalis* | *larvatus* |  | Colobinae | Presbytini | USNM 145325 | Female |  |
| *Nasalis* | *larvatus* |  | Colobinae | Presbytini | USNM 151817 | Female |  |
| *Nasalis* | *larvatus* |  | Colobinae | Presbytini | USNM 196786 | Male |  |
| *Nasalis* | *larvatus* |  | Colobinae | Presbytini | USNM 196787 | Male |  |
| *Nasalis* | *larvatus* |  | Colobinae | Presbytini | USNM 196788 | Male |  |
| *Nasalis* | *larvatus* |  | Colobinae | Presbytini | USNM 196789 | Male |  |
| *Nasalis* | *larvatus* |  | Colobinae | Presbytini | USNM 196793 | Female |  |
| *Nasalis* | *larvatus* |  | Colobinae | Presbytini | USNM 196794 | Female |  |
| *Nasalis* | *larvatus* |  | Colobinae | Presbytini | USNM 196795 | Female |  |
| *Nasalis* | *larvatus* |  | Colobinae | Presbytini | USNM 196796 | Female |  |
| *Nasalis* | *larvatus* |  | Colobinae | Presbytini | USNM 197644 | Male |  |
| *Nasalis* | *larvatus* |  | Colobinae | Presbytini | USNM 198276 | Male |  |
| *Nasalis* | *larvatus* |  | Colobinae | Presbytini | USNM 198845 | Female |  |
| *Nasalis* | *larvatus* |  | Colobinae | Presbytini | USNM 198846 | Male |  |
| *Nasalis* | *larvatus* |  | Colobinae | Presbytini | USNM 198847 | Female |  |
| *Nasalis* | *larvatus* |  | Colobinae | Presbytini | USNM 521841 | Male |  |
| *Papio* | *anubis* |  | Cercopithecinae | Papionina | RMCA 960 | Male |  |
| *Papio* | *anubis* |  | Cercopithecinae | Papionina | RMCA 4200(2400) | Male |  |
| *Papio* | *anubis* |  | Cercopithecinae | Papionina | RMCA 6026 | Male |  |
| *Papio* | *anubis* |  | Cercopithecinae | Papionina | RMCA 8311 | Male |  |
| *Papio* | *anubis* |  | Cercopithecinae | Papionina | RMCA 8312 | Female |  |
| *Papio* | *anubis* |  | Cercopithecinae | Papionina | RMCA 8461 | Female |  |
| *Papio* | *anubis* |  | Cercopithecinae | Papionina | RMCA 8462 | Female |  |
| *Papio* | *anubis* |  | Cercopithecinae | Papionina | RMCA 8464 | Male |  |
| *Papio* | *anubis* |  | Cercopithecinae | Papionina | RMCA 9244 | Male |  |
| *Papio* | *anubis* |  | Cercopithecinae | Papionina | RMCA 10222 | Female |  |
| *Papio* | *anubis* |  | Cercopithecinae | Papionina | RMCA 12441 | Female |  |
| *Papio* | *anubis* |  | Cercopithecinae | Papionina | RMCA 14691 | Female |  |
| *Papio* | *anubis* |  | Cercopithecinae | Papionina | RMCA 17691 | Female |  |
| *Papio* | *anubis* |  | Cercopithecinae | Papionina | RMCA 17738 | Male |  |
| *Papio* | *anubis* |  | Cercopithecinae | Papionina | RMCA 18470 | Male |  |
| *Papio* | *anubis* |  | Cercopithecinae | Papionina | RMCA 18890 | Female |  |
| *Papio* | *anubis* |  | Cercopithecinae | Papionina | RMCA 21120 | Female |  |
| *Papio* | *anubis* |  | Cercopithecinae | Papionina | RMCA 21593 | Male |  |
| *Papio* | *anubis* |  | Cercopithecinae | Papionina | RMCA 25474 | Male |  |
| *Papio* | *anubis* |  | Cercopithecinae | Papionina | RMCA 83006M9 | Female |  |
| *Papio* | *ursinus* |  | Cercopithecinae | Papionina | RMCA 85049M3 | Male |  |
| *Papio* | *anubis* |  | Cercopithecinae | Papionina | RMCA 85049M4 | Male |  |
| *Papio* | *anubis* |  | Cercopithecinae | Papionina | RMCA 85049M6 | Male |  |
| *Papio* | *anubis* |  | Cercopithecinae | Papionina | RMCA 18472 | Female |  |
| *Papio* | *anubis* |  | Cercopithecinae | Papionina | USNM 397476 | Female |  |
| *Papio* | *anubis* |  | Cercopithecinae | Papionina | USNM 162899 | Male |  |
| *Papio* | *anubis* |  | Cercopithecinae | Papionina | AMNH 52674 | Female |  |
| *Papio* | *papio* |  | Cercopithecinae | Papionina | RMCA 37565 | Male |  |
| *Papio* | *papio* |  | Cercopithecinae | Papionina | RMCA 37567 | Male |  |
| *Papio* | *papio* |  | Cercopithecinae | Papionina | RMCA 37569 | Male |  |
| *Papio* | *papio* |  | Cercopithecinae | Papionina | USNM 378669 | Male |  |
| *Papio* | *papio* |  | Cercopithecinae | Papionina | USNM 381430 | Female |  |
| *Papio* | *cynocephalus* | | Cercopithecinae | Papionina | AMNH 54227 | Male |  |
| *Papio* | *cynocephalus* | | Cercopithecinae | Papionina | AMNH 161734 | Male |  |
| *Papio* | *hamadryas* |  | Cercopithecinae | Papionina | USNM 258502 | Male |  |
| *Papio* | *anubis* |  | Cercopithecinae | Papionina | AMNH 51380 | Male |  |
| *Papio* | *ursinus* |  | Cercopithecinae | Papionina | AMNH 81811 | ? |  |
| *Papio* | *ursinus* |  | Cercopithecinae | Papionina | AMNH 216250 | ? |  |
| *Papio* | *ursinus* |  | Cercopithecinae | Papionina | AMNH 216251 | ? |  |
| *Papio* | *anubis* |  | Cercopithecinae | Papionina | RMCA 2230 | Male |  |
| *Papio* | *anubis* |  | Cercopithecinae | Papionina | RMCA 7309M49 | Female |  |
| *Papio* | *anubis* |  | Cercopithecinae | Papionina | RMCA 8044M102 | Male |  |
| *Papio* | *anubis* |  | Cercopithecinae | Papionina | RMCA 90042M224 | Male |  |
| *Papio* | *anubis* |  | Cercopithecinae | Papionina | RMCA 90042M225 | Male |  |
| *Papio* | *anubis* |  | Cercopithecinae | Papionina | RMCA 90042M228 | Female |  |
| *Papio* | *anubis* |  | Cercopithecinae | Papionina | RMCA 12575 | Male |  |
| *Papio* | *anubis* |  | Cercopithecinae | Papionina | RMCA 21422 | Female |  |
| *Papio* | *anubis* |  | Cercopithecinae | Papionina | RMCA 25734 | Male |  |
| *Papio* | *anubis* |  | Cercopithecinae | Papionina | RMCA 25735 | Male |  |
| *Papio* | *anubis* |  | Cercopithecinae | Papionina | RMCA 25738 | Male |  |
| *Papio* | *anubis* |  | Cercopithecinae | Papionina | RMCA 25739 | Male |  |
| *Papio* | *anubis* |  | Cercopithecinae | Papionina | RMCA 73015M336 | Female |  |
| *Papio* | *anubis* |  | Cercopithecinae | Papionina | RMCA 73018M187 | Female |  |
| *Papio* | *anubis* |  | Cercopithecinae | Papionina | RMCA 83006M1 | Male |  |
| *Papio* | *anubis* |  | Cercopithecinae | Papionina | RMCA 83006M2 | Male |  |
| *Papio* | *anubis* |  | Cercopithecinae | Papionina | RMCA 83006M3 | Male |  |
| *Papio* | *anubis* |  | Cercopithecinae | Papionina | RMCA 83006M4 | Male |  |
| *Papio* | *anubis* |  | Cercopithecinae | Papionina | RMCA 83006M5 | Male |  |
| *Papio* | *anubis* |  | Cercopithecinae | Papionina | RMCA 83006M6 | Male |  |
| *Papio* | *anubis* |  | Cercopithecinae | Papionina | RMCA 83006M10 | Female |  |
| *Papio* | *anubis* |  | Cercopithecinae | Papionina | RMCA 84029M9 | Male |  |
| *Papio* | *anubis* |  | Cercopithecinae | Papionina | RMCA 84029M10 | Male |  |
| *Piliocolobus* | *badius* | *epieni* | Colobinae | Colobini | AMNH ED5 | Female |  |
| *Piliocolobus* | *badius* | *badius* | Colobinae | Colobini | MCZ 24080 | Male |  |
| *Piliocolobus* | *badius* | *badius* | Colobinae | Colobini | MCZ 24775 | Male |  |
| *Piliocolobus* | *badius* | *badius* | Colobinae | Colobini | MCZ 25627 | Male |  |
| *Piliocolobus* | *badius* | *badius* | Colobinae | Colobini | MCZ 25810 | Female |  |
| *Piliocolobus* | *gordonorum* |  | Colobinae | Colobini | MCZ 26552 | Female |  |
| *Piliocolobus* | *gordonorum* |  | Colobinae | Colobini | MCZ 26553 | Female |  |
| *Piliocolobus* | *preussi* |  | Colobinae | Colobini | MCZ 27108 | Female |  |
| *Piliocolobus* | *foai* |  | Colobinae | Colobini | RMCA 91.060-M-0069 | Female |  |
| *Piliocolobus* | *badius* | *temminckii* | Colobinae | Colobini | USNM 378673 | Male |  |
| *Piliocolobus* | *badius* | *temminckii* | Colobinae | Colobini | USNM 378674 | Female |  |
| *Piliocolobus* | *badius* | *badius* | Colobinae | Colobini | USNM 481792 | Female |  |
| *Piliocolobus* | *badius* | *badius* | Colobinae | Colobini | USNM 481795 | Male |  |
| *Piliocolobus* | *sp* |  | Colobinae | Colobini | KAS 175 | Female | This study |
| *Piliocolobus* | *badius* |  | Colobinae | Colobini | KUPRI 454 | Female | https://www.morphosource.org |
| *Piliocolobus* | *badius* |  | Colobinae | Colobini | KUPRI 456 | Female |  |
| *Piliocolobus* | *badius* |  | Colobinae | Colobini | KUPRI 458 | Female |  |
| *Piliocolobus* | *badius* |  | Colobinae | Colobini | KUPRI 464 | Female |  |
| *Piliocolobus* | *badius* |  | Colobinae | Colobini | KUPRI 500 | Female |  |
| *Piliocolobus* | *badius* |  | Colobinae | Colobini | KUPRI 507 | Female |  |
| *Piliocolobus* | *badius* |  | Colobinae | Colobini | KUPRI 518 | Female |  |
| *Piliocolobus* | *badius* |  | Colobinae | Colobini | KUPRI 520 | Female |  |
| *Piliocolobus* | *badius* |  | Colobinae | Colobini | KUPRI 522 | Female |  |
| *Piliocolobus* | *badius* |  | Colobinae | Colobini | KUPRI 524 | Female |  |
| *Piliocolobus* | *badius* |  | Colobinae | Colobini | KUPRI 396 | Male |  |
| *Piliocolobus* | *badius* |  | Colobinae | Colobini | KUPRI 441 | Male |  |
| *Piliocolobus* | *badius* |  | Colobinae | Colobini | KUPRI 447 | Male |  |
| *Piliocolobus* | *badius* |  | Colobinae | Colobini | KUPRI 462 | Male |  |
| *Piliocolobus* | *badius* |  | Colobinae | Colobini | KUPRI 465 | Male |  |
| *Piliocolobus* | *badius* |  | Colobinae | Colobini | KUPRI 477 | Male |  |
| *Piliocolobus* | *badius* |  | Colobinae | Colobini | KUPRI 490 | Male |  |
| *Piliocolobus* | *badius* |  | Colobinae | Colobini | KUPRI 4786 | Male |  |
| *Piliocolobus* | *badius* |  | Colobinae | Colobini | KUPRI 4787 | Male |  |
| *Piliocolobus* | *badius* |  | Colobinae | Colobini | KUPRI 4791 | Male |  |
| *Piliocolobus* | *kirkii* |  | Colobinae | Colobini | USNM 452646 | Female |  |
| *Piliocolobus* | *tephrosceles* |  | Colobinae | Colobini | USNM 452644 | Male |  |
| *Piliocolobus* | *rufomitratus* | | Colobinae | Colobini | OM 3046 | Male | This study |
| *Piliocolobus* | *sp* |  | Colobinae | Colobini | KAS 178 | ? |  |
| *Piliocolobus* | *badius* |  | Colobinae | Colobini | RMCA 8107M98 | ? | https://www.morphosource.org |
| *Piliocolobus* | *badius* |  | Colobinae | Colobini | RMCA 8107M97 | ? |  |
| *Piliocolobus* | *badius* |  | Colobinae | Colobini | RMCA 8107M118 | ? |  |
| *Presbytis* | *natunae* |  | Colobinae | Presbytini | USNM 104845 | Female |  |
| *Presbytis* | *natunae* |  | Colobinae | Presbytini | USNM 104847 | Male |  |
| *Presbytis* | *potenziani* |  | Colobinae | Presbytini | USNM 121668 | Male |  |
| *Presbytis* | *siberu* |  | Colobinae | Presbytini | USNM 121692 | Female |  |
| *Presbytis* | *siberu* |  | Colobinae | Presbytini | USNM 544968 | Male |  |
| *Presbytis* | *potenziani* |  | Colobinae | Presbytini | USNM 544970 | Female |  |
| *Presbytis* | *potenziani* |  | Colobinae | Presbytini | KUPRI 1145 | Female | This study |
| *Presbytis* | *potenziani* |  | Colobinae | Presbytini | KUPRI 1147 | Female |  |
| *Presbytis* | *potenziani* |  | Colobinae | Presbytini | KUPRI 1151 | Male |  |
| *Presbytis* | *potenziani* |  | Colobinae | Presbytini | KUPRI 1153 | Male |  |
| *Presbytis* | *potenziani* |  | Colobinae | Presbytini | KUPRI 1154 | Male |  |
| *Presbytis* | *potenziani* |  | Colobinae | Presbytini | KUPRI 1156 | Male |  |
| *Presbytis* | *potenziani* |  | Colobinae | Presbytini | KUPRI 1160 | Male |  |
| *Presbytis* | *potenziani* |  | Colobinae | Presbytini | KUPRI 1161 | Female |  |
| *Presbytis* | *potenziani* |  | Colobinae | Presbytini | KUPRI 1163 | ?Female |  |
| *Presbytis* | *potenziani* |  | Colobinae | Presbytini | KUPRI 1163-2 | ?Male |  |
| *Presbytis* | *potenziani* |  | Colobinae | Presbytini | KUPRI 1164 | Male |  |
| *Presbytis* | *potenziani* |  | Colobinae | Presbytini | KUPRI 1167 | Male |  |
| *Presbytis* | *potenziani* |  | Colobinae | Presbytini | KUPRI 1173 | Female |  |
| *Presbytis* | *potenziani* |  | Colobinae | Presbytini | KUPRI 1175 | Male |  |
| *Presbytis* | *potenziani* |  | Colobinae | Presbytini | KUPRI 1180 | Male |  |
| *Presbytis* | *potenziani* |  | Colobinae | Presbytini | KUPRI 1181 | Female |  |
| *Presbytis* | *potenziani* |  | Colobinae | Presbytini | KUPRI 1182 | Female |  |
| *Presbytis* | *potenziani* |  | Colobinae | Presbytini | KUPRI 1183 | Female |  |
| *Presbytis* | *potenziani* |  | Colobinae | Presbytini | KUPRI 1197 | Female |  |
| *Presbytis* | *potenziani* |  | Colobinae | Presbytini | KUPRI 1877 | Female |  |
| *Presbytis* | *canicrus* |  | Colobinae | Presbytini | USNM 198282 | Female | https://www.morphosource.org |
| *Presbytis* | *rubicunda* |  | Colobinae | Presbytini | AMNH 103634 | Female |  |
| *Presbytis* | *rubicunda* |  | Colobinae | Presbytini | AMNH 103637 | Male |  |
| *Presbytis* | *rubicunda* |  | Colobinae | Presbytini | MCZ 35704 | Female |  |
| *Presbytis* | *rubicunda* |  | Colobinae | Presbytini | MCZ 35705 | Female |  |
| *Presbytis* | *rubicunda* |  | Colobinae | Presbytini | MCZ 35712 | Male |  |
| *Presbytis* | *rubicunda* |  | Colobinae | Presbytini | MCZ 37666 | Female |  |
| *Presbytis* | *rubicunda* |  | Colobinae | Presbytini | MCZ 37776 | Male |  |
| *Presbytis* | *rubicunda* |  | Colobinae | Presbytini | USNM 125157 | Female |  |
| *Presbytis* | *rubicunda* |  | Colobinae | Presbytini | USNM 153790 | Female |  |
| *Presbytis* | *siamensis* | *catemana* | Colobinae | Presbytini | KUPRI 4557 | Female | This study |
| *Presbytis* | *siamensis* | *catemana* | Colobinae | Presbytini | KUPRI 4560 | Female |  |
| *Presbytis* | *siamensis* | *catemana* | Colobinae | Presbytini | KUPRI 4561 | Female |  |
| *Presbytis* | *siamensis* | *catemana* | Colobinae | Presbytini | KUPRI 4559 | Female |  |
| *Presbytis* | *siamensis* | *catemana* | Colobinae | Presbytini | KUPRI 4558 | Male |  |
| *Presbytis* | *siamensis* | *catemana* | Colobinae | Presbytini | KUPRI 4565 | Male |  |
| *Presbytis* | *siamensis* | *catemana* | Colobinae | Presbytini | KUPRI 4568 | Male |  |
| *Presbytis* | *siamensis* | *catemana* | Colobinae | Presbytini | KUPRI 4564 | Male |  |
| *Presbytis* | *siamensis* | *paenulata* | Colobinae | Presbytini | KUPRI 10004 | Female |  |
| *Presbytis* | *siamensis* | *paenulata* | Colobinae | Presbytini | KUPRI 4550 | Female |  |
| *Presbytis* | *siamensis* | *paenulata* | Colobinae | Presbytini | KUPRI 4553 | Female |  |
| *Presbytis* | *siamensis* | *paenulata* | Colobinae | Presbytini | KUPRI 4555 | Female |  |
| *Presbytis* | *siamensis* | *paenulata* | Colobinae | Presbytini | KUPRI 4551 | Male |  |
| *Presbytis* | *siamensis* | *paenulata* | Colobinae | Presbytini | KUPRI 4549 | Male |  |
| *Presbytis* | *bicolor* |  | Colobinae | Presbytini | KUPRI 4512 | Female |  |
| *Presbytis* | *bicolor* |  | Colobinae | Presbytini | KUPRI 4519 | Female |  |
| *Presbytis* | *bicolor* |  | Colobinae | Presbytini | KUPRI 4520 | Female |  |
| *Presbytis* | *bicolor* |  | Colobinae | Presbytini | KUPRI 4526 | Female |  |
| *Presbytis* | *bicolor* |  | Colobinae | Presbytini | KUPRI 4527 | Female |  |
| *Presbytis* | *bicolor* |  | Colobinae | Presbytini | KUPRI 4528 | Female |  |
| *Presbytis* | *bicolor* |  | Colobinae | Presbytini | KUPRI 4533 | Female |  |
| *Presbytis* | *bicolor* |  | Colobinae | Presbytini | KUPRI 4536 | Female |  |
| *Presbytis* | *bicolor* |  | Colobinae | Presbytini | KUPRI 4541 | Female |  |
| *Presbytis* | *bicolor* |  | Colobinae | Presbytini | KUPRI 4545 | Female |  |
| *Presbytis* | *bicolor* |  | Colobinae | Presbytini | KUPRI 4518 | Male |  |
| *Presbytis* | *bicolor* |  | Colobinae | Presbytini | KUPRI 4530 | Male |  |
| *Presbytis* | *bicolor* |  | Colobinae | Presbytini | KUPRI 4532 | Male |  |
| *Presbytis* | *bicolor* |  | Colobinae | Presbytini | KUPRI 4535 | Male |  |
| *Presbytis* | *bicolor* |  | Colobinae | Presbytini | KUPRI 4542 | Male |  |
| *Presbytis* | *bicolor* |  | Colobinae | Presbytini | KUPRI 4543 | Male |  |
| *Presbytis* | *bicolor* |  | Colobinae | Presbytini | KUPRI 4544 | Male |  |
| *Presbytis* | *bicolor* |  | Colobinae | Presbytini | KUPRI 4546 | Male |  |
| *Presbytis* | *bicolor* |  | Colobinae | Presbytini | KUPRI 4548 | Male |  |
| *Presbytis* | *bicolor* |  | Colobinae | Presbytini | KUPRI 5841 | Male |  |
| *Procolobus* | *verus* |  | Colobinae | Colobini | AMNH 89438 | Female | https://www.morphosource.org |
| *Procolobus* | *verus* |  | Colobinae | Colobini | AMNH 89439 | Male |  |
| *Procolobus* | *verus* |  | Colobinae | Colobini | RMCA 86.002-M-0066 | Female |  |
| *Procolobus* | *verus* |  | Colobinae | Colobini | RMCA 86.002-M-0069 | Male |  |
| *Procolobus* | *verus* |  | Colobinae | Colobini | RMCA 84.037-M-0176 | Male |  |
| *Procolobus* | *verus* |  | Colobinae | Colobini | USNM 477327 | Female |  |
| *Procolobus* | *verus* |  | Colobinae | Colobini | KUPRI 354 | Female | This study |
| *Procolobus* | *verus* |  | Colobinae | Colobini | KUPRI 378 | Female |  |
| *Procolobus* | *verus* |  | Colobinae | Colobini | KUPRI 381 | Female |  |
| *Procolobus* | *verus* |  | Colobinae | Colobini | KUPRI 383 | Female |  |
| *Procolobus* | *verus* |  | Colobinae | Colobini | KUPRI 388 | Female |  |
| *Procolobus* | *verus* |  | Colobinae | Colobini | KUPRI 390 | Female |  |
| *Procolobus* | *verus* |  | Colobinae | Colobini | KUPRI 391 | Female |  |
| *Procolobus* | *verus* |  | Colobinae | Colobini | KUPRI 392 | Female |  |
| *Procolobus* | *verus* |  | Colobinae | Colobini | KUPRI 372 | Male |  |
| *Procolobus* | *verus* |  | Colobinae | Colobini | KUPRI 373 | Male |  |
| *Procolobus* | *verus* |  | Colobinae | Colobini | KUPRI 374 | Male |  |
| *Procolobus* | *verus* |  | Colobinae | Colobini | KUPRI 375 | Male |  |
| *Procolobus* | *verus* |  | Colobinae | Colobini | KUPRI 379 | Male |  |
| *Procolobus* | *verus* |  | Colobinae | Colobini | KUPRI 377 | Male |  |
| *Procolobus* | *verus* |  | Colobinae | Colobini | KUPRI 393 | Male |  |
| *Procolobus* | *verus* |  | Colobinae | Colobini | KUPRI 394 | Male |  |
| *Procolobus* | *verus* |  | Colobinae | Colobini | KUPRI 395 | Male |  |
| *Pygathrix* | *nigripes* |  | Colobinae | Presbytini | USNM 269798 | Male | https://www.morphosource.org |
| *Pygathrix* | *nemaeus* |  | Colobinae | Presbytini | USNM 356574 | Female |  |
| *Pygathrix* | *nemaeus* |  | Colobinae | Presbytini | USNM 356577 | Male |  |
| *Pygathrix* | *nemaeus* |  | Colobinae | Presbytini | KUPRI 12438 | Male | This study |
| *Pygathrix* | *nigripes* |  | Colobinae | Presbytini | USNM 257998 | Female | https://www.morphosource.org |
| *Rhinopithecus* | *roxellana* |  | Colobinae | Presbytini | USNM 26887 | Male |  |
| *Rhinopithecus* | *roxellana* |  | Colobinae | Presbytini | USNM 26888 | Male |  |
| *Semnopithecus* | *entellus* |  | Colobinae | Presbytini | AMNH A992285 | Male |  |
| *Semnopithecus* | *entellus* |  | Colobinae | Presbytini | AMNH A992286 | Male |  |
| *Semnopithecus* | *entellus* |  | Colobinae | Presbytini | BSZM AP 187 | Male | This study |
| *Semnopithecus* | *entellus* |  | Colobinae | Presbytini | USNM 269058 | ? | https://www.morphosource.org |
| *Semnopithecus* | *johnii* |  | Colobinae | Presbytini | USNM 520675 | Female |  |
| *Semnopithecus* | *johnii* |  | Colobinae | Presbytini | USNM A49701 | Male |  |
| *Semnopithecus* | *thersites* |  | Colobinae | Presbytini | BSZM 19051056 | Female | This study |
| *Semnopithecus* | *thersites* |  | Colobinae | Presbytini | BSZM 19051058 | Male |  |
| *Semnopithecus* | *thersites* |  | Colobinae | Presbytini | BSZM 2161 | Male |  |
| *Semnopithecus* | *priam* |  | Colobinae | Presbytini | USNM 122634 | Female | https://www.morphosource.org |
| *Semnopithecus* | *priam* |  | Colobinae | Presbytini | BSZM 190614 | Female | This study |
| *Semnopithecus* | *priam* |  | Colobinae | Presbytini | USNM 196986 | Male | https://www.morphosource.org |
| *Semnopithecus* | *schistaceus* |  | Colobinae | Presbytini | USNM 21843 | Female |  |
| *Semnopithecus* | *schistaceus* |  | Colobinae | Presbytini | USNM 290066 | Male |  |
| *Simias* | *concolor* |  | Colobinae | Presbytini | KUPRI 1218 | Female | This study |
| *Simias* | *concolor* |  | Colobinae | Presbytini | KUPRI 1246 | Female |  |
| *Simias* | *concolor* |  | Colobinae | Presbytini | KUPRI 1248 | Female |  |
| *Simias* | *concolor* |  | Colobinae | Presbytini | KUPRI 1270 | Female |  |
| *Simias* | *concolor* |  | Colobinae | Presbytini | KUPRI 1284 | Female |  |
| *Simias* | *concolor* |  | Colobinae | Presbytini | KUPRI 1294 | Female |  |
| *Simias* | *concolor* |  | Colobinae | Presbytini | KUPRI 1301 | Female |  |
| *Simias* | *concolor* |  | Colobinae | Presbytini | KUPRI 1304 | Female |  |
| *Simias* | *concolor* |  | Colobinae | Presbytini | KUPRI 1311 | Female |  |
| *Simias* | *concolor* |  | Colobinae | Presbytini | KUPRI 1312 | Female |  |
| *Simias* | *concolor* |  | Colobinae | Presbytini | KUPRI 1221 | Male |  |
| *Simias* | *concolor* |  | Colobinae | Presbytini | KUPRI 1255 | Male |  |
| *Simias* | *concolor* |  | Colobinae | Presbytini | KUPRI 1269 | Male |  |
| *Simias* | *concolor* |  | Colobinae | Presbytini | KUPRI 1271 | Male |  |
| *Simias* | *concolor* |  | Colobinae | Presbytini | KUPRI 1277 | Male |  |
| *Simias* | *concolor* |  | Colobinae | Presbytini | KUPRI 1283 | Male |  |
| *Simias* | *concolor* |  | Colobinae | Presbytini | KUPRI 1295 | Male |  |
| *Simias* | *concolor* |  | Colobinae | Presbytini | KUPRI 1302 | Male |  |
| *Simias* | *concolor* |  | Colobinae | Presbytini | KUPRI 1307 | Male |  |
| *Simias* | *concolor* |  | Colobinae | Presbytini | KUPRI 1309 | Male |  |
| *Simias* | *concolor* |  | Colobinae | Presbytini | AMNH 103369 | Male | https://www.morphosource.org |
| *Simias* | *concolor* |  | Colobinae | Presbytini | AMNH 103371 | Female |  |
| *Simias* | *concolor* |  | Colobinae | Presbytini | USNM 121654 | Female |  |
| *Simias* | *concolor* |  | Colobinae | Presbytini | USNM 121659 | Male |  |
| *Simias* | *concolor* |  | Colobinae | Presbytini | USNM 121663 | Male |  |
| *Theropithecus* | *gelada* |  | Cercopithecinae | Papionina | AMNH 60568 | Male |  |
| *Theropithecus* | *gelada* |  | Cercopithecinae | Papionina | AMNH 80126 | Male |  |
| *Theropithecus* | *gelada* |  | Cercopithecinae | Papionina | FMNH 8174 | Male |  |
| *Theropithecus* | *gelada* |  | Cercopithecinae | Papionina | FMNH 27039 | Male |  |
| *Theropithecus* | *gelada* |  | Cercopithecinae | Papionina | FMNH 27040 | Male |  |
| *Theropithecus* | *gelada* |  | Cercopithecinae | Papionina | FMNH 27184 | Male |  |
| *Theropithecus* | *gelada* |  | Cercopithecinae | Papionina | FMNH 27187 | Male |  |
| *Theropithecus* | *gelada* |  | Cercopithecinae | Papionina | FMNH 27233 | Male |  |
| *Theropithecus* | *gelada* |  | Cercopithecinae | Papionina | FMNH 27234 | Female |  |
| *Theropithecus* | *gelada* |  | Cercopithecinae | Papionina | FMNH 35077 | ? |  |
| *Theropithecus* | *gelada* |  | Cercopithecinae | Papionina | FMNH 47767 | Female |  |
| *Theropithecus* | *gelada* |  | Cercopithecinae | Papionina | FMNH 57583 | ? |  |
| *Theropithecus* | *gelada* |  | Cercopithecinae | Papionina | FMNH 354990 | Female |  |
| *Theropithecus* | *gelada* |  | Cercopithecinae | Papionina | USNM 240885 | Male |  |
| *Theropithecus* | *gelada* |  | Cercopithecinae | Papionina | USNM 305107 | Male |  |
| *Trachypithecus* | *cristatus* |  | Colobinae | Presbytini | MCZ 35567 | Female |  |
| *Trachypithecus* | *cristatus* |  | Colobinae | Presbytini | MCZ 35584 | Female |  |
| *Trachypithecus* | *cristatus* |  | Colobinae | Presbytini | MCZ 35597 | Female |  |
| *Trachypithecus* | *cristatus* |  | Colobinae | Presbytini | MCZ 35603 | Female |  |
| *Trachypithecus* | *cristatus* |  | Colobinae | Presbytini | MCZ 35604 | Female |  |
| *Trachypithecus* | *cristatus* |  | Colobinae | Presbytini | USNM 113170 | Male |  |
| *Trachypithecus* | *cristatus* |  | Colobinae | Presbytini | USNM 113174 | Female |  |
| *Trachypithecus* | *cristatus* |  | Colobinae | Presbytini | USNM 198294 | Female |  |
| *Trachypithecus* | *cristatus* |  | Colobinae | Presbytini | USNM 198830 | Male |  |
| *Trachypithecus* | *cristatus* |  | Colobinae | Presbytini | KUPRI 2309 | Female | This study |
| *Trachypithecus* | *cristatus* |  | Colobinae | Presbytini | KUPRI 9478 | Female |  |
| *Trachypithecus* | *cristatus* |  | Colobinae | Presbytini | KUPRI 9492 | Female |  |
| *Trachypithecus* | *cristatus* |  | Colobinae | Presbytini | KUPRI 9494 | Female |  |
| *Trachypithecus* | *ebenus* |  | Colobinae | Presbytini | USNM 240489 | Female | https://www.morphosource.org |
| *Trachypithecus* | *germaini* |  | Colobinae | Presbytini | USNM 236627 | Female |  |
| *Trachypithecus* | *germaini* |  | Colobinae | Presbytini | USNM 252277 | Male |  |
| *Trachypithecus* | *germaini* |  | Colobinae | Presbytini | USNM 307717 | Male |  |
| *Trachypithecus* | *obscurus* |  | Colobinae | Presbytini | USNM 83259 | Female |  |
| *Trachypithecus* | *obscurus* |  | Colobinae | Presbytini | USNM 104446 | Female |  |
| *Trachypithecus* | *obscurus* |  | Colobinae | Presbytini | USNM 123993 | Male |  |
| *Trachypithecus* | *obscurus* |  | Colobinae | Presbytini | KUPRI 12131 | Male | This study |
| *Trachypithecus* | *françoisi* |  | Colobinae | Presbytini | KUPRI 12434 | Female |  |
| *Trachypithecus* | *françoisi* |  | Colobinae | Presbytini | KUPRI 12435 | Female |  |
| *Trachypithecus* | *françoisi* |  | Colobinae | Presbytini | KUPRI 12436 | Male |  |
| *Trachypithecus* | *phayrei* |  | Colobinae | Presbytini | USNM 307723 | Female | https://www.morphosource.org |
| *Trachypithecus* | *phayrei* |  | Colobinae | Presbytini | USNM 307737 | Male |  |
| *Trachypithecus* | *shortridgei* |  | Colobinae | Presbytini | USNM 277618 | Female |  |

SOM Table S3: List of the specimens included in this study.

SOM Table S4: Normality and homoscedasticity of the model residuals that test for the effect of taxonomy on morphometric ratios.

| Formula | Normality (Shapiro-Wilk) | Homoscedasticity (Bartlett) |
| --- | --- | --- |
| STT/ITT ~ Subfamily | W = 0.963  *p*-value < 0.001 | K^2^ = 15.73  *p*-value < 0.001 |
| STT/ITT ~ Tribes | W = 0.952  *p*-value < 0.001 | K^2^ = 31.85  *p*-value < 0.001 |
| STT/ITT ~ Genera | W = 0.93  *p*-value < 0.001 | K^2^ = 226.5  *p*-value < 0.001 |
| RPAL ~ Subfamily | W = 0.983  *p*-value < 0.001 | K^2^ = 1.77  *p*-value = 0.183 |
| RPAL ~ Tribes | W = 0.976  *p*-value < 0.001 | K^2^ = 29.06  *p*-value < 0.001 |
| RPAL ~ Genera | W = 0.96  *p*-value < 0.001 | K^2^ = 124.3  *p*-value < 0.001 |
| SA ~ Subfamily | W = 0.997  *p*-value = 0.216 | K^2^ = 3.87  *p*-value = 0.049 |
| SA ~ Tribes | W = 0.997  *p*-value = 0.337 | K^2^ = 20.19  *p*-value < 0.001 |
| SA ~ Genera | W = 0.997  *p*-value = 0.59 | K^2^ = 48.4  *p*-value < 0.001 |

SOM Table S5: Normality and homoscedasticity of the model residuals that test for the effect of sex on morphometric ratios.

| Formula | Normality (Shapiro-Wilk) | Homoscedasticity (Bartlett) |
| --- | --- | --- |
| *Co. guereza* STT/ITT ~ Sex | W = 0.957  *p*-value = 0.602 | K^2^ = 3.31  *p*-value = 0.069 |
| *Co. polykomos* STT/ITT ~ Sex | W = 0.969  *p*-value = 0.093 | K^2^ = 0.81  *p*-value = 0.368 |
| *N. larvatus* STT/ITT ~ Sex | W = 0.963  *p*-value = 0.134 | K^2^ = 8.50  *p*-value < 0.01 |
| *Pi. badius* STT/ITT ~ Sex | W = 0.976  *p*-value = 0.743 | K^2^ = 2.88  *p*-value = 0.09 |
| *Pr. verus* STT/ITT ~ Sex | W = 0.935  *p*-value = 0.142 | K^2^ = 4.72  *p*-value = 0.03 |
| *Pre. bicolor* STT/ITT ~ Sex | W = 0.955  *p*-value = 0.455 | K^2^ = 0.268  *p*-value = 0.604 |
| *Ce. mitis* STT/ITT ~ Sex | W = 0.90  *p*-value = 0.005 | K^2^ = 2.767  *p*-value = 0.100 |
| *Ch. aethiops* STT/ITT ~ Sex | W = 0.98  *p*-value = 0.913 | K^2^ = 0.739  *p*-value = 0.39 |
| *M. fascicularis* STT/ITT ~ Sex | W = 0.97  *p*-value = 0.04 | K^2^ = 0.391  *p*-value = 0.53 |
| *M. fuscata* STT/ITT ~ Sex | W = 0.95  *p*-value = 0.249 | K^2^ = 0.003  *p*-value = 0.954 |
| *P. anubis* STT/ITT ~ Sex | W = 0.90  *p*-value = 0.06 | K^2^ = 3.583  *p*-value = 0.058 |
| *Co. guereza* RPAL ~ Sex | W = 0.90  *p*-value = 0.07 | K^2^ = 0.03  *p*-value = 0.868 |
| *Co. polykomos* RPAL ~ Sex | W = 0.98  *p*-value = 0.24 | K^2^ = 0.12  *p*-value = 0.73 |
| *N. larvatus* RPAL ~ Sex | W = 0.98  *p*-value = 0.63 | K^2^ = 1.39  *p*-value = 0.24 |
| *Pi. badius* RPAL ~ Sex | W = 0.95  *p*-value = 0.22 | K^2^ = 3.31  *p*-value = 0.07 |
| *Pr. verus* RPAL ~ Sex | W = 0.95  *p*-value = 0.26 | K^2^ = 0.58  *p*-value = 0.45 |
| *Pre. bicolor* RPAL ~ Sex | W = 0.95  *p*-value = 0.35 | K^2^ = 0.52  *p*-value = 0.47 |
| *Ce. mitis* RPAL ~ Sex | W = 0.96  *p*-value = 0.35 | K^2^ = 0.38  *p*-value = 0.54 |
| *Ch. aethiops* RPAL ~ Sex | W = 0.94  *p*-value = 0.30 | K^2^ = 1.69  *p*-value = 0.19 |
| *M. fascicularis* RPAL ~ Sex | W = 0.97  *p*-value = 0.15 | K^2^ = 0.73  *p*-value = 0.39 |
| *M. fuscata* RPAL ~ Sex | W = 0.96  *p*-value = 0.39 | K^2^ = 0.52  *p*-value = 0.47 |
| *P. anubis* RPAL ~ Sex | W = 0.97  *p*-value = 0.53 | K^2^ = 0.35  *p*-value = 0.55 |
| *Co. guereza* SA ~ Sex | W = 0.96  *p*-value = 0.75 | K^2^ = 0.02  *p*-value = 0.88 |
| *Co. polykomos* SA ~ Sex | W = 0.98  *p*-value = 0.27 | K^2^ = 1.37  *p*-value = 0.24 |
| *N. larvatus* SA ~ Sex | W = 0.96  *p*-value = 0.11 | K^2^ = 1.51  *p*-value = 0.22 |
| *Pi. badius* SA ~ Sex | W = 0.98  *p*-value = 0.77 | K^2^ = 0.10  *p*-value = 0.75 |
| *Pr. verus* SA ~ Sex | W = 0.95  *p*-value = 0.35 | K^2^ = 4.41  *p*-value = 0.03 |
| *Pre. bicolor* SA ~ Sex | W = 0.97  *p*-value = 0.72 | K^2^ = 3.92  *p*-value = 0.05 |
| *Ce. mitis* SA ~ Sex | W = 0.99  *p*-value = 0.96 | K^2^ = 0.93  *p*-value = 0.33 |
| *Ch. aethiops* SA ~ Sex | W = 0.96  *p*-value = 0.62 | K^2^ = 3.03  *p*-value = 0.08 |
| *M. fascicularis* SA ~ Sex | W = 0.98  *p*-value = 0.24 | K^2^ = 1.74  *p*-value = 0.19 |
| *M. fuscata* SA ~ Sex | W = 0.97  *p*-value = 0.49 | K^2^ = 0.17  *p*-value = 0.68 |
| *P. anubis* SA ~ Sex | W = 0.98  *p*-value = 0.61 | K^2^ = 1.79  *p*-value = 0.18 |

SOM Table S6: Normality and homoscedasticity of the model residuals that test for the effect of taxonomy on principal component scores.

| Formula | Normality (Shapiro-Wilk) | Homoscedasticity (Bartlett) |
| --- | --- | --- |
| PC1 ~ Subfamily | W = 0.10  *p*-value = 0.06 | K^2^ = 1.90  *p*-value = 0.17 |
| PC1 ~ Tribes | W = 0.99  *p*-value = 0.001 | K^2^ = 1.21  *p*-value = 0.27 |
| PC1 ~ Genera | W = 0.99  *p*-value < 0.001 | K^2^ = 54.41  *p*-value < 0.001 |
| PC2 ~ Subfamily | W = 0.99  *p*-value < 0.001 | K^2^ = 115.4  *p*-value < 0.001 |
| PC2 ~ Tribes | W = 0.99  *p*-value = 0.013 | K^2^ = 21.3  *p*-value < 0.001 |
| PC2 ~ Genera | W = 0.99  *p*-value = 0.008 | K^2^ = 60.5  *p*-value < 0.001 |

SOM Table S7: Mean body mass of male and female taxa included in this study.

| Genus | Species | Sex | log(BodyMass) | Body mass (g) | Body Mass (kg) | References |
| --- | --- | --- | --- | --- | --- | --- |
| *Allenopithecus* | *nigroviridis* | Female | 3,519 | 3300 | 3,3 | Gautier-Hion, 2013a; Hill, 1966 |
| *Allenopithecus* | *nigroviridis* | Male | 3,792 | 6200 | 6,2 |  |
| *Allochrocebus* | *lhoesti* | Female | 3,544 | 3500 | 3,5 | Colyn, 1994; Sarmiento, 2013 |
| *Allochrocebus* | *lhoesti* | Male | 3,785 | 6100 | 6,1 |  |
| *Allochrocebus* | *preussi* | Female | 3,544 | 3500 | 3,5 | Butynski et al., 2009; Butynski, 2013 |
| *Allochrocebus* | *preussi* | Male | 3,74 | 5500 | 5,5 |  |
| *Cercocebus* | *agilis* | Female | 3,633 | 4300 | 4,3 | Colyn, 1994; Shah, 2013 |
| *Cercocebus* | *agilis* | Male | 3,954 | 9000 | 9 |  |
| *Cercocebus* | *atys* | Male | 4,025 | 10600 | 10,6 | Oates et al., 1990; McGraw, 2013 |
| *Cercocebus* | *atys* | Female | 3,792 | 6200 | 6,2 |  |
| *Cercocebus* | *galeritus* | Female | 3,633 | 4300 | 4,3 | No data so body weight of *Cer. agilis* presented in Oates et al., 1990; McGraw, 2013 was used |
| *Cercocebus* | *galeritus* | Male | 3,954 | 9000 | 9 |  |
| *Cercocebus* | *torquatus* | Female | 3,763 | 5800 | 5,8 | Malbrandt and Maclatchy, 1949; Ehardt, 2013 |
| *Cercocebus* | *torquatus* | Male | 4,025 | 10600 | 10,6 |  |
| *Cercopithecus* | *albogularis* | Female | 3,556 | 3600 | 3,6 | Lawes et al., 2013 |
| *Cercopithecus* | *albogularis* | Male | 3,756 | 5700 | 5,7 |  |
| *Cercopithecus* | *ascanius* | Female | 3,477 | 3000 | 3 | Colyn, 1994; Cords and Sarmiento, 2013 |
| *Cercopithecus* | *ascanius* | Male | 3,568 | 3700 | 3,7 |  |
| *Cercopithecus* | *cephus* | Female | 3,447 | 2800 | 2,8 | Gautier-Hion et al., 1999; Gautier-Hion, 2013b |
| *Cercopithecus* | *mitis* | Female | 3,591 | 3900 | 3,9 | Colyn, 1994; Lawes et al., 2013 |
| *Cercopithecus* | *mitis* | Male | 3,763 | 5800 | 5,8 |  |
| *Cercopithecus* | *mona* | Female | 3,447 | 2800 | 2,8 | Glenn and Bensen, 1998; Glenn et al., 2013 |
| *Cercopithecus* | *mona* | Male | 3,672 | 4700 | 4,7 |  |
| *Cercopithecus* | *neglectus* | Male | 3,892 | 7800 | 7,8 | Napier, 1981; Gautier-Hion, 2013c |
| *Cercopithecus* | *nictitans* | Female | 3,613 | 4100 | 4,1 | Gautier-Hion et al., 1999; Gautier-Hion, 2013d |
| *Cercopithecus* | *nictitans* | Male | 3,826 | 6700 | 6,7 |  |
| *Cercopithecus* | *petaurista* | Female | 3,462 | 2900 | 2,9 | Oates et al., 1990; McGraw et al., 2013 |
| *Cercopithecus* | *petaurista* | Male | 3,602 | 4000 | 4 |  |
| *Chlorocebus* | *aethiops* | Female | 3,447 | 2800 | 2,8 | Butynski and Kingdon, 2013a |
| *Chlorocebus* | *aethiops* | Male | 3,623 | 4200 | 4,2 |  |
| *Colobus* | *angolensis* | Female | 3,851 | 7100 | 7,1 | Bocian and Anderson, 2013 |
| *Colobus* | *angolensis* | Male | 3,949 | 8900 | 8,9 |  |
| *Colobus* | *caudatus* | Female | 3,869 | 7400 | 7,4 | No data so body weight of *Co. g. occidenalis* presented in Delson et al., 2000 was used |
| *Colobus* | *guereza* | Female | 3,869 | 7400 | 7,4 | Delson et al., 2000 |
| *Colobus* | *guereza* | Male | 3,968 | 9300 | 9,3 |  |
| *Colobus* | *polykomos* | Female | 3,919 | 8300 | 8,3 | O'Leary, 2003; Korstjens and Galat-Luong, 2013 |
| *Colobus* | *polykomos* | Male | 3,996 | 9900 | 9,9 |  |
| *Colobus* | *satanas* | Male | 4,045 | 11100 | 11,1 | Fleury and Brugière, 2013 |
| *Colobus* | *vellerosus* | Female | 3,839 | 6900 | 6,9 | Oates et al., 1994; Saj and Sicotte, 2013 |
| *Erythrocebus* | *patas* | Female | 4,146 | 14000 | 14,0 | Galat-Luong et al., 1996; Isbell, 2013 |
| *Erythrocebus* | *patas* | Male | 3,954 | 9000 | 9,0 |  |
| *Lophocebus* | *albigena* | Female | 3,806 | 6400 | 6,4 | Gautier-Hion and Gautier, 1976; Olupot and Waser, 2013 |
| *Lophocebus* | *albigena* | Male | 3,954 | 9000 | 9 |  |
| *Lophocebus* | *atterimus* | Female | 3,748 | 5600 | 5,6 | Colyn, 1994; Gautier-Hion, 2013e |
| *Lophocebus* | *atterimus* | Male | 3,898 | 7900 | 7,9 |  |
| *Lophocebus* | *johnstoni* | Female | 3,785 | 6100 | 6,1 | Olupot and Waser, 2013 |
| *Lophocebus* | *johnstoni* | Male | 3,914 | 8200 | 8,2 |  |
| *Macaca* | *arctoides* | Female | 3,924 | 8400 | 8,4 | Fooden, 1990; Smith and Jungers,1997 |
| *Macaca* | *arctoides* | Male | 4,086 | 12200 | 12,2 |  |
| *Macaca* | *assamensis* | Female | 3,833 | 6800 | 6,8 | Delson et al., 2000 |
| *Macaca* | *assamensis* | Male | 4,053 | 11300 | 11,3 |  |
| *Macaca* | *brunnescens* | Female | 3,954 | 9000 | 9 | No data so body weight of *Ma. tonkeana* presented in Supriatna, 1991 and Smith and Jungers, 1997 was used |
| *Macaca* | *brunnescens* | Male | 4,22 | 16600 | 16,6 |  |
| *Macaca* | *cyclopis* | Female | 3,69 | 4900 | 4,9 | Rothenfluh, 1976; Smith and Jungers, 1997 |
| *Macaca* | *cyclopis* | Male | 3,778 | 6000 | 6 |  |
| *Macaca* | *fascicularis* | Female | 3,491 | 3100 | 3,1 | Delson et al., 2000 |
| *Macaca* | *fascicularis* | Male | 3,69 | 4900 | 4,9 |  |
| *Macaca* | *fuscata* | Female | 3,903 | 8000 | 8 | Kimura and Hanada, 1995; Smith and Jungers,1997 |
| *Macaca* | *fuscata* | Male | 4,041 | 11000 | 11 |  |
| *Macaca* | *leonina* | Female | 3,699 | 5000 | 5 | Delson et al., 2000 |
| *Macaca* | *leonina* | Male | 3,892 | 7800 | 7,8 |  |
| *Macaca* | *maura* | Female | 3,778 | 6000 | 6 | Supriatna, 1991; Smith and Jungers, 1997 |
| *Macaca* | *maura* | Male | 3,964 | 9200 | 9,2 |  |
| *Macaca* | *mulatta* | Female | 3,944 | 8800 | 8,8 | Schwartz and Kemnitz, 1992; Smith and Jungers,1997 |
| *Macaca* | *mulatta* | Male | 4,041 | 11000 | 11 |  |
| *Macaca* | *nemestrina* | Female | 3,806 | 6400 | 6,4 | Delson et al., 2000 |
| *Macaca* | *nemestrina* | Male | 4,049 | 11200 | 11,2 |  |
| *Macaca* | *nigra* | Female | 3,653 | 4500 | 4,5 | Delson et al., 2000 |
| *Macaca* | *radiata* | Female | 3,785 | 6100 | 6,1 | Fooden, 1981; Smith and Jungers,1997 |
| *Macaca* | *radiata* | Male | 3,949 | 8900 | 8,9 |  |
| *Macaca* | *sinica* | Female | 3,491 | 3100 | 3,1 | No data so body weight of *Ma. fascicularis* presented in Delson et al., 2000 was used |
| *Macaca* | *sinica* | Male | 3,771 | 5900 | 5,9 | Isler et al., 2008 |
| *Macaca* | *sylvanus* | Female | 3,996 | 9900 | 9,9 | Fooden, 2007; Fa, 2013 |
| *Macaca* | *sylvanus* | Male | 4,161 | 14500 | 14,5 |  |
| *Macaca* | *thibetana* | Female | 4,149 | 14100 | 14,1 | Delson et al., 2000 |
| *Macaca* | *thibetana* | Male | 4,248 | 17700 | 17,7 |  |
| *Macaca* | *tonkeana* | Female | 3,954 | 9000 | 9 | Supriatna, 1991; Smith and Jungers,1997 |
| *Macaca* | *tonkeana* | Male | 4,173 | 14900 | 14,9 |  |
| *Mandrillus* | *leucophaeus* | Male | 4,301 | 20000 | 20 | Butynski et al., 2009; Schaaf et al., 2013 |
| *Mandrillus* | *sphinx* | Male | 4,483 | 30400 | 30,4 | Abernethy and White, 2013 |
| *Miopithecus* | *ogouensis* | Female | 3,053 | 1130 | 1,13 | Gautier-Hion et al., 1999; Gautier-Hion, 2013f |
| *Miopithecus* | *ogouensis* | Male | 3,14 | 1380 | 1,38 |  |
| *Nasalis* | *larvatus* | Female | 3,991 | 9800 | 9,8 | Delson et al., 2000 |
| *Nasalis* | *larvatus* | Male | 4,29 | 19500 | 19,5 |  |
| *Papio* | *anubis* | Male | 4,358 | 22800 | 22,8 | Jolly et al., 1997; Palombit, 2013 |
| *Papio* | *anubis* | Female | 4,09 | 12300 | 12,3 |  |
| *Papio* | *cynocephalus* | Male | 4,358 | 25800 | 25,8 | Altmann et al., 1993; Altmann et al., 2013 |
| *Papio* | *hamadryas* | Male | 4,23 | 17000 | 17 | Phillips-Conroy and Jolly, 1981; Swedell, 2013 |
| *Papio* | *papio* | Female | 4,146 | 14000 | 14 | Galat-Luong and Galat, 2013 |
| *Papio* | *papio* | Male | 4,415 | 26000 | 26 |  |
| *Papio* | *ursinus* | Female | 4,179 | 15100 | 15,1 | Colinshaw, 2013 |
| *Papio* | *ursinus* | Male | 4,452 | 28300 | 28,3 |  |
| *Piliocolobus* | *badius* | Female | 3,892 | 7800 | 7,8 | Delson et al., 2000 |
| *Piliocolobus* | *badius* | Male | 3,924 | 8400 | 8,4 |  |
| *Piliocolobus* | *foai* | Female | 3,857 | 7200 | 7,2 | Delson et al., 2000 |
| *Piliocolobus* | *gordonorum* | Female | 3,851 | 7100 | 7,8 | No data so body weight of *Pi. badius* presented in Delson et al., 2000 was used |
| *Piliocolobus* | *kirkii* | Female | 3,845 | 7000 | 7 | Siex and Struhsaker, 2013 |
| *Piliocolobus* | *preussi* | Female | 3,863 | 7300 | 7,3 | Butynski and Kingdon, 2013 |
| *Piliocolobus* | *rufomitratus* | Male | 3,987 | 9700 | 9,7 | Delson et al., 2000 |
| *Piliocolobus* | *tephrosceles* | Male | 3,987 | 9700 | 9,7 | No data so body weight of *Pi. rufomitratus* presented in Delson et al., 2000 was used |
| *Presbytis* | *bicolor* | Female | 3,799 | 6300 | 6,3 | Strasser, 1992 |
| *Presbytis* | *bicolor* | Male | 3,813 | 6500 | 6,5 |  |
| *Presbytis* | *canicrus* | Female | 3,799 | 6300 | 6,3 | No data so body weight of *Pre. bicolor* presented in Strasser, 1992 was used |
| *Presbytis* | *natunae* | Male | 3,813 | 6500 | 6,5 | No data so body weight of *Pre. bicolor* presented in Strasser, 1992 was used |
| *Presbytis* | *natunae* | Female | 3,708 | 5100 | 5,1 | Isler et al., 2008 |
| *Presbytis* | *potenziani* | Female | 3,806 | 6400 | 6,4 | Delson et al., 2000 |
| *Presbytis* | *potenziani* | Male | 3,785 | 6100 | 6,1 |  |
| *Presbytis* | *rubicunda* | Male | 3,792 | 6200 | 6,2 | Jungers, 1985; Strasser,1992 |
| *Presbytis* | *rubicunda* | Female | 3,756 | 5700 | 5,7 |  |
| *Presbytis* | *siamensis* | Male | 3,732 | 5400 | 5,4 | Isler et al., 2008 |
| *Presbytis* | *siamensis* | Female | 3,806 | 6400 | 6,4 |  |
| *Presbytis* | *siberu* | Male | 3,732 | 5400 | 5,4 | Isler et al., 2008 |
| *Presbytis* | *siberu* | Female | 3,732 | 5400 | 5,4 |  |
| *Procolobus* | *verus* | Female | 3,633 | 4300 | 4,3 | Delson et al., 2000 |
| *Procolobus* | *verus* | Male | 3,672 | 4700 | 4,7 |  |
| *Pygathrix* | *nemaeus* | Female | 3,908 | 8100 | 8,1 | Delson et al., 2000 |
| *Pygathrix* | *nemaeus* | Male | 4,037 | 10900 | 10,9 |  |
| *Pygathrix* | *nigripes* | Female | 3,908 | 8100 | 8,1 | No data so body weight of *Py. nemaeus* presented in Delson et al., 2000 was used |
| *Pygathrix* | *nigripes* | Male | 4,037 | 10900 | 10,9 |  |
| *Rhinopithecus* | *roxellana* | Male | 4,265 | 18400 | 18,4 | Delson et al., 2000 |
| *Semnopithecus* | *entellus* | Male | 4,258 | 18100 | 18,1 | Isler et al., 2008 |
| *Semnopithecus* | *johnii* | Female | 4,049 | 11200 | 11,2 | Delson et al., 2000 |
| *Semnopithecus* | *johnii* | Male | 4,068 | 11700 | 11,7 |  |
| *Semnopithecus* | *priam* | Female | 3,82 | 6600 | 6,6 | Isler et al., 2008 |
| *Semnopithecus* | *priam* | Male | 4,021 | 10500 | 10,5 |  |
| *Semnopithecus* | *schistaceus* | Male | 4,326 | 21200 | 21,2 | Delson et al., 2000 |
| *Semnopithecus* | *schistaceus* | Female | 4,185 | 15300 | 15,3 |  |
| *Semnopithecus* | *thersites* | Female | 3,845 | 7000 | 7 | Delson et al., 2000 |
| *Semnopithecus* | *thersites* | Male | 4,057 | 11400 | 11,4 |  |
| *Simias* | *concolor* | Female | 3,833 | 6800 | 6,8 | Delson et al., 2000 |
| *Simias* | *concolor* | Male | 3,964 | 9200 | 9,2 |  |
| *Theropithecus* | *gelada* | Female | 4,068 | 11700 | 11,7 | Dechow, 1983; Bergman and Beehner, 2013 |
| *Theropithecus* | *gelada* | Male | 4,279 | 19000 | 19 |  |
| *Trachypithecus* | *cristatus* | Female | 3,771 | 5900 | 5,9 | Jungers, 1985; Strasser, 1992 |
| *Trachypithecus* | *cristatus* | Male | 3,839 | 6900 | 6,9 |  |
| *Trachypithecus* | *ebenus* | Female | 3,771 | 5900 | 5,9 | No data so body weight of *Tr. cristatus* presented in Jungers, 1985 and Strasser, 1992 was used |
| *Trachypithecus* | *françoisi* | Female | 3,978 | 9500 | 9,5 | Smith and Jungers, 1997 |
| *Trachypithecus* | *françoisi* | Male | 4,033 | 10800 | 10,8 |  |
| *Trachypithecus* | *germaini* | Female | 3,771 | 5900 | 5,9 | No data so body weight of *Tr. cristatus* presented in Jungers, 1985 and Strasser, 1992 was used |
| *Trachypithecus* | *germaini* | Male | 3,833 | 6800 | 6,8 | Isler et al., 2008 |
| *Trachypithecus* | *obscurus* | Female | 3,82 | 6600 | 6,6 | Napier, 1981,1985; Strasser, 1992 |
| *Trachypithecus* | *obscurus* | Male | 3,863 | 7300 | 7,3 |  |
| *Trachypithecus* | *phayrei* | Female | 3,857 | 7200 | 7,2 | Isler et al., 2008 |
| *Trachypithecus* | *phayrei* | Male | 3,892 | 7800 | 7,8 |  |
| *Trachypithecus* | *shortridgei* | Female | 3,978 | 9500 | 9,5 | Isler et al., 2008 |
